# Genomic epidemiology of European *Aspergillus fumigatus* causing COVID-19-associated pulmonary aspergillosis in Europe

**DOI:** 10.1101/2023.07.21.550109

**Authors:** B.C. Simmons, J. Rhodes, T.R. Rogers, A.F. Talento, A. Griffin, M Mansfield, D. Sheehan, A. Abdolrasouli, P.E. Verweij, T. Bosch, S. Schelenz, S. Hemmings, M.C. Fisher

## Abstract

The opportunistic fungus *Aspergillus fumigatus* has been found to cause coinfections in patients with severe SARS-CoV-2 virus infection, leading to COVID-19-associated pulmonary aspergillosis (CAPA). The CAPA all-cause mortality rate is approximately 50% and may be complicated by azole-resistance. Genomic epidemiology can help shed light on the genetics of *A. fumigatus* causing CAPA including the prevalence of alleles that are associated with azole-resistance. Here, a population genomic analysis of 21 CAPA isolates from four European countries is presented. The CAPA isolates were compared with *A. fumigatus* from a wider population of 167 non-CAPA clinical isolates and 73 environmental isolates. Bioinformatic analysis and antifungal susceptibility testing were performed to quantify resistance and identify possible genetically-encoded azole-resistant mechanisms. Phylogenetic analysis of the 21 CAPA isolates showed a lack of genetic distinction from the wider *A. fumigatus* population, with isolates distributed within two distinct clades (A and B), with the majority of the CAPA isolates in clade B (71.4%). The prevalence of phenotypic azole-resistance in CAPA was 14.3% (*n*=3/21); all three CAPA isolates contained a known resistance-associated *cyp51A* polymorphism. CAPA isolates are drawn from the wider *A. fumigatus* population rather than forming a unique genetic background showing that COVID-19 patients are susceptible to the entire *A. fumigatus* population. However, the relatively high prevalence of azole-resistance alleles that we document poses a threat to treatment success rates, warranting enhanced detection and surveillance of *A. fumigatus* genotypes in these patients. Furthermore, potential changes to antifungal first-line treatment guidelines may be needed to improve patient outcomes.

## Introduction

*Aspergillus fumigatus* is the predominant causative agent of invasive aspergillosis (IA) (1, 2). The estimated annual prevalence of IA is >250,000 cases globally with a mortality rate of 30-95% (3, 4). Normally, only those who are immunocompromised or immunosuppressed are highly susceptible to developing IA (2), however, there is a widening group of patients who are at risk of IA including those with severe influenza or COVID-19.

In 2019-2020, the novel virus severe acute respiratory syndrome coronavirus 2 (SARS-CoV-2), causing COVID-19 disease, spread across the globe. Opportunistic pathogens have been widely reported as causing secondary infections in COVID-19 patients with lung damage and a notable proportion of these are fungal coinfections (12.6%) (5). A large proportion of fungal coinfections in COVID-19 cases are caused by *Aspergillus* species, with the predominant causal agent being *A. fumigatus*, giving rise to COVID-19-associated pulmonary aspergillosis (CAPA) (6, 7). The European Confederation for Medical Mycology (ECMM) and the International Society for Human and Animal Mycology (ISHAM) developed a consensus definition for CAPA at the end of 2020, stating that CAPA is a form of IA that is “in temporal proximity to a preceding SARS-CoV-2 infection” (7). The gap between the SARS-CoV-2 positive test and diagnosis of IA should be less than 2 weeks to consider the delay in the onset of CAPA (7, 8). The global prevalence of CAPA is reported to be between 3.8 – 40%, with 15.1% of ICU-admitted COVID-19 patients fulfilling the ECMM definition of CAPA (6–10). Furthermore, CAPA is characterised by low survival, with mortality ranging from 44 – 75% (10–12). The MYCOVID cohort study showed that COVID-19 patients who received intensive care treatment and positive sputum cultures for *A. fumigatus* but could not be classified as CAPA had higher mortality (45.8%) than those who had negative *A. fumigatus* cultures (32.1%) (10). However, this was lower than the group who had CAPA (61.8%) (10). Therefore, so-called colonisation may not be as harmless as previously thought.

In recent years, isolates of *A. fumigatus* resistant to the antifungal class azoles have emerged. The prevalence of antifungal drug resistance has rapidly increased but has only recently been declared a public health issue (13, 14). In the environment, population genomic studies have determined that depending on the country, between 2.2 – 20% of isolates are azole-resistant and this can be as high as 95.2% in Vietnam (15–20). A cause for concern is the increase in the proportion of azole-resistant *A. fumigatus* (AR*Af*) identified in patients, as this is associated with treatment failure, increased mortality rates and a doubling of health care costs (13, 14). Over the past 30 years, the UK has observed AR*Af* prevalence in clinical isolates to increase from 0.43% to 2.2% (21). In the Netherlands, there was a doubling in prevalence in clinical isolates between the periods 2007 – 2009 (5.3%) and 2013 – 2018 (11.7%) (22, 23). In CAPA, the prevalence of azole-resistance has not yet been established, as the sample size of the studies has been too small. To date three of the five CAPA genomic epidemiology studies used the European Committee on Antimicrobial Susceptibility Testing (EUCAST) (24) methods to conduct susceptibility tests. Of these two have reported no resistant isolates (25, 26) while one German study reported 22.2% (n=6/27) (27) as resistant. Using the Clinical and Laboratory Standards Institute (CLSI) method, a second German study (*n*=4) did not find any resistant isolates (28). Finally, a Portuguese study reported a high prevalence of AR*Af* of 45.5% (n=5/11) (29), however, that study did not state the method of determining how the isolates were sampled or found to be resistant.

Genomic epidemiological methods have played a key role in delineating the genetic basis of AR*Af* (13, 30, 31). The primary locus, with a high-frequency of non-synonymous SNPs (nsSNPs), known to be involved in azole-resistance is the *cyp51A* gene encoding the 14α-sterol demethylase enzyme (13, 30). The most common amino acid substitution hotspots are G54, L98H, G138, M220, and G448 (Supp Info Table 1). Point mutations occur in isolates cultured from patients who have had long-term exposure to azole therapy (30). Additionally, several tandem repeats (TRs) in the gene promoter region of *cyp51A* lead to the overexpression of the gene and are commonly associated with point mutations in the *cyp51A* gene (Supp Info Table 1). Genotypes of *A. fumigatus* that contain TR-mediated resistance are more common in environmental isolates and isolates obtained from azole-naïve patients. However, there is increasing awareness that resistance mechanisms in AR*Af* are complex and involve multiple genes (Supp Info Table 1) (16, 30–36). In studies that identified AR*Af* in CAPA isolates, Kirchoff *et al* (27) discovered only one CAPA isolate that contained TR_34_/L98H and five polymorphisms of AR*Af* via a non-*cyp51A* route. In another study, Morais *et al* (29) did not conduct bioinformatic analyses to delineate the genetic basis of their resistant CAPA isolates.

Whilst the development of a clinical definition for CAPA has aided clinicians in its diagnosis and ensuring treatment is commenced in a timely manner to improve patient outcomes, important questions remain as to the genetic characteristics and identity of *A. fumigatus* causing CAPA and the prevalence of antifungal resistance (13, 25–28, 37). In *A. fumigatus*, genomic epidemiological methods have begun to unravel the genetics of environmental and clinical antifungal resistance and the potential mechanisms of dispersion (1, 13, 16, 36). Therefore, there is potential for these methods to be adopted to extend the knowledge of CAPA (1). To date, there are only five genomic epidemiologic studies of CAPA using multiple methods (25–29). The results of these analyses were inconclusive on the genetic and epidemiological relatedness of *A. fumigatus* causing CAPA. The current study is the largest transnational genomic epidemiological investigation to date of CAPA isolates to determine where the genotypes of CAPA isolates group in the wider *A. fumigatus* population. Secondly, the study aimed to identify the mechanisms and determine the frequency of azole-resistant polymorphisms in CAPA isolates.

## Methods

### CAPA definition

The definition of CAPA was based on the 2020 ECMM/ISHAM consensus criteria (7). In summary, patients must have a positive SARS-CoV-2 polymerase chain reaction (PCR), require intensive care for COVID-19 and have signs of invasive pulmonary aspergillosis (IPA) infection. Ideally, IPA is confirmed through histopathological or direct microscopic detection of fungal hyphae obtained by lung biopsy, therefore, showing signs of tissue invasion and/or damage (7, 38). However, samples from bronchoalveolar lavage, bronchial aspirate and tracheal aspirate were used as alternatives as lung biopsies could rarely be performed in CAPA patients (7). A positive SARS-CoV-2 PCR test must occur between two weeks prior to hospital admission and up to 96hrs after ICU admission. CAPA may be further categorised depending on the sensitivity of the diagnostic method used. That is CAPA can be either proven, probable or possible (7, 11). A fourth category was added in cases where *A. fumigatus* was isolated from the sputum of a patient with COVID-19 receiving intensive care, but there were no clear signs of invasive disease (10). Furthermore, there was no follow-up bronchoscopy to determine if there was an invasive disease. Therefore, for the first analysis, the patients were categorised as non-CAPA and only colonised with *A. fumigatus* (‘colonising’). In the second analysis, the colonising isolates were categorised as clinical non-CAPA isolates.

### Fungal Isolates

Twenty-one CAPA isolates from four European countries between 2020-2021 were included (Table 1). In this study, all CAPA isolates were recovered as per ‘standard of care’ based on the ECMM criteria (7). Four CAPA isolates (CAPA-A – D) were from four separate cases that originated from two Cologne hospitals, Germany (28, 39). Six CAPA isolates were from London, UK (C422-C425, C611, C612). A further two isolates originated from two hospitals in the Netherlands (C403, C408). A further nine isolates were from two hospitals in Dublin, Ireland (C434-C441, C444). Isolate C444, has previously been described in a case report (40).

**Table 1.** Characteristics of CAPA isolates of *A. fumigatus* used in this study and details of alignments.

|  |  |  |  |  |  |
| --- | --- | --- | --- | --- | --- |
| Isolate ID | Country | Sample Type | No. of aligned reads (Millions) | Mean depth of coverage (x) | Percentage of reference genome covered (%) |
| CAPA-A | Germany | CAPA - Possible | 35.8 | 146.7 | 98.4 |
| CAPA-B | Germany | CAPA - Probable | 40.1 | 166.7 | 98.5 |
| CAPA-C | Germany | CAPA - Probable | 37.5 | 157.4 | 98.1 |
| CAPA-D | Germany | CAPA - Possible | 34.2 | 147.5 | 97.3 |
| C403 | Netherlands | CAPA - Possible | 37.2 | 165.0 | 98.6 |
| C408 | Netherlands | CAPA - Possible | 28.5 | 127.7 | 98.3 |
| C422 | UK | CAPA - Probable | 11.8 | 45.5 | 96.2 |
| C423 | UK | CAPA - Probable | 15.6 | 56.2 | 96.5 |
| C424 | UK | CAPA - Probable | 11.2 | 45.1 | 98.6 |
| C425 | UK | CAPA - Probable | 11.0 | 43.0 | 97.2 |
| C434 | Ireland | CAPA - Probable | 10.6 | 41.9 | 96.6 |
| C435 | Ireland | CAPA - Probable | 10.3 | 41.4 | 97.7 |
| C436 | Ireland | CAPA - Probable | 9.8 | 39.3 | 98.2 |
| C437 | Ireland | CAPA - Probable | 9.6 | 38.9 | 97.4 |
| C438 | Ireland | CAPA - Probable | 10.5 | 42.6 | 97.7 |
| C439 | Ireland | CAPA - Probable | 11.4 | 45.1 | 98.3 |
| C440 | Ireland | CAPA - Probable | 7.8 | 27.6 | 97.9 |
| C441 | Ireland | CAPA - Probable | 9.2 | 36.3 | 97.9 |
| C443 | Ireland | CAPA - Probable | 9.7 | 39.2 | 97.5 |
| C444 | Ireland | CAPA - Probable | 11.0 | 43.2 | 97.9 |
| C611 | UK | CAPA - Probable | 11.0 | 43.7 | 97.6 |
| C612 | UK | CAPA - Possible | 11.0 | 44.0 | 97.5 |

In this study, the genetic relatedness of CAPA isolates was compared to, 1) *A. fumigatus* isolates from non-CAPA IA cases, and isolates colonising non-CAPA patients; 2) Non-CAPA clinical and environmental isolates from the wider *A. fumigatus* population. The first analysis involved comparing CAPA isolates to two other groups. The first group comprised twelve *A. fumigatus* isolates from eleven patients with different types of IA (C120, C137-C140, C143, C307, C323, C360, C372, C376, C442). This included four isolates from a patient with necrotising aspergillosis, one trauma patient, two allergic bronchopulmonary aspergillosis (ABPA) patients, two with ABPA and asthma, and one with chronic pulmonary aspergillosis (CPA), and two with IPA (Supp Info Table 2). The selection criteria for these IA isolates were based upon the European Organization for Research and Treatment of Cancer and the Mycoses Study Group definition for invasive disease (38), and from the UK and Ireland. The second group comprised eight *‘*colonising’ isolates seven from three Dutch Hospitals (C402, C404-C407, C409, C410) and one from a patient in Ireland (C443) (36, 41–43). The second analysis involved comparing 21 CAPA, 167 non-CAPA clinical and 73 environmental *A. fumigatus* isolates. In summary, 218 UK and Irish *A. fumigatus* isolates that had been analysed by Rhodes *et al.* (47) and 23 additional *A. fumigatus* isolates (C307, C323, C360, C372, C376, C402, C404-C407, C409, C410, C426-C433, C442, C443) (Supp Info Table 2). All 261 isolates can be found in the Microreact project(41) at https://microreact.org/project/tWCYrr2hvKQJU3ygWnSJdY-capa.

### Clinical characterisation of the CAPA isolates

Applying the ECMM/ISHAM criteria, sixteen isolates were from probable CAPA cases (C422-C425, C434-C441, C444, C611, CAPA-C and CAPA-D) and five were from possible CAPA cases (C403, C408, C612, CAPA-A and CAPA-B).

### Antifungal susceptibility testing

Azole-resistance was defined as when an isolate that contained a polymorphism that is known to be associated with phenotypic azole-resistance (e.g. TR_34_/L98H in *cyp51A* gene) was identified. Susceptibility tests were carried out to determine whether the isolate was phenotypically resistant to azoles.

Susceptibility to azole antifungal agents was carried out on all 21 of the CAPA isolates within this study. Briefly, isolates were inoculated in 25cm^3^ tissue culture flasks containing Sabouraud dextrose agar and culture for a minimum of 2 days at 45°C. Conidia were collected in sterilised 0.05% (v/v) Tween-80 (Sigma-Aldrich) by filtration through sterilised glass-wool. Spore concentrations were adjusted to a final standardised spore suspension of 0.09-0.12 McFarland using sterilised 0.05% (v/v) Tween-80 using a haemocytometer. Twenty microlitres of final standardised spore suspensions were inoculated into Tebucheck multi-well and VIPcheck^TM^ (Mediaproducts BV, NL) plates. Initial screening was conducted with Tebucheck to determine which is resistant and susceptible and then VIPcheck^TM^ to give more information on which medical azoles the isolates were resistant to. The Tebucheck multi-well plate consists of four wells comprising of drug-free (growth control well), 6 mg/L, 8 mg/L and 16 mg/L of tebuconazole (44). All Tebucheck multi-well plates were made in-house as per the protocol set out in Brackin *et al*. (44). The VIPcheck^TM^ multi-well plate consists of wells with 4 mg/L itraconazole, 2 mg/L voriconazole, 0.5 mg/L posaconazole, and a drug-free growth control well (45). Cultured plates were left at 45°C until there was complete growth in the control well (48-72hrs). Each strain was compared to a non-CAPA clinical isolate C154 which was EUCAST susceptible and confirmed by both Tebucheck and VIPcheck^TM^.

The scoring of Tebucheck plates is as follows: all plates with growth only in well 1 (drug-free) were scored 1; if there was growth also in well 2 the isolate was scored 2; if there was growth in wells 1, 2 and 3 the isolate was scored 3; and if there was growth in all 4 wells the isolate was scored 4 (44). Any isolate with a score greater than 1 was considered resistant to tebuconazole (44).

The scoring of VIPcheck^TM^ can be found in Buil *et al*. (45). Briefly, after 48hrs if there is uninhibited growth in any well which contained an azole, then the well would be scored 2 and the isolate would be resistant to that azole. If there was minimal growth then they would be scored 1 and therefore be partly resistant. If there was no growth in the azole-containing wells then the well would be scored a 0 and the isolate would be considered susceptible to the azole.

Antifungal susceptibility data from previously published research was included in this study for isolates from Rhodes *et al.* (36) (C1-C200, E9-E206 and U1-3), C444 by Mohamed *et al.* (40) and CAPA-A – D by Steenwyk *et al.* (28). For the isolates, C307, C323, C360, C372 and C376 susceptibility testing was done as per the protocol set out in Rhodes *et al.* (36) by Armstrong-James and Scourfield (42, 43) based upon the recent EUCAST methodology (24). The Dutch isolates (C403-C410) had susceptibility testing done using EUCAST methodology (24). Finally, two Irish isolates (C438 and C441) that were azole-resistant minimum inhibitory concentrations (MICs) were performed using CLSI broth dilution technique M38-A2 by the Mycology Reference Laboratory Bristol UK (46).

### Genomic DNA preparation and whole genome sequencing

Whole-genome libraries of the *A. fumigatus* isolates used in their analysis were prepared and sequenced by different research teams and at different times, thus different platforms were used. Extraction of genomic DNA from the Irish and UK CAPA isolates (*n*=15) and some additional Irish non-CAPA isolates (clinical, *n*=2, and environmental, *n*=8) was carried out at Imperial College London. Briefly, gDNA was extracted using the MasterPure^TM^ Complete DNA and RNA Purification Kit (Lucigen, USA), including an additional bead-beating step using a FastPrep-24^TM^. Extracted gDNA was purified using a DNeasy Blood and Tissue Kit (Qiagen, Germany), with the concentration measured using a Qubit fluorometer and dsDNA Broad Range Assay kit (ThermoFisher Scientific, UK). DNA purity was assessed using NanoDrop^TM^ spectrophotometry. Purified gDNA was stored at −20°C prior to the construction of gDNA libraries, normalisation and indexing (Earlham Institute, UK). Libraries were run on a NovaSeq 6000 SP v1.5 flow cell (Illumina) to generate 150 bp paired-end reads.

CAPA (C403, C408) and colonising (clinical non-CAPA) (C402, C404-C407, C409, C410) isolates from the Netherlands (*n*=9) were sequenced using NextSeq 550 sequencer (Illumina), and 218 UK and Irish isolates and the five IA isolates (C307, C323, C360, C372, C376) were all sequenced using a HiSeq2500 (Illumina) sequencer (36). All these generated 50-150bp paired-end reads. The four CAPA isolates obtained from Germany were sequenced using the MGISEQ2000 from single-stranded circular DNA (ssCir DNA) (28, 47).

### Bioinformatics analysis

Raw Illumina whole-genome sequence (WGS) reads were quality checked using FastQC v0.11.9 (Brabham institute) and subsequently aligned to the Af293 reference genome using the Burrow-Wheeler Aligner alignment tool with maximal exact methods algorithm (48, 49). The quality of the alignment was improved and converted to sorted BAM format using sequence alignment/map (SAM) tools v1.15. To minimise inaccurate identification of single-nucleotide polymorphisms (SNPs) due to PCR duplication error, duplicated reads were marked using Picard v2.18.7. SNPs were identified using ‘Haplotype Caller’ from the Genome Analysis Toolkit v4.0. To ensure high confidence calls, SNPs had to achieve at least 1 parameter, QD < 2.0, fisher strand > 60.0, mapping quality < 40.0, mapping quality rank sum test < −12.5, read positive rank sum test < −8.0, and SOR > 4.0. The above expressions have been rigorously tested and benchmarked(36).

SNPs were mapped to genes using vcf-annotator v0.5 (Broad Institute). Tandem repeat sequences within the promoter region of *cyp51A*, and non-synonymous SNPs (nsSNPs) in the genic region of *cyp51A* were summarised for all isolates. In addition, nsSNPs were identified in other genes reported to have a role in azole drug resistance: *mshA* (Afu3g09850), Afu7g01960, *hmg1* (Afu2g03700), *erg6* (Afu4g03630), *erg25* (Afu4g04820), *aarA* (alpha amino reductase-A, Afu4g11240), *C2H2* (Afu5g07960), ATP-binding cassette (ABC) drug transporter (Afu1g17440), *nct* (Negative cofactor-2 complex)-*A* (Afu2g14250) and *nctB* (Afu3g02340).

### *A. fumigatus* and MAT identification

Previously, 223 UK and Irish *A. fumigatus* isolates have been confirmed to be a member of the *A. fumigatus* species complex using molecular methods (36). In this study, the CAPA isolates were confirmed as *A. fumigatus* for congruency with Af293 using the nucleotide sequence of the calmodulin (*CaM*) gene from Af293 utilising the basic local alignment searching tool (BLAST, v2.13.0) from the National Centre of Biotechnology Information (NCBI) (50, 51). A CAPA isolate was identified as *A. fumigatus* if the top BLAST hit had a percentage identification of >99% (52).

The mating type of all 261 *A. fumigatus* isolates was identified using BLAST v2.13.0 (NCBI) using sequences from Pyrzak *et al.* (53).

### Phylogenetic and spatial analyses

Whole-genome SNP data were converted to the presence/absence of an SNP with respect to the reference. SNPs identified as low confidence during variant filtration were assigned as missing. In the first analysis, as there were fewer than 50 taxa, maximum likelihood (ML) phylogenies were constructed using the gamma model (GTRGAMMA) of rate heterogeneity and rapid bootstraps over 1000 replicates in RAxML v8.2.9 (Stematakis, 2006 RAxML-VI-HPC) (54). In the second analysis, as there were greater than 50 taxa, maximum likelihood (ML) phylogenies were constructed using the CAT approximation (GTRCAT) of rate heterogeneity and rapid bootstraps over 1000 replicates in RAxML v8.2.9 (Stematakis, 2006 RAxML-VI-HPC) (54). Phylogenies were annotated and visualised in ggtree and ggtreeExtra in R (v4.2.2).

Genetic similarity and population allocation were investigated *via* principal component analysis (PCA) and discriminant analysis of principal components (DAPC) based on whole genome SNP data in R (v4.2.2). DAPC was performed using adegenet (55). To determine the number of principal components (PCs) to retain, the *a*-score method in the adegenet package was used. As the loadings of the PCs themselves are generally uninformative, to gain insight into how each isolate contributes to the cluster’s composition compoplots (adegenet) were generated.

### Statistical analyses

Descriptive analysis of data was conducted in R v4.2.2 and Microsoft Excel. Chi (χ^2^)-test was performed to address the association between CAPA and azole-drug resistance. Significant value was considered for *P*-values <0.05. Yates continuity correction was used to calculate χ^2^ in cases when values for a variable were less than 20.

### Data availability

All raw reads have been submitted to the European Nucleotide Archive (ENA) under Project Accession no. PRJEB60964. Two hundred and twenty-one raw reads of clinical non-CAPA and environmental *A. fumigatus* isolates have also been deposited under project accession no. PRJEB27135. The raw short reads of four German CAPA isolates have previously been deposited to the NCBI’s GenBank database under BioProject accession no. PRJNA673120.

### Ethics

All clinical isolates were collected and analysed as part of standard of care and all isolates were anonymised. The investigation of cases from whom these isolates (C1-C200, E9-E206, U1-U3, four German CAPA isolates and C444) originated had ethics approval and data are publicly available (28, 36, 40). Patients whose isolates were C434-C441 were identified as having CAPA as part of an audit/service improvement that had prior approval of the Research and Innovation, and of the Clinical Service Director, Office, St. James’s Hospital, Dublin, Ireland.

## Results

### WGS of 21 CAPA isolates

Reference-guided methods were used to analyse the genetic diversity of twenty-one CAPA isolates from four European countries (Table 1). All the isolates were confirmed to be *A. fumigatus*, as they shared 100% similarity with *CaM* of the Af293 genome. All the sequenced genomes mapped >96.2% (mean 97.8%) to the reference genome Af293 (Table 1), with a mean coverage of 151x (128-167x) for German and Dutch CAPA isolates, and 42.1x (27.6-56.2x) for UK and Irish isolates, reflecting the different sequencing methods and platforms used. Normalised whole-genome depth of coverage confirmed an absence of aneuploidy events in CAPA isolates. However, deletions and duplications were observed: All the isolates observed had approximately 260 kilo-base pairs (Kbp) deletion in chromosome IV (Supp Info Figure 1). This phenomenon has been observed in previous WGS studies and the region of the rRNA repeat cluster may be included in this deletion (48, 56). Deletions were also found in chromosome I (region of >200Kb), VII (region of >300kb) and VIII (approx. 60Kb region) in 7 (24.1%), 14 (48.3%) and 8 (27.5%) isolates respectively (Supp Info Figure 1). These regions have not been identified as having any known importance in the context of azole-resistance. Multiple peaks were observed in all the chromosomes from each isolate, which relate to the presence of copy number variations (CNVs).

In the first phylogenetic analysis, the mean depth of coverage was 73.4x and the sequenced genomes mapped to the *de novo* reference genome Af293 by a mean of 97.8% (Supp Info Table 2).

For the second phylogenetic analysis, the mean depth of coverage was 44.9X and the sequenced genomes mapped to the *de novo* reference genome Af293 by a mean of 97.4% (Supp Info Table 2).

An even proportion of mating idiomorphs in the CAPA isolates was observed, with 52% (*n*=11/21) containing the *MAT1-1* gene and 48% (*n*=10/21) that contained the *MAT1-2* gene. This is consistent with what is expected in the wild and suggestive of a sexually recombining population(53, 57). Combining the CAPA isolates with the UK and Ireland *A. fumigatus* isolates there was a bias towards the *MAT1-2* idiomorph (60.9%), which was statistically significant (χ^2^- test, p-value=0.014, degrees of freedom, d.f.=2). This means that the population may be more likely to reproduce asexually(36, 53, 57). Although, there was no significant difference between source types (Supp Info Table 3).

On average CAPA isolates differed from each other by 26,706 SNPs. There was only 1 CAPA isolate pair (C439-C440) that was highly similar (<2,670.6 SNPs), however, these were obtained from different patients. When CAPA isolates were compared to IA and colonised isolates, they differed by 30,986 SNPs. In the second analysis, the isolates differed by 25,449 SNPs.

### Phylogenetic and spatial analysis shows CAPA isolates are highly related to non-CAPA clinical and environmental *A. fumigatus*

To test the phylogenetic relatedness of *A. fumigatus* cultured from CAPA patients with the existing collection of *A. fumigatus* from IA and colonised patients, phylogenetic analysis of twenty-one CAPA, twelve non-CAPA IA isolates and eight non-CAPA colonising isolates from four European countries was conducted (Supp Info Table 2). This analysis revealed two broadly divergent clades which are referred to as clade A and clade B as previously reported (Figure 1 and Supp Figure 2) (36, 58). All branches had greater than 75% bootstrap support unless stated. Of the CAPA isolates the majority (*n*=15, 71.4%) were in clade B, and the rest were in clade A (*n*=6, 28.6%) (Figure 1 and Supp Table 3). All AR*Af* isolates (C438, C441, C444) were situated in clade A. All the Dutch CAPA isolates were from clade B, whereas only 55.6% of the Irish CAPA isolates were from clade B. The UK and German CAPA isolates had 84.3% and 75% of isolates in Clade B, respectively. Of the IA isolates (C120, C137-C140, C143, C307, C323, C360, C372, C376, C442), eight (66.6%) were in clade A and four (33.3%) in clade B. The IA cohort contained ten AR*Af* isolates, which all contained *cyp51A* polymorphisms. The eight IA isolates in clade A were AR*Af* and two (C120 and C372) of the four IA isolates in clade B were AR*Af* isolates (Figure 1 and Supp Info Table 5). Finally, seven of the eight colonising isolates comprised of clade B and all the isolates where not AR*Af*.

**Figure 1:**
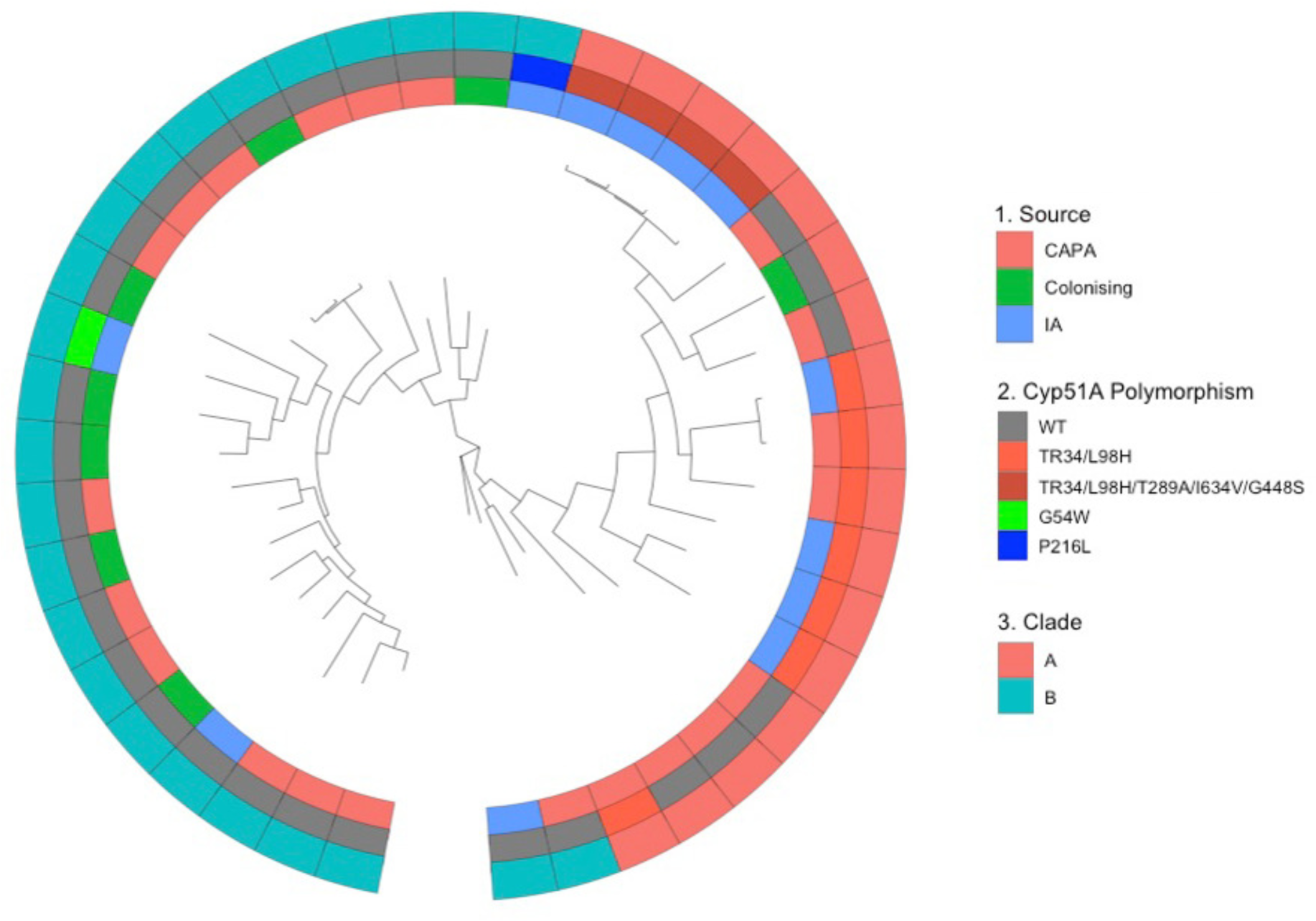
Phylogenetic tree of isolates from CAPA, IA and colonising patients. Unrooted maximum likelihood (ML) phylogenetic tree over 1000 replicates performed on WGS SNP data, showing: Inner track 1., the source of the isolate (CAPA, IA or colonising); Middle track 2., if the isolate contains *cyp51A* polymorphism; and Outer track 3., the clade the isolate is located.

Multivariate methods were used to identify and describe genetically related clusters. The principal component analysis (4 principal components, PCs) identified that CAPA isolates come from the same population as IA isolates as there was a high degree of overlap between CAPA and IA isolates (Figure 2A). Interestingly, the colonising isolates formed a subsection of the CAPA isolates (Figure 2A). Discriminant analysis of principal components (DAPC) and composition plot (5 PCs) confirmed this observation (Figures 2B&C). All the isolate genotypes contained a mixture of CAPA, IA and coloniser, with twenty-one of the isolates containing >20% CAPA genome (Figures 2B&C). Despite this, isolates labelled as IA or coloniser are not genetically distinct from CAPA isolates.

**Figure 2:**
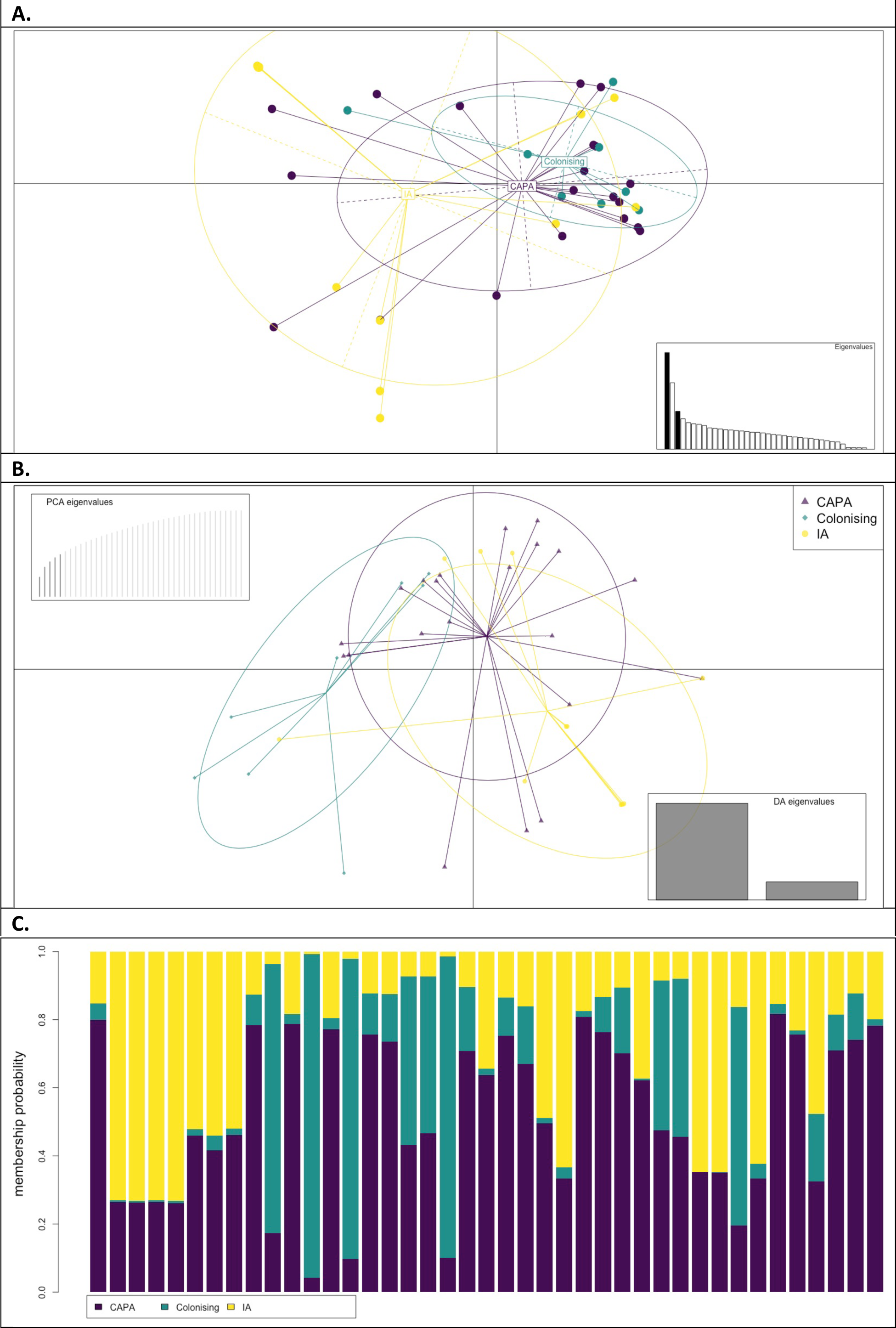
Multivariate analysis of isolates from CAPA, IA and colonising patients. **A**. Scatterplot of the principal component analysis (PCA) *A. fumigatus* genotypes using the first four principal components (PCs) illustrating the genetic identity of CAPA and IA control isolates. **B**. Density plot of the Discriminant PCA (DAPC), broadly identifying two distinct clusters, CAPA and control. **C**. Composition plot highlights that all the isolates’ genotype is composed of genetic material identified as CAPA, IA or colonising. Four isolates contain 70% control compared to CAPA, whereas there are 21 isolates whose genome membership is mostly CAPA (>80%).

To gain a more in-depth understanding of the genetic relatedness of CAPA isolates and whether they represent a distinct genetic identity or are drawn from the wider population of *A. fumigatus*, the phylogenetic relationship of CAPA isolates with the wider *A. fumigatus* population was investigated. Phylogenetic analysis was conducted on 240 clinical and environmental isolates and twenty-one CAPA isolates. All branches had greater than 75% bootstrap support unless stated (Supp Figure 4). Again, two broadly divergent clades were observed: clades A and B (Figure 3 and Supp Figures 3&4).

**Figure 3:**
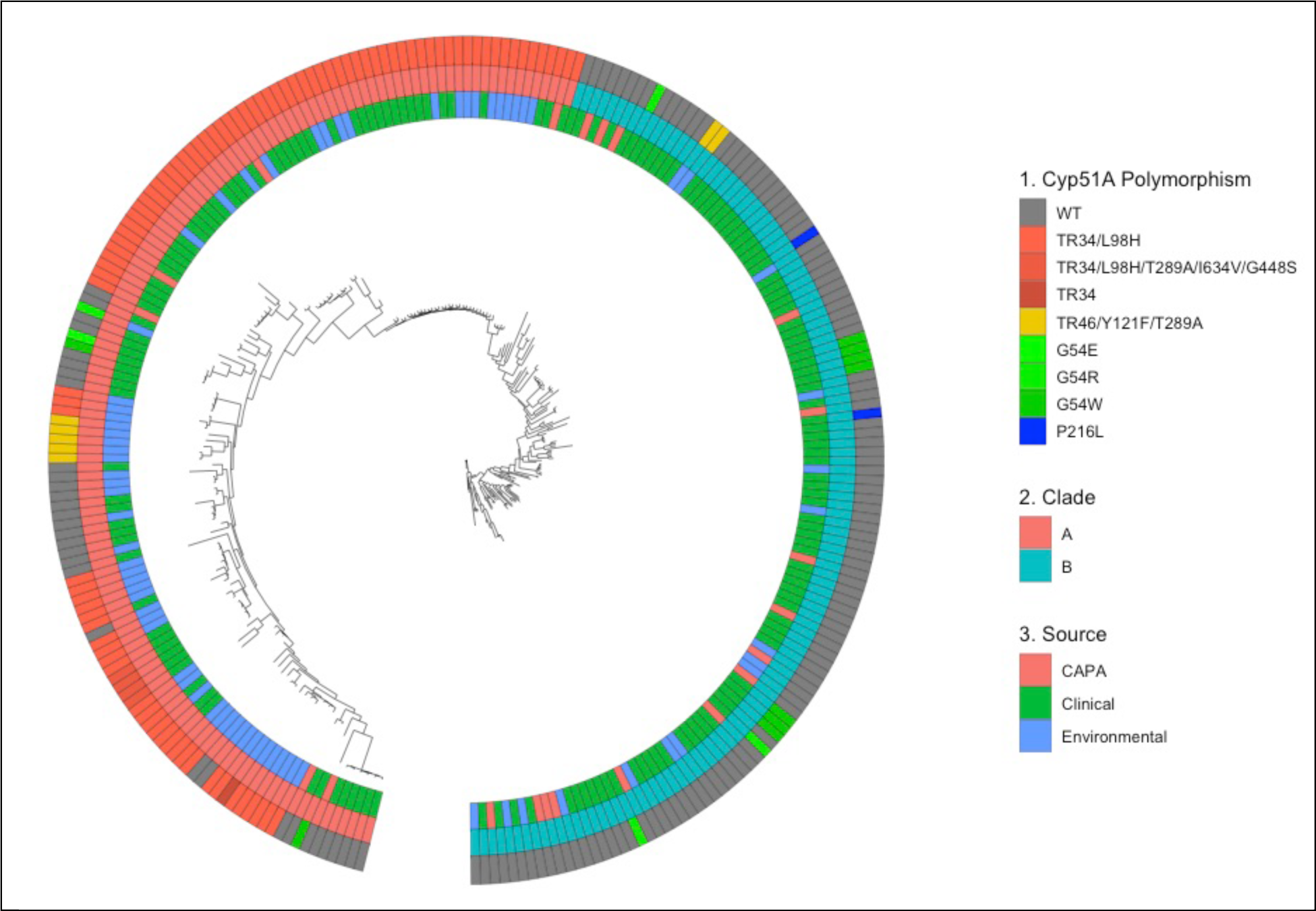
Phylogenetic tree of CAPA and *A. fumigatus* clinical and environmental isolates from Ireland and the UK. Unrooted ML phylogenetic tree over 1000 replicates performed on WGS SNP data, showing: Outer track 1., if the isolate contains *cyp51A* polymorphism; Middle track 2., the clade the isolate is located; and Inner track 3., the source of the isolate (CAPA, clinical or environmental).

In total 137 isolates (52.5%) lay within clade A and 124 isolates (47.5%) were situated in clade B. As seen in the first analysis, the CAPA isolates were largely associated with clade B (*n*=15, 71.4%) (Figure 3). The proportions of each isolate (CAPA, non-CAPA clinical and environmental) source type were associated with different clades, which were significantly different (χ^2^-test, *P*<0.001; d.f. = 2) (Supp Info Table 3). Additionally, isolates containing azole-resistant polymorphisms were significantly associated with clade A (χ^2^-test, *P*<0.001; d.f. = 1).

Multivariate analysis of the underlying genomic structure of the isolates confirmed that the genomes of CAPA isolates could not be distinguished from non-CAPA clinical and environmental *A. fumigatus* isolates originating from the UK and Ireland (Figure 4).

**Figure 4:**
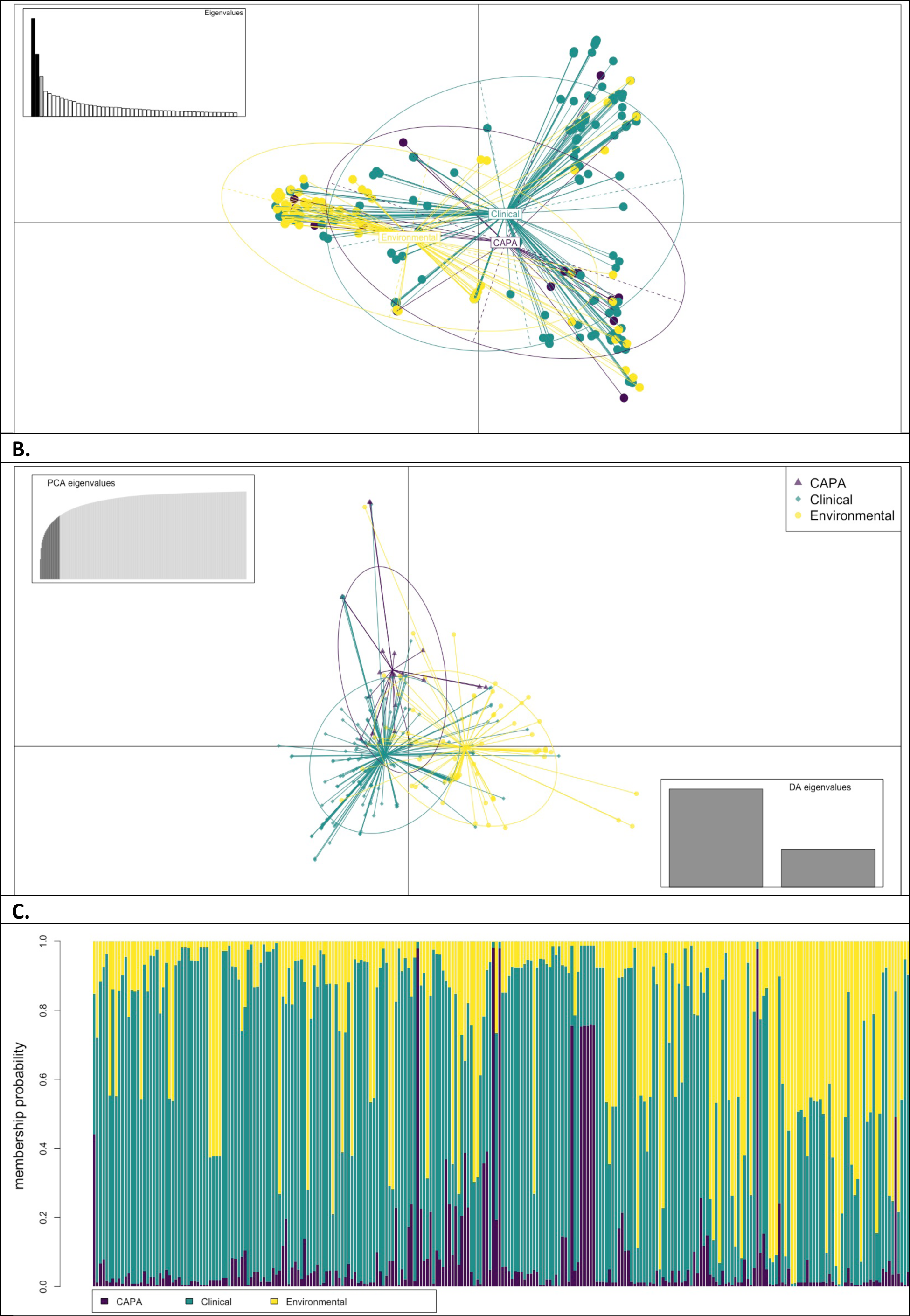
Multivariate analysis of CAPA and *A. fumigatus* clinical and environmental isolates from Ireland and the UK. Multivariate analysis, **A.** Scatterplot of the PCA *A. fumigatus* genotypes using the first five PCs illustrating the genetic identity of CAPA isolates, and clinical and environmental isolates from Ireland and the UK. **B.** Scatterplot of the DPCA identifying 3 types form separate clusters, with CAPA and clinical with some overlap. **C.** A composition plot comparing the genetic composition of each isolate. The plot highlights that each isolate is a mixture of genotypes identified as CAPA, environment and clinical non-CAPA.

### Azole-resistance within CAPA isolates primarily centred on known polymorphisms within *cyp51A*

Three CAPA (C438, C441, C444) isolates obtained from different locations in Ireland contained azole-resistant polymorphisms (TR_34_/L98H) (Table 3 and Supp Info Tables 4&5). No known drug-resistant polymorphisms were identified in the Dutch, UK or German CAPA isolates.

**Table 2.**
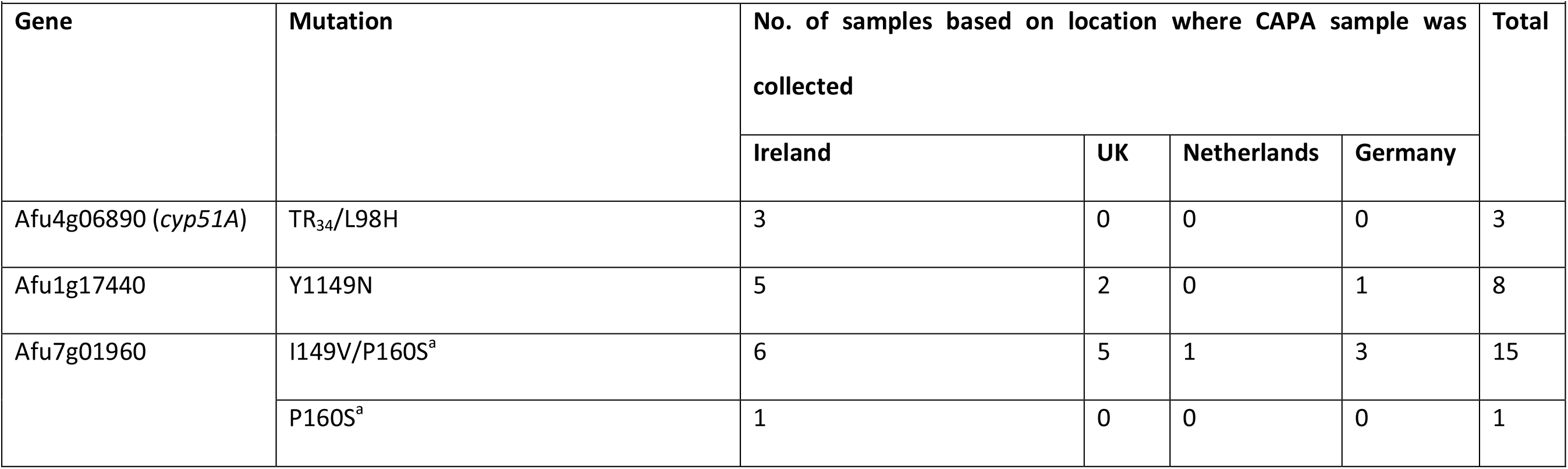
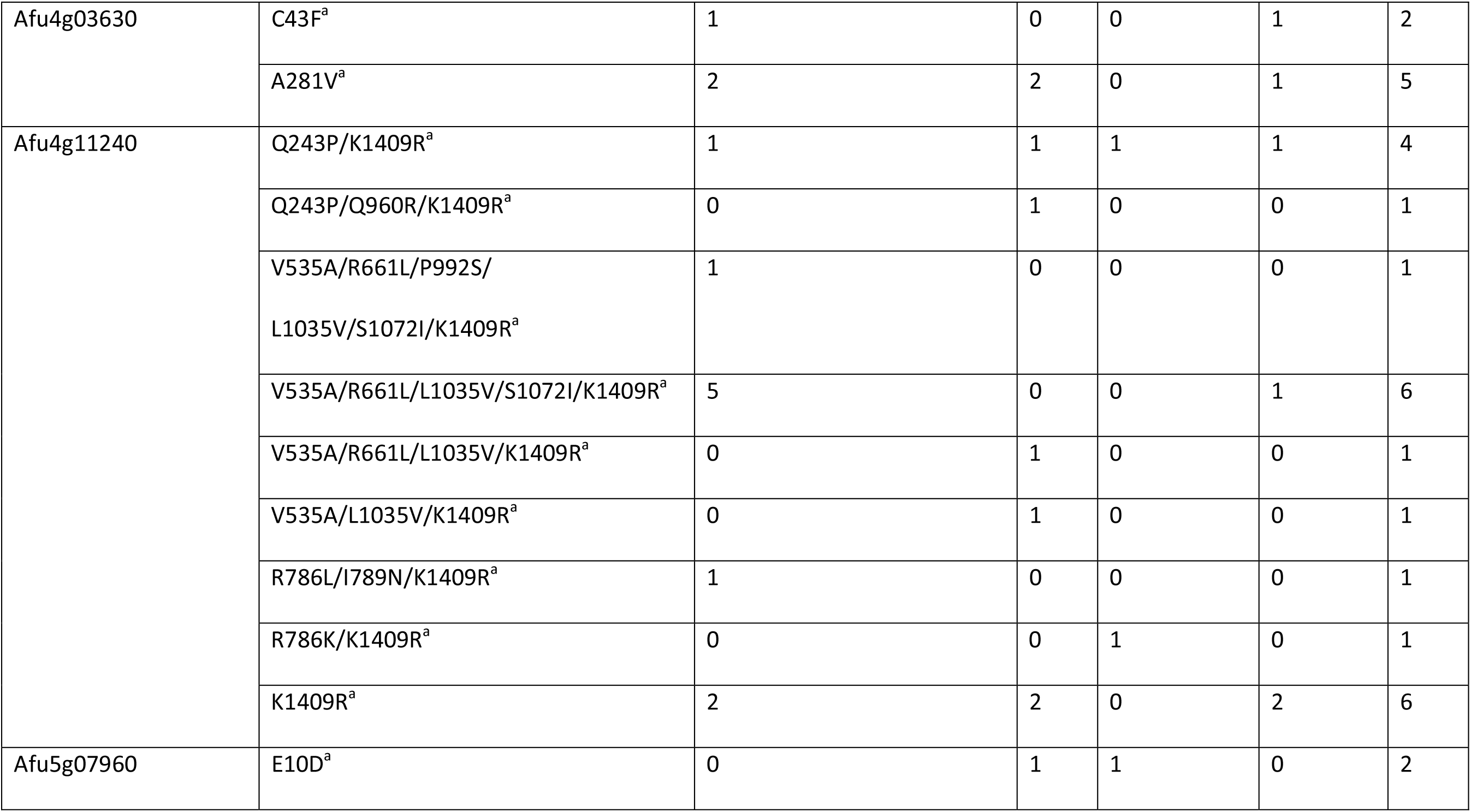
Number of Polymorphisms identified in the CAPA samples. X^a^ – Point mutations reported in this study, that have not previously been reported.

**Table 3.**
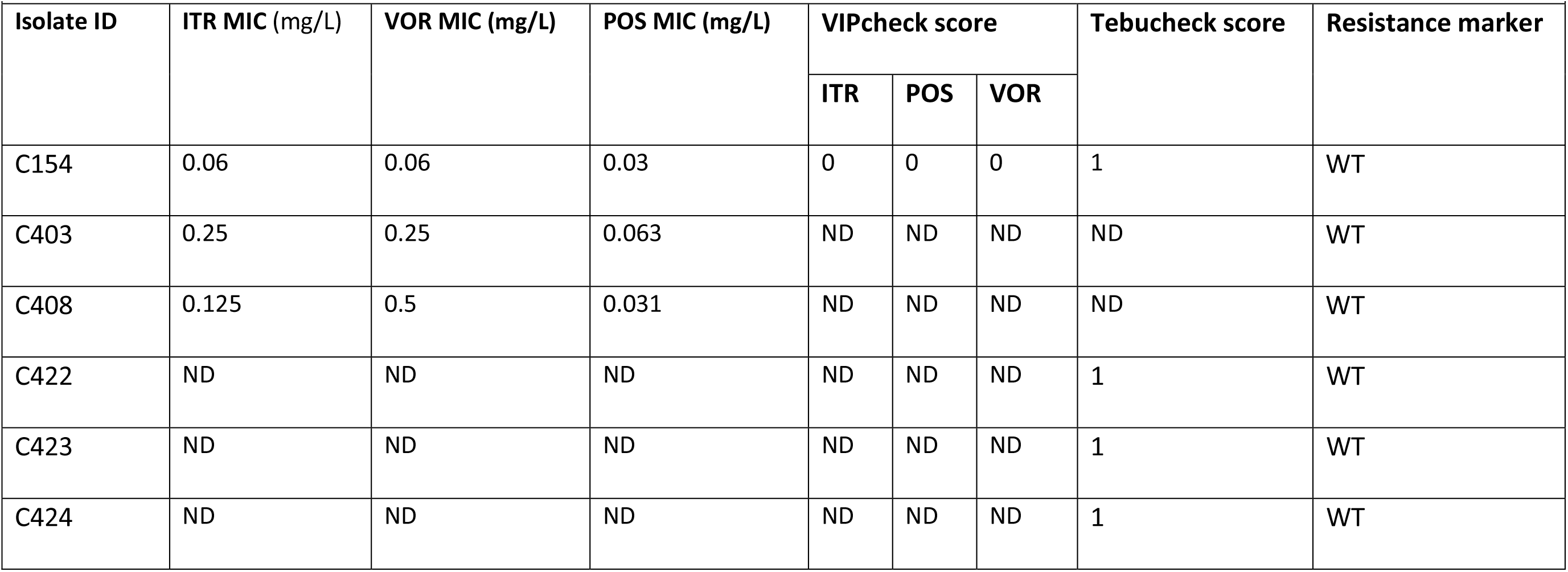
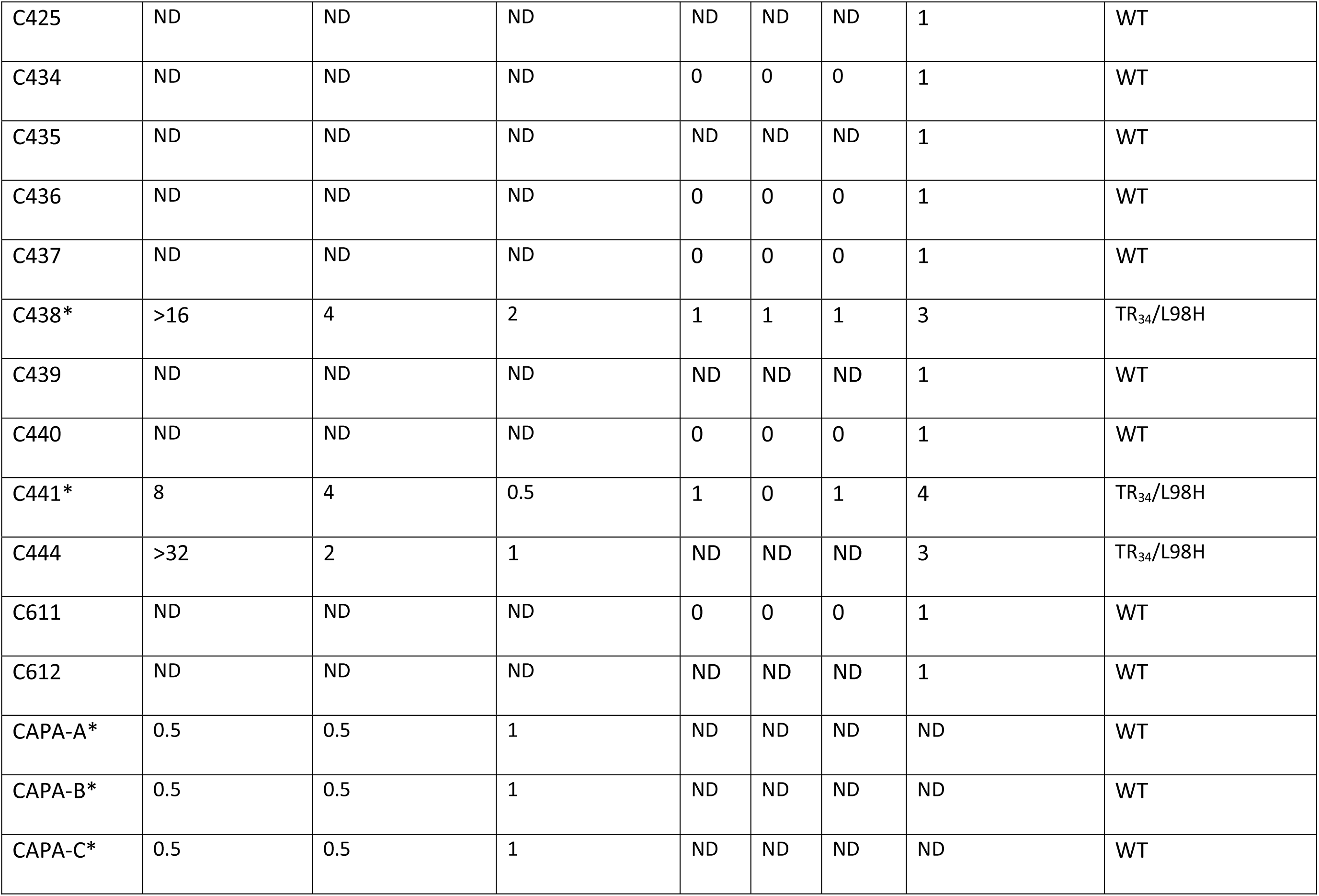
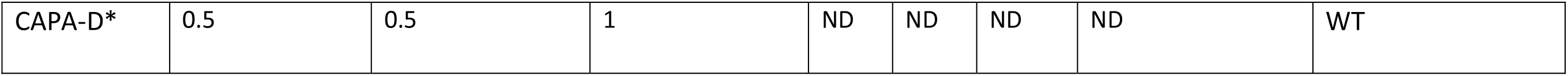
Azole drug susceptibility of Probable CAPA isolates grown in minimal medium and the associated candidate polymorphisms. C154 was used as a control as had previously been shown to be EUCAST susceptible(1) and in this study confirmed by Tebucheck and VIPcheck^TM^. ITR, Itraconazole (clinical breakpoint > 1 mg/L); VOR, Voriconazole (clinical breakpoint > 1 mg/L); POS, Posaconazole (clinical breakpoint > 0.25 mg/L)(2). VIPcheck score is 2 unihibited growth, 1 for minimal growth in well and 0 for no growth. The wells contain either ITR 4mg/L, POS 0.5 mg/L or VOR 2mg/L(3). Tebucheck score, 1 = fully susceptible to tebuconazole (0mg/L); 2 = resistant, growth in 6 mg/L of tebuconazole; 3 = resistant, growth in 8 mg/L of tebuconazole; 4 = resistant, growth in 16 mg/L of tebuconazole(4). C444 had clinical azole susceptibility testing done by Mohamed *et al*.(5). CAPA-A – D* were tested in the study by Steenwyk *et al.* using the Clinical and Laboratory Standards Institute (CLSI) method method(6).

Susceptibility testing was performed to confirm an isolate containing azole-resistant polymorphisms has raised MICs to azole drugs. Of the CAPA isolates three (C403, C408, C444) had standard EUCAST broth microdilution antifungal susceptibility testing performed (Table 3)(40). The German (CAPA-A – D) and two Irish (C438 and C441) isolates were tested using the CLSI method. All three isolates were triazole-resistant (Table 3). Sixteen of the CAPA isolates had Tebucheck performed, and of these 9 had VIPcheck^TM^ (Table 3). The results of the Tebucheck and VIPcheck^TM^ confirmed that only isolates C438, C441 and C444 were resistant to at least one azole (Table 3).

## Discussion

The COVID-19 pandemic has created a large and growing global cohort of patients at risk of developing *A. fumigatus* coinfections. Additionally, CAPA has higher mortality than COVID-19 itself (16-25% excess mortality rate) (7). This in part has been due to the challenges of determining whether COVID-19 patients have IA (7, 10). Therefore, delays in commencing prompt antifungal treatment may have led to higher mortality rates (59). Furthermore, WGS could address these issues by providing information on potential common sources, the genetic relatedness of the CAPA isolates with the wider *A. fumigatus* population and the presence of AR*Af*. Therefore, genomics has the potential to inform clinicians about the most effective way to treat CAPA patients (1, 13). In this study, WGS was carried out on twenty-one CAPA isolates from four European countries to explore the genomic epidemiology of *A. fumigatus* causing CAPA.

The present genomic analysis of CAPA isolates yielded three major findings. First, CAPA isolates were comprised of a diverse range of *A. fumigatus* genotypes. Secondly, CAPA genomes are an even mix of *A. fumigatus* genotypes found in previous studies in either the clinical environment or the environment. Finally, our study adds support to the growing body of evidence that the prevalence of AR*Af* is elevated in at-risk patient cohorts. However, an important caveat is that the only AR*Af* found were from one country likely due to the overall small sample size of *A. fumigatus* studied.

To gain insight into CAPA’s genetic and epidemiological relatedness, twenty-one CAPA isolates were compared to different types of *A. fumigatus* samples including clinical non-CAPA and environmental isolates. This analysis provided evidence that genotypes of *A. fumigatus* isolates from CAPA patients are genetically diverse, with two broadly divergent clades A and B. A proportion of CAPA isolates clustered in clade B (*n*=22, 78.6%). In the sample of IA isolates, 64% were in clade A. The phenomenon of the two-clade structure of *A. fumigatus* species was first identified in a large global genetic epidemiology study of over 4,000 *A. fumigatus* isolates and has since been replicated in different *A. fumigatus* populations (36, 58, 60).

Even though the genotypes of CAPA isolates in this study were biased towards clade B, the isolates were genetically diverse. The genetic relatedness between CAPA isolates is on average 26,076 SNPs. The genetic relatedness between isolates was higher when compared across twelve IA and eight colonising isolates (average 30,986 SNPs) and the wider *A. fumigatus* population (25,448 SNPs). This is comparable to what was previously reported (36, 60). In an earlier study, the genetic diversity was higher in UK isolates (27,914SNPs) compared to India (5,100 SNPs) (60). This result may have occurred as the 21 CAPA isolates were from four different European countries, whereas the previous studies reporting genetic distances were from smaller geographical distances (36, 60). However, in this study, the majority of non-CAPA isolates were from the UK and Ireland, so future work should sequence A. fumigatus from a wider geographical area. Only 1 pair of isolates was highly related (<2,607.6 SNPs; C439-C440) and originated from the same country, but from different patients. This could be because of patients being exposed to the same environmental source. However, no epidemiological information was available to investigate this possibility.

Secondly, using hypothesis-free population genetic methods we identified that CAPA isolates were a mixture of *A. fumigatus* genotypes. PCA and DAPC allow for the genetic structuring of populations to be better understood (61). In this study, the genotypes from the 21 CAPA isolates showed genetic overlapping with *A. fumigatus* genotypes isolated from patients with IA, and those obtained from other patients and the environment. In the five previous genomic epidemiological studies of CAPA, the genotypes from CAPA isolates were only compared with non-CAPA clinical isolates or COVID-19 patients who were colonised with *A. fumigatus* and had no active IA (25–29). The results of Steenwyk *et al.* suggest that overall CAPA genotypes are drawn from the wider clinical population of *A. fumigatus* isolates despite observing some genetic clustering (28). This has been replicated in a larger transnational genomic study (25). A larger multicentre study using microsatellite methods to genotype isolates identified that the genotypes obtained were genetically distinct from one another (27). This finding was replicated in one Spanish and one Portuguese study using tandem repeats within exons of surface proteins and *erg* coding genes (TRESPERG), and microsatellite methods, respectively, to compare isolates obtained from CAPA patients and the environment (26, 29). The authors identified that the CAPA isolates were from a diverse genetic pool (26, 27, 29). However, compared to WGS, TRESPERG and microsatellite methods are limited in the power to provide an in-depth analysis of the genetic structure and diversity within a population (62, 63). This is because, unlike WGS, TRESPERG and microsatellite methods genotype only a small proportion of the whole organism’s genome (63). Therefore, this study is the first to conduct an in-depth analysis to delineate the genetic diversity of *A. fumigatus*, in CAPA cases. Furthermore, to show that the genotypes of *A. fumigatus* obtained from patients with CAPA are a genetically diverse mixture of clinical and environmental *A. fumigatus* genotypes, WGS is best placed to investigate this (31, 62). Therefore, using genomic analysis alone, it is difficult to distinguish between the *A. fumigatus* genotypes that colonise patients’ airways and cause CAPA and IA. However, host factors could help explain the progression from colonisation to invasive disease in CAPA patients.

COVID-19 patients who develop CAPA may have acquired *A. fumigatus* isolates from the environment. This is supported by the evidence that the genotypes of CAPA isolates in this study are a mixture of genomes representative of both clinical and environmental *A. fumigatus* isolates. Previously, studies have identified that clinical non-CAPA *A. fumigatus* isolates are genetically similar to those sourced from the environment (26, 29, 36, 60). Additionally, AR*Af* has been isolated from several azole-naïve IA and CAPA cases (40, 64, 65) suggesting their resistance was pre-acquired from environmental (community and hospital) inocula. It has been hypothesised that patients who develop IA are first colonised by *A. fumigatus* isolates from airborne conidia in their environment. Thus, CAPA patients may acquire and become colonised with *A. fumigatus* on exposure to airborne *A. fumigatus* conidia in either the community or from the hospital environment. These conidia under favourable conditions (immunosuppression from SARs-CoV-2 and drugs used to treat COVID-19: such as dexamethasone and tazoluzimab), develop hyphae and subsequently invade the surrounding tissue leading to the onset of CAPA. If the patient is being exposed or treated with triazoles, it is at this stage that resistant inocula will remain in the patient’s airway.

Currently, the frequency of AR*Af* within the CAPA population is unknown. The present study used bioinformatic tools and drug susceptibility tests to identify three AR*Af* out of 21 CAPA isolates containing the predominant *cyp51A* polymorphism TR_34_/L98H and phenotypically resistant to at least one azole. Furthermore, all three were pan-azole-resistant (40). In the rest of the sample of CAPA isolates, none had known azole-resistant polymorphisms and were susceptible to azoles phenotypically. This study, therefore, represents the largest and most detailed findings on AR*Af* population genomics in CAPA from multiple countries to date. In a large Spanish tertiary hospital of 28 CAPA patients, AR*Af* prevalence was zero (26). This is despite AR*Af* being previously isolated from patients throughout Spain (66). Similarly, a recently published small transnational study (*n*=11) and a small German study (*n*=4) did not identify any AR*Af* in the sample of CAPA isolates (25, 28). In a German multicentre study, the prevalence of AR*Af* isolates was found to be 22.2% (*n*=6/27)(27). Only one of the six CAPA isolates was identified to have known *cyp51A* polymorphisms conferring resistance (27). However, the methodology lacked sufficient detail for it to be reproducible (26). A recent small Portuguese study (*n*=11) identified a high prevalence of 45.5% (29). However, that study did not specify how these were tested for azole-resistance and confirmed as AR*Af*. Furthermore, it is difficult to generalise the results due to the small sample size and the isolates were sampled from a single centre. In the Netherlands, a screening programme of CAPA patients identified one patient in a cohort of twenty-two to have an AR*Af* strain (4.5%) (67). Other studies identifying AR*Af* in CAPA patients were from case reports (40, 64, 68). In the wider patient population prevalence of AR*Af* has been increasing in the last 30 years from 0.43% to 2.2% in London (21). Different countries have recently reported varying levels, Germany at 3.5%, Denmark at 6.1%, Spain at 7.4%, Netherlands at 11.7% and Japan at 12.7% (17, 20, 22, 23, 66). In the UK, the prevalence of AR*Af* in the air, to which people are exposed, was recently found to range from 3-9% depending on the season (16, 19). Thus, consideration of AR*Af* surveillance in CAPA and wider patient populations who may be exposed to *Aspergillus* conidia in the hospital air is recommended owing to this near-ubiquitous exposure.

Future directions should focus on developing a global surveillance methodology for *A. fumigatus* for both clinical and environmental isolates, which is user-friendly and easily accessible. Additionally, surveillance programmes should include methods of monitoring antifungal resistance, owing to the high prevalence of resistance to frontline clinical azoles (e.g., itraconazole, voriconazole, isavuconazole and posaconazole). For example, each laboratory submits a random sample isolate to a national centre for antifungal susceptibility testing, either quarterly or annually depending on prevalence. Furthermore, an exposure assessment could be used to aid clinical decisions on the most effective course of antifungal treatment. Finally, a review of local antifungal treatment guidelines in response to the increasing prevalence of AR*Af* needs to be further considered.

## Acknowledgements

This research was supported by NERC (nos. NE/P001165/1 and NE/P000916/1), the UK Medical Research Council (MRC) (no. MR/R015600/1) and Wellcome Trust (no. 219551/Z/19/Z). MCF is supported by the CIFAR Fungal Kingdoms Program.

## Supplementary Information Tables

**Supp Info Table 1.**
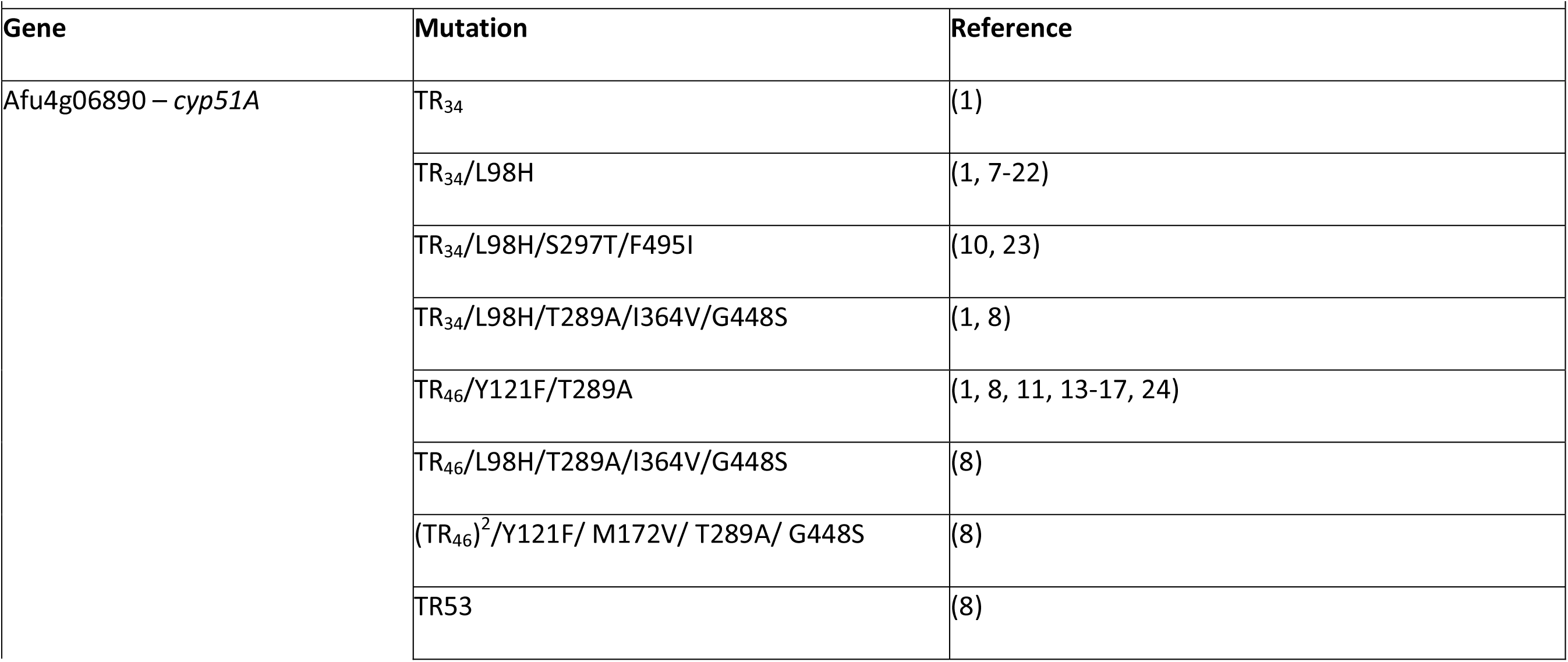
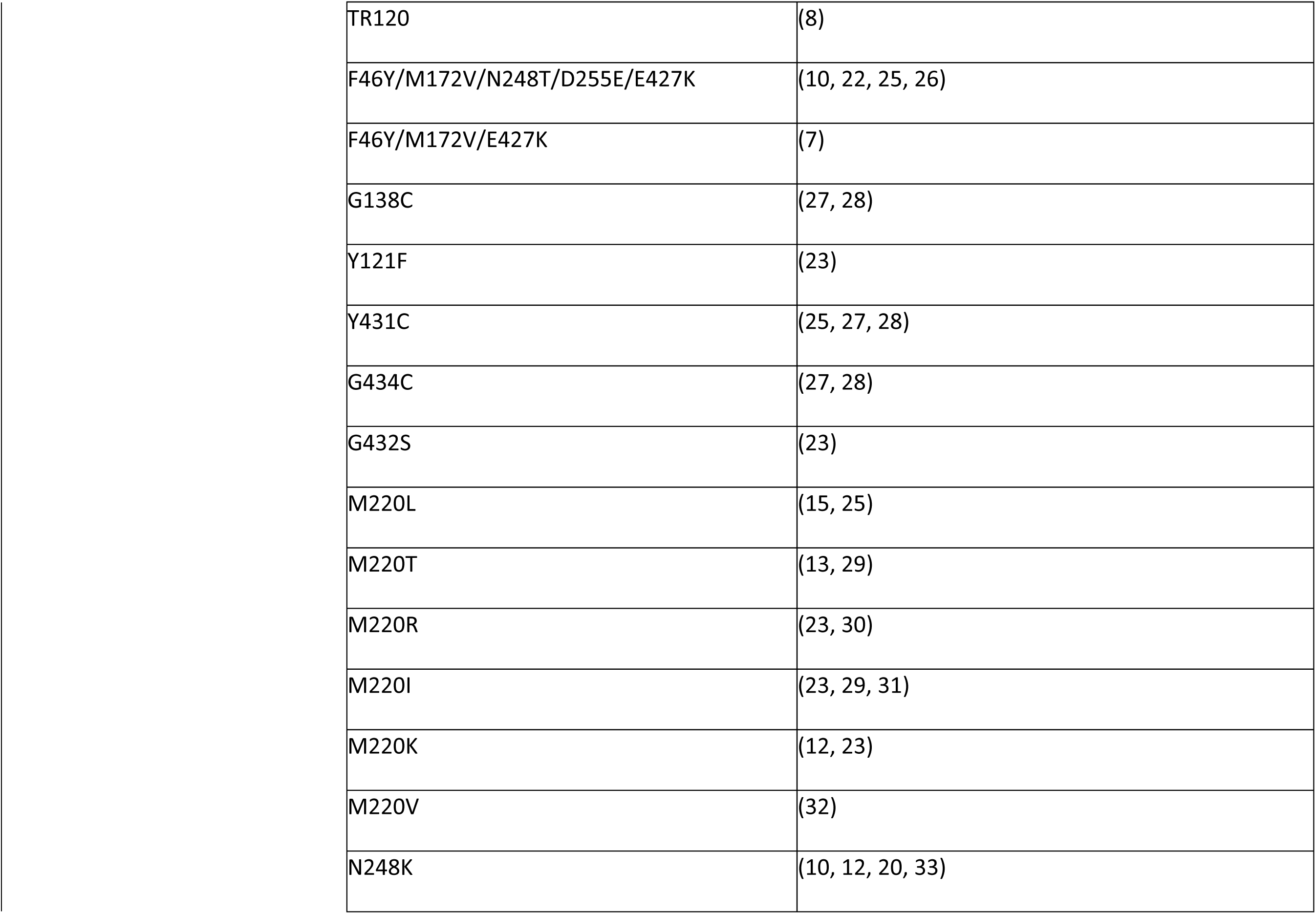
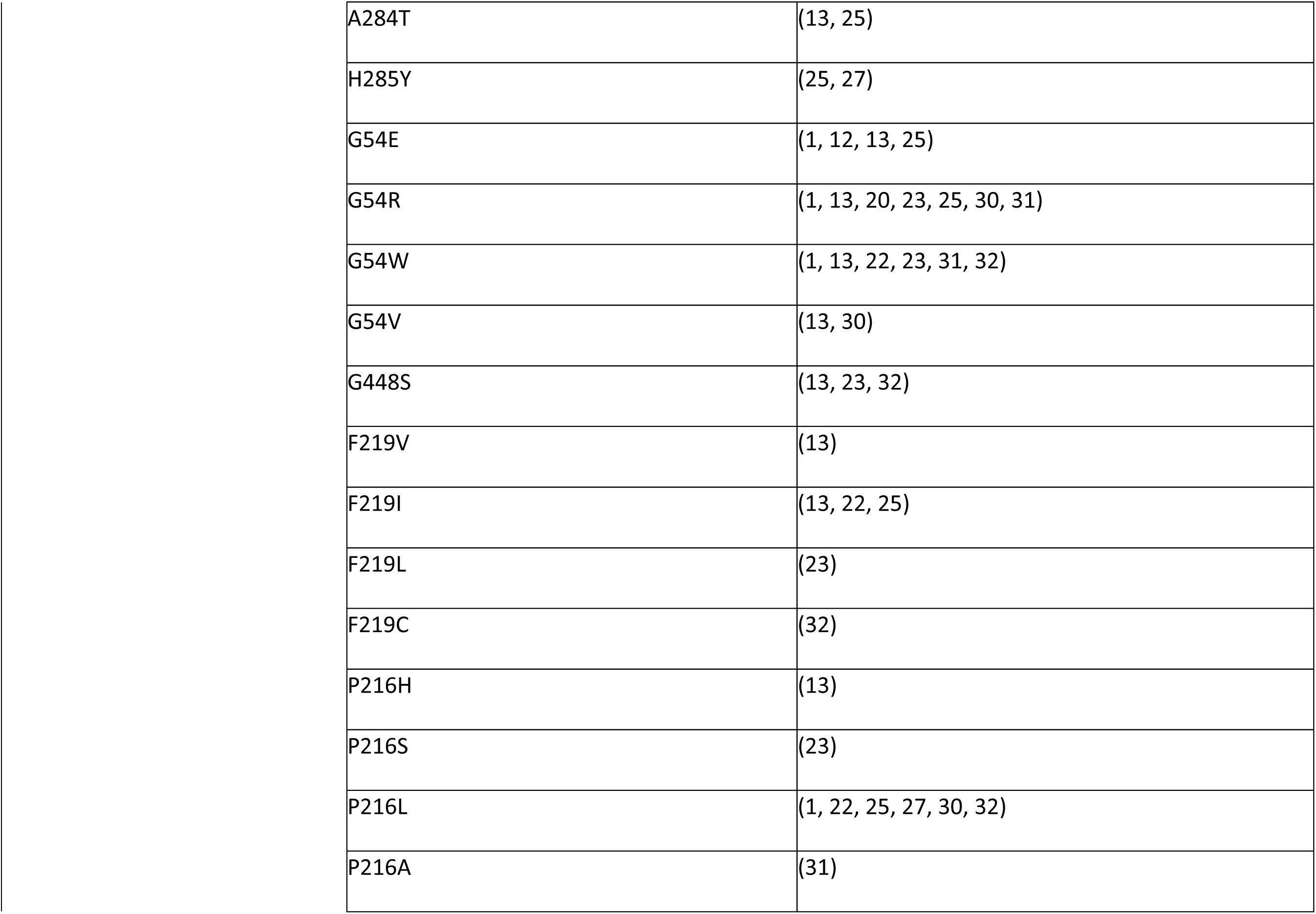
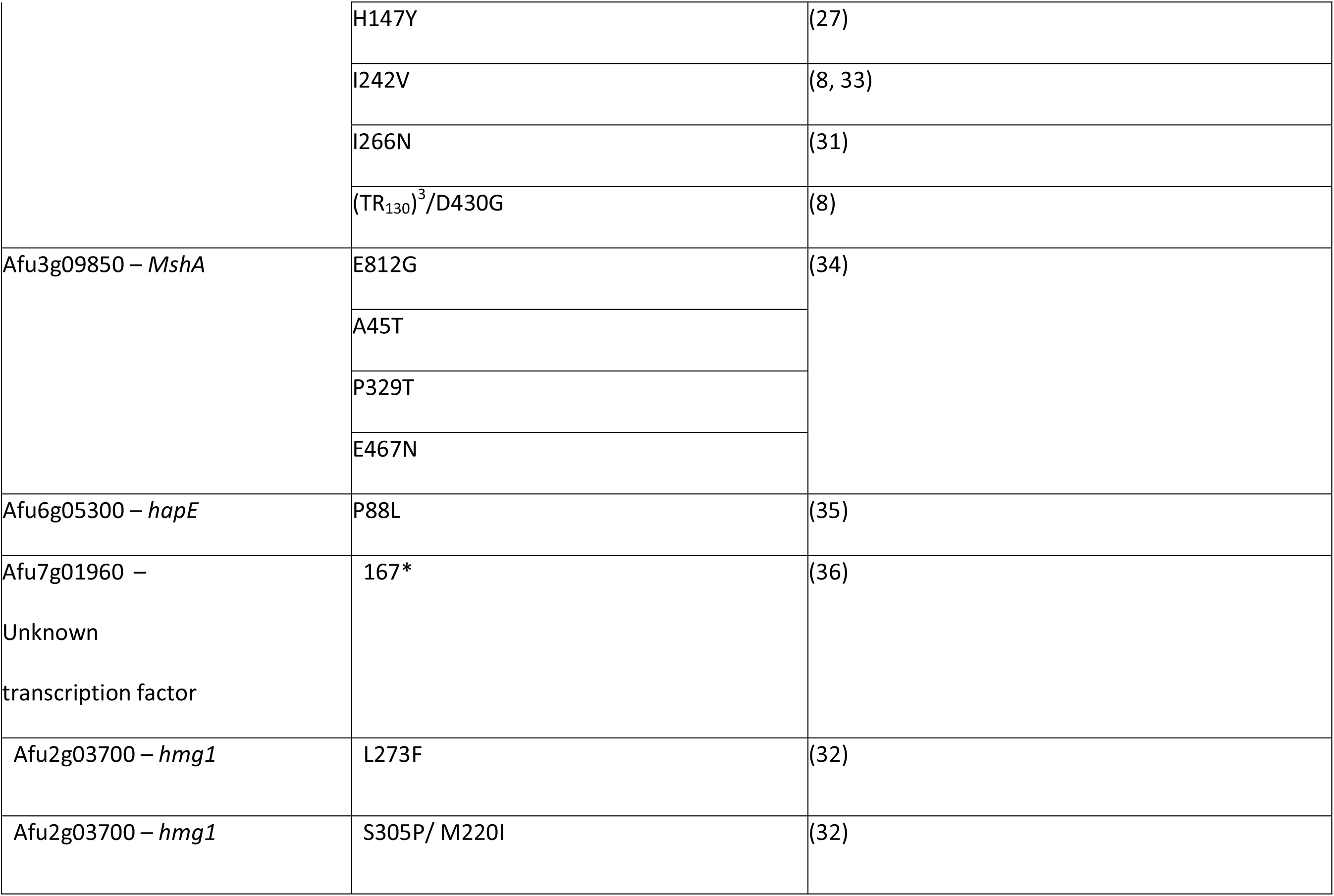
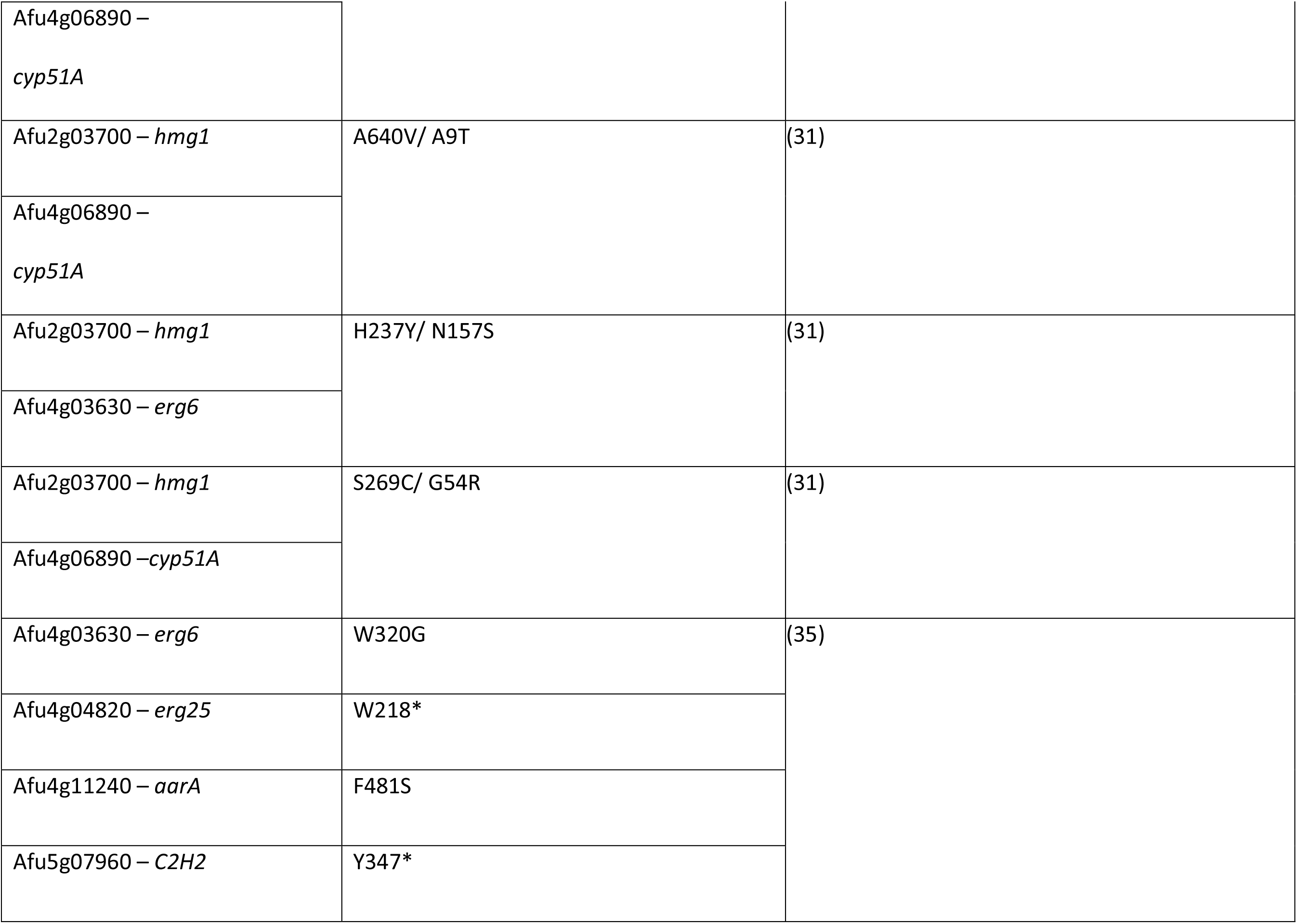
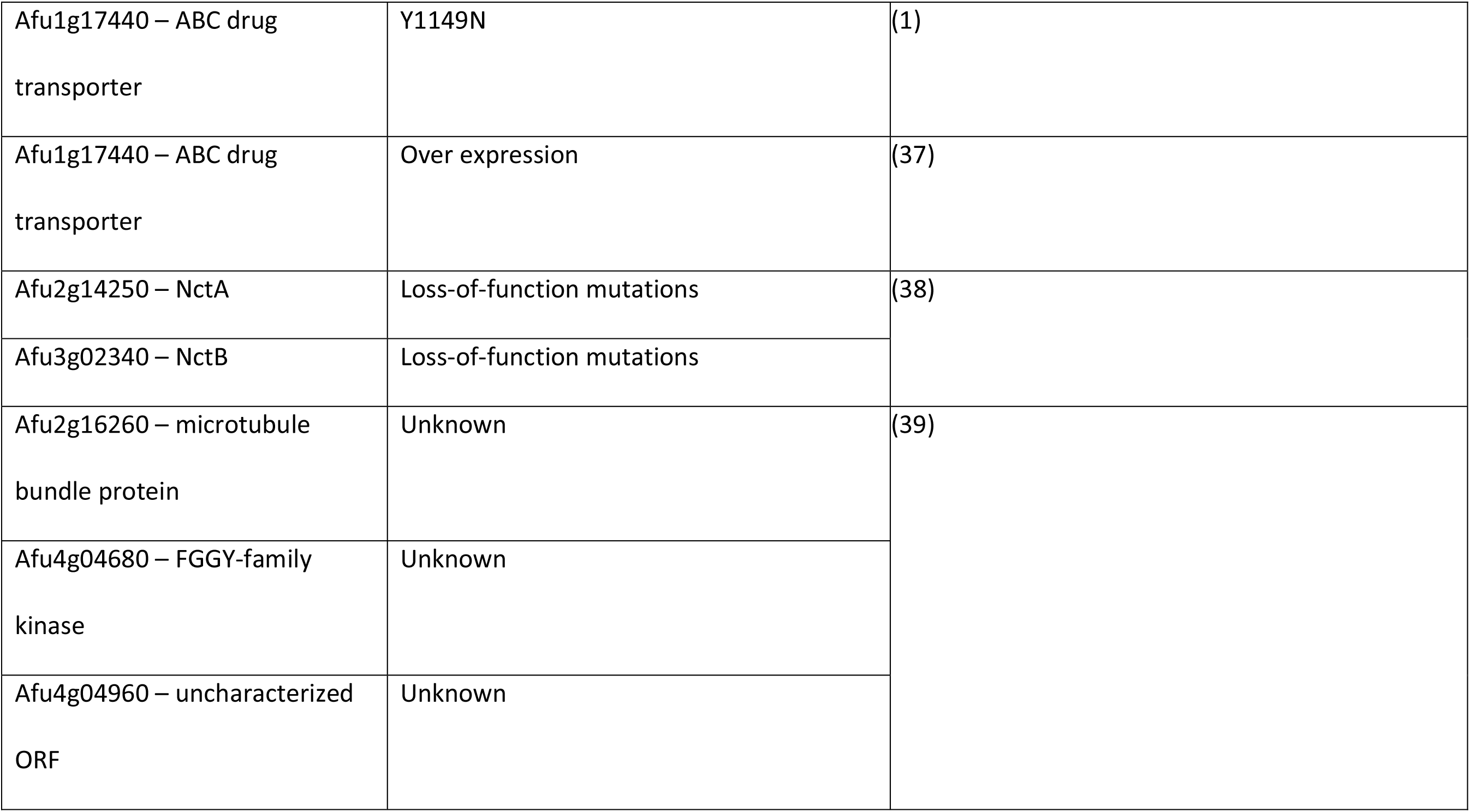
Polymorphisms reported in the literature that are associated with azole-resistance in *A. fumigatus* in the CAPA isolates used in this study.

|  |  |  |  |  |
| --- | --- | --- | --- | --- |
| Gene | Mutation |  |  | Reference |
| Afu4g06890 – *cyp51A* | TR<sub>34</sub> |  |  | (1) |
|  | TR<sub>34</sub>/L98H |  |  | (1, 7-22) |
|  | TR<sub>34</sub>/L98H/S297T/F495I |  |  | (10, 23) |
|  | TR<sub>34</sub>/L98H/T289A/I364V/G448S |  |  | (1, 8) |
|  | TR<sub>46</sub>/Y121F/T289A |  |  | (1, 8, 11, 13-17, 24) |
|  | TR<sub>46</sub>/L98H/T289A/I364V/G448S |  |  | (8) |
|  | (TR<sub>46</sub>)<sup>2</sup>/Y121F/ M172V/ T289A/ G448S |  |  | (8) |
|  | TR53 |  |  | (8) |

|  |  |
| --- | --- |
| TR120 | (8) |
| F46Y/M172V/N248T/D255E/E427K | (10, 22, 25, 26) |
| F46Y/M172V/E427K | (7) |
| G138C | (27, 28) |
| Y121F | (23) |
| Y431C | (25, 27, 28) |
| G434C | (27, 28) |
| G432S | (23) |
| M220L | (15, 25) |
| M220T | (13, 29) |
| M220R | (23, 30) |
| M220I | (23, 29, 31) |
| M220K | (12, 23) |
| M220V | (32) |
| N248K | (10, 12, 20, 33) |

|  |  |
| --- | --- |
| A284T | (13, 25) |
| H285Y | (25, 27) |
| G54E | (1, 12, 13, 25) |
| G54R | (1, 13, 20, 23, 25, 30, 31) |
| G54W | (1, 13, 22, 23, 31, 32) |
| G54V | (13, 30) |
| G448S | (13, 23, 32) |
| F219V | (13) |
| F219I | (13, 22, 25) |
| F219L | (23) |
| F219C | (32) |
| P216H | (13) |
| P216S | (23) |
| P216L | (1, 22, 25, 27, 30, 32) |
| P216A | (31) |

|  |  |  |  |
| --- | --- | --- | --- |
|  | H147Y |  | (27) |
|  | I242V |  | (8, 33) |
|  | I266N |  | (31) |
|  | (TR<sub>130</sub>)<sup>3</sup>/D430G |  | (8) |
| Afu3g09850 – *MshA* | E812G |  | (34) |
|  | A45T |  |  |
|  | P329T |  |  |
|  | E467N |  |  |
| Afu6g05300 – *hapE* | P88L |  | (35) |
| Afu7g01960 –<br>Unknown<br>transcription factor | 167* |  | (36) |
| Afu2g03700 – *hmg1* | L273F |  | (32) |
| Afu2g03700 – *hmg1* | S305P/ M220I |  | (32) |

|  |  |  |
| --- | --- | --- |
| Afu4g06890 –<br>*cyp51A* |  |  |
| Afu2g03700 – *hmg1* | A640V/ A9T | (31) |
| Afu4g06890 –<br>*cyp51A* |  |  |
| Afu2g03700 – *hmg1* | H237Y/ N157S | (31) |
| Afu4g03630 – *erg6* |  |  |
| Afu2g03700 – *hmg1* | S269C/ G54R | (31) |
| Afu4g06890 –*cyp51A* |  |  |
| Afu4g03630 – *erg6* | W320G | (35) |
| Afu4g04820 – *erg25* | W218* |  |
| Afu4g11240 – *aara* | F481S |  |
| Afu5g07960 – C2H2 | Y347* |  |
| Afu1g17440 – ABC drug transporter | Y1149N | (1) |
| Afu1g17440 – ABC drug transporter | Over expression | (37) |
| Afu2g14250 – NctA | Loss-of-function mutations | (38) |
| Afu3g02340 – NctB | Loss-of-function mutations |  |
| Afu2g16260 – microtubule bundle protein | Unknown | (39) |
| Afu4g04680 – FGGY-family kinase | Unknown |  |
| Afu4g04960 – uncharacterized ORF | Unknown |  |

**Supp Info Table 2.**
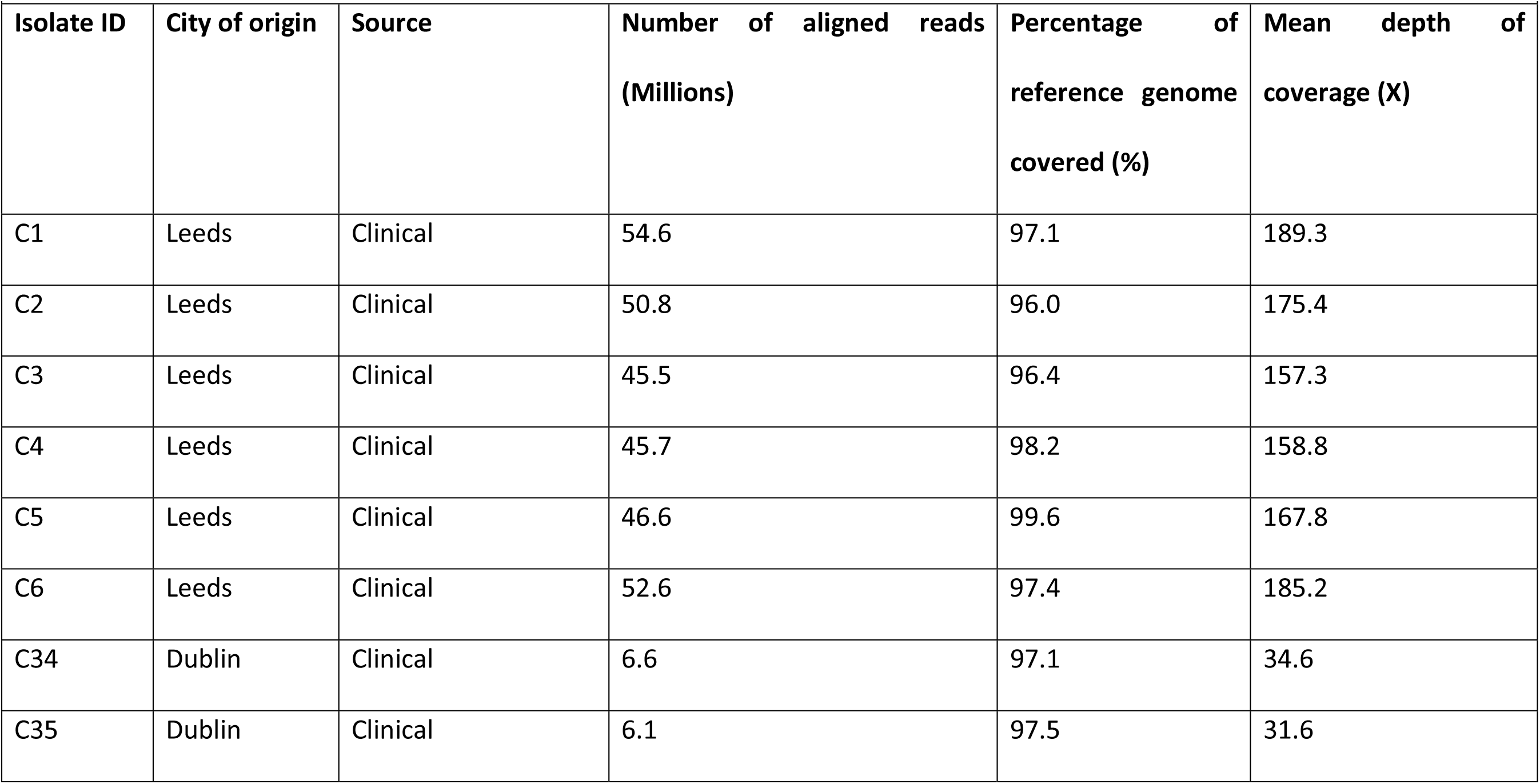
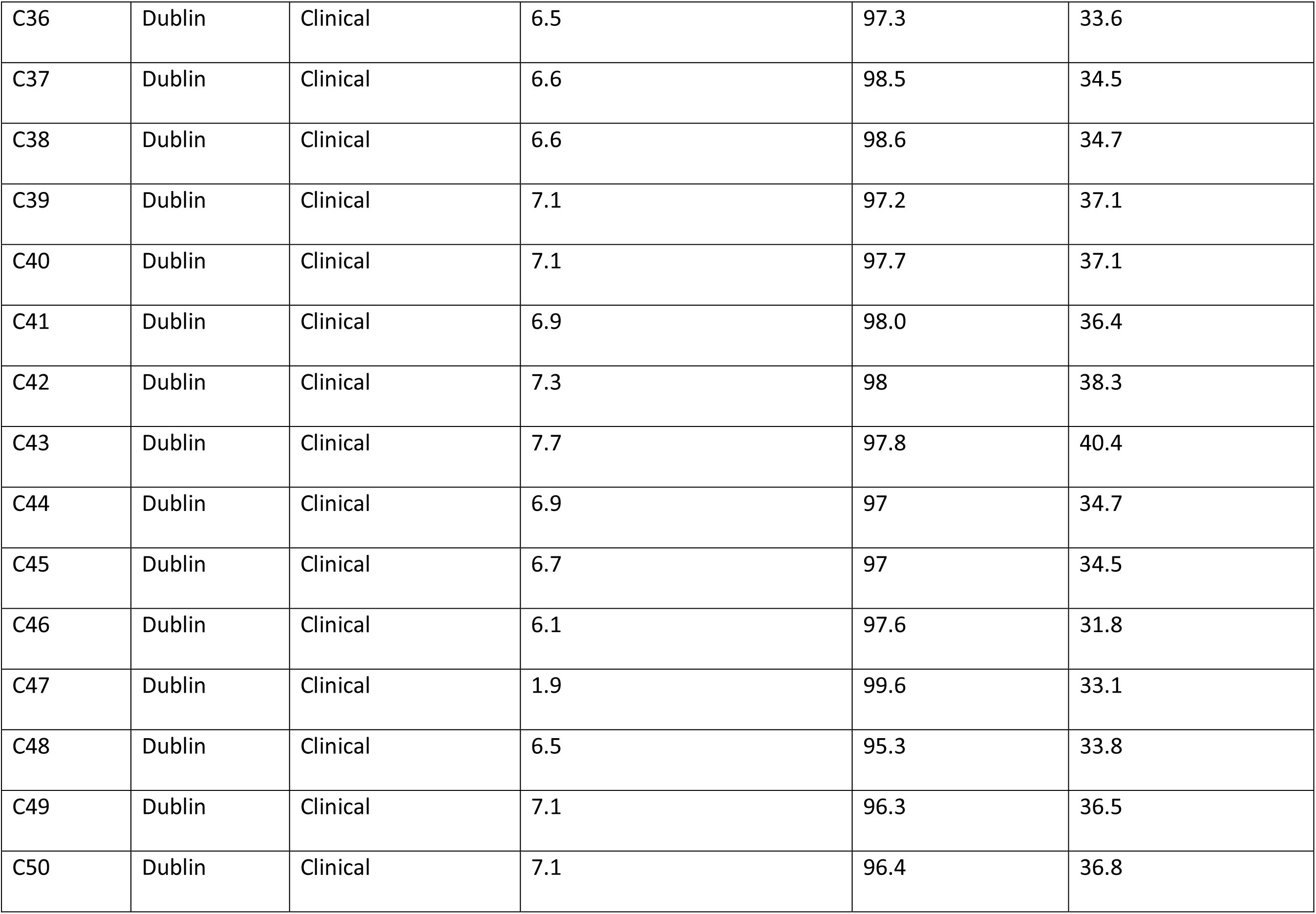
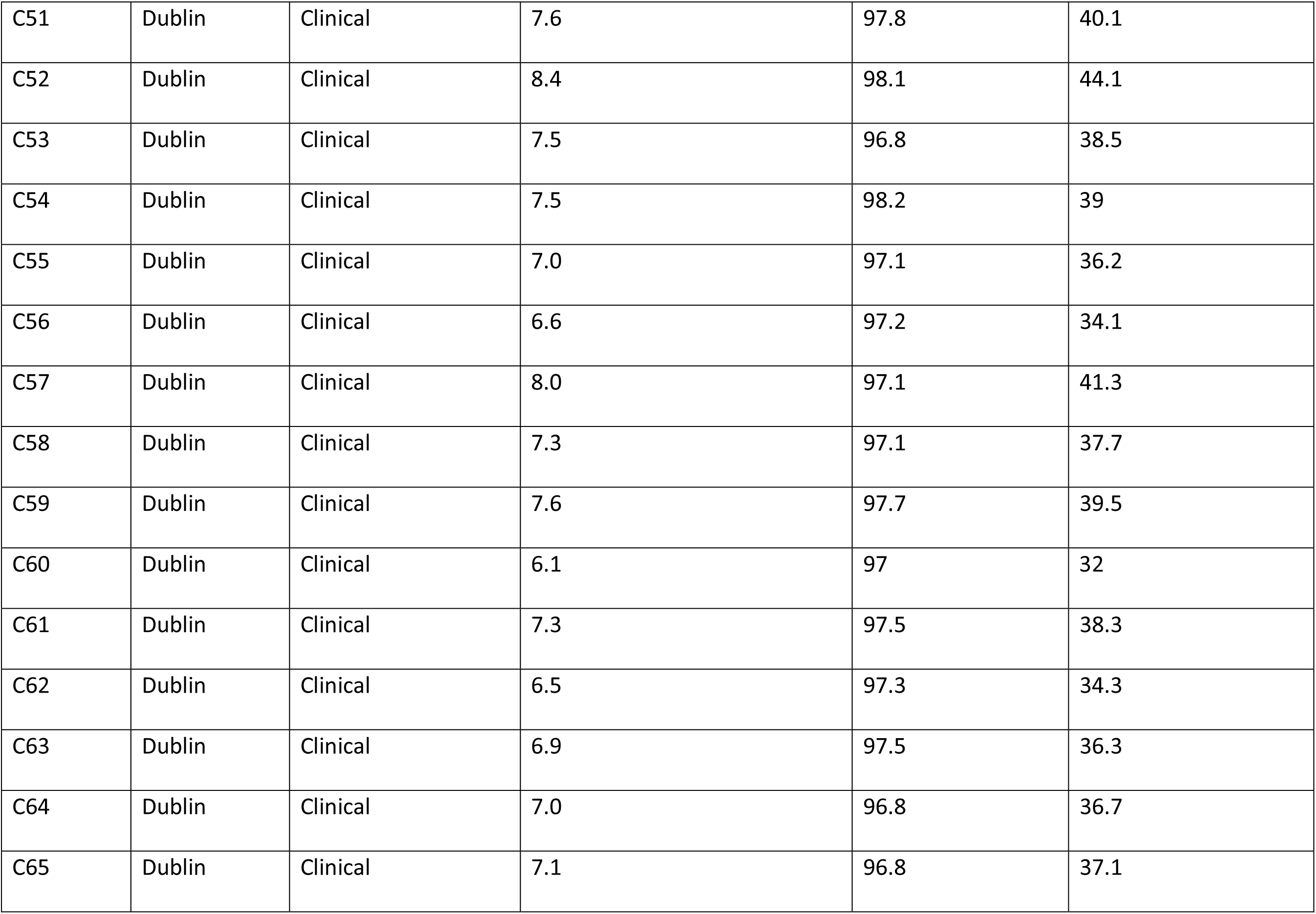
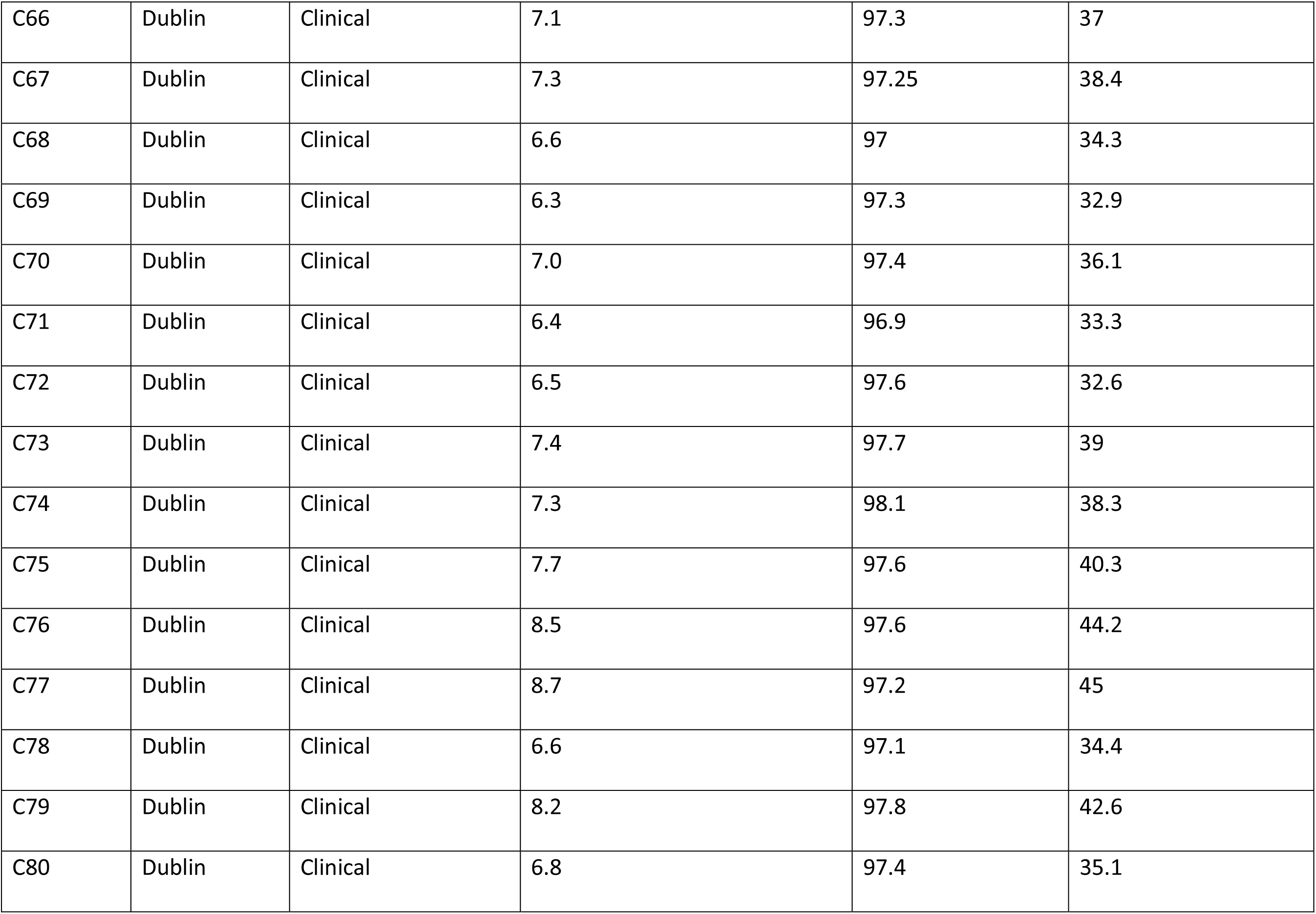
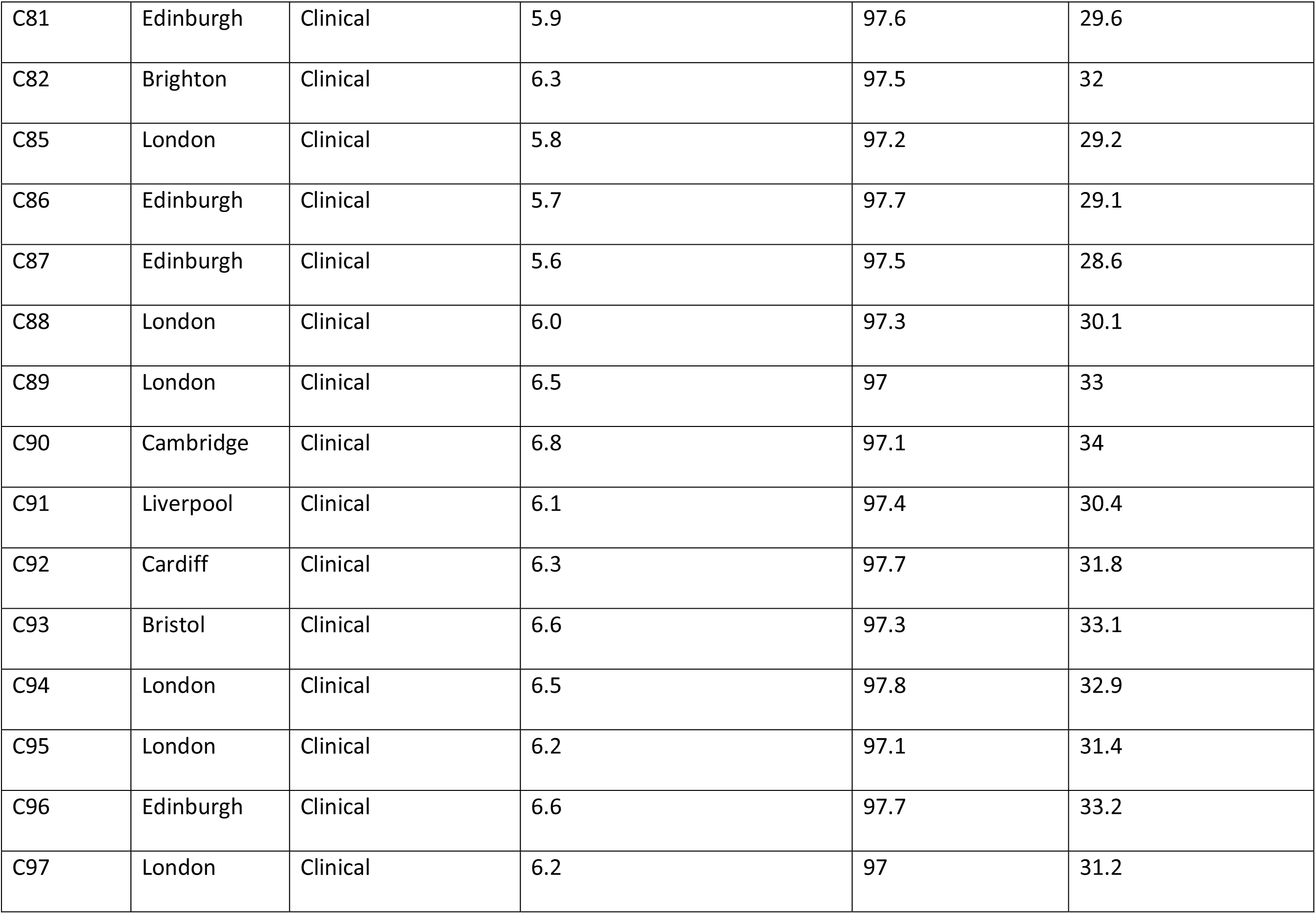
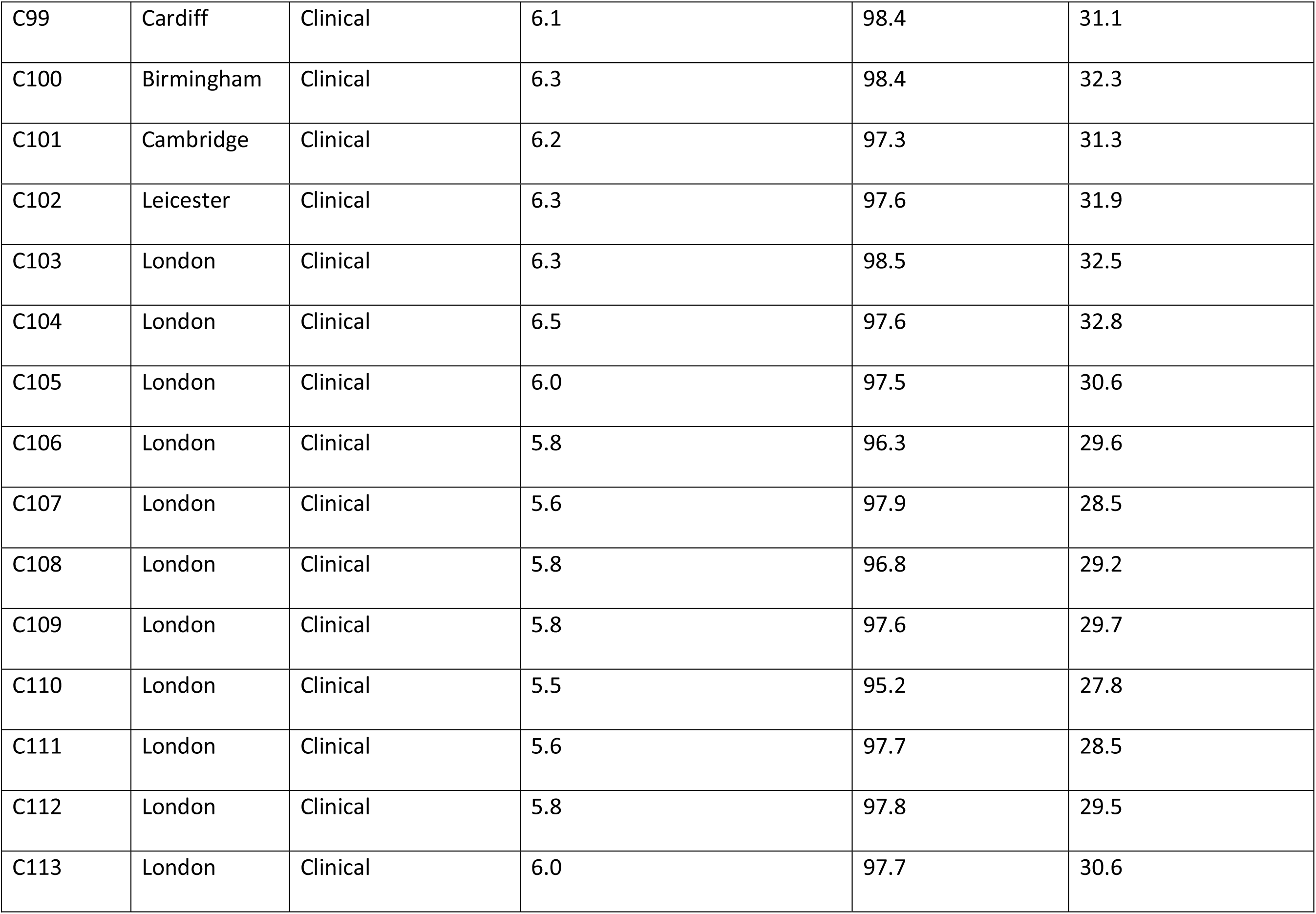
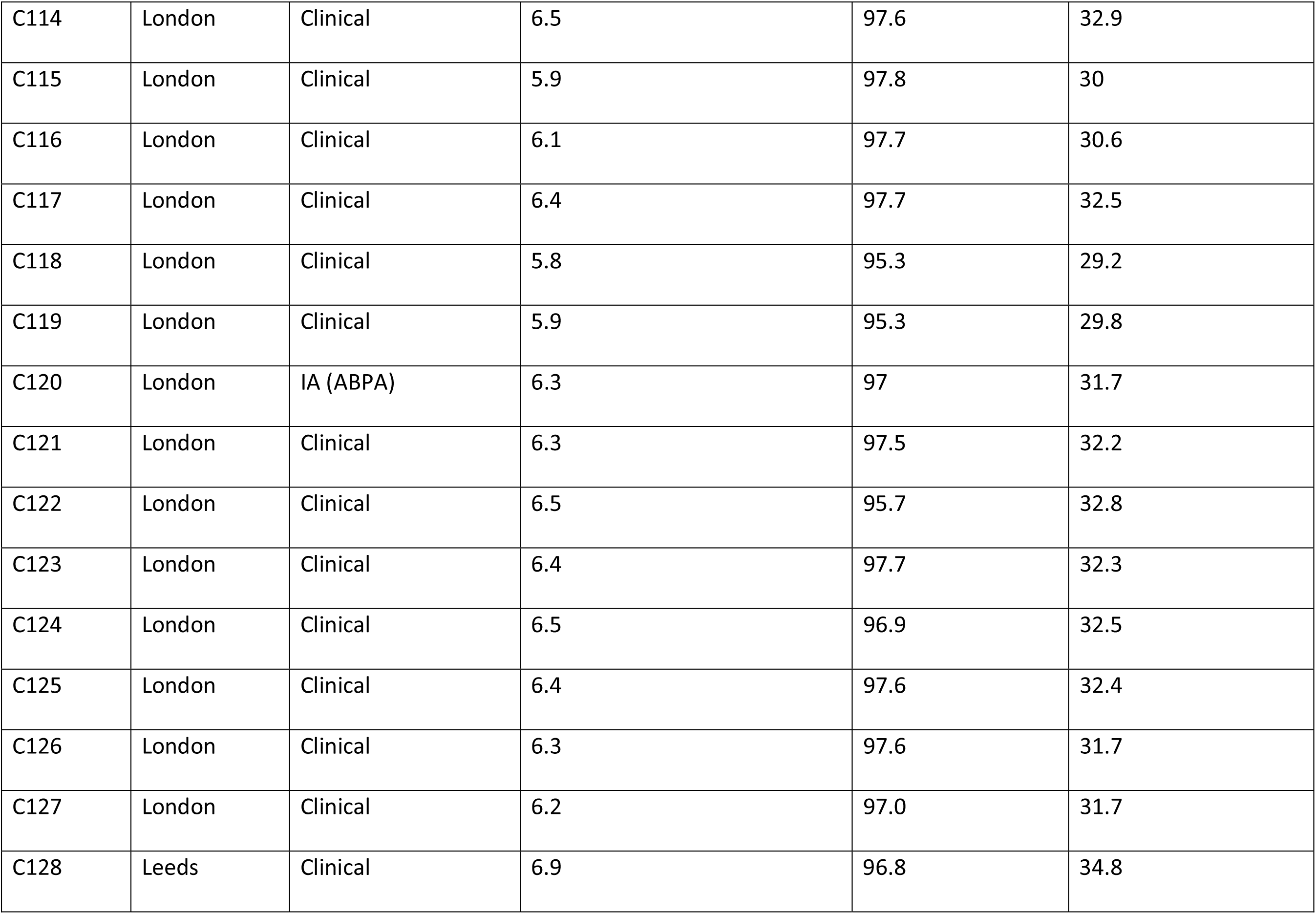
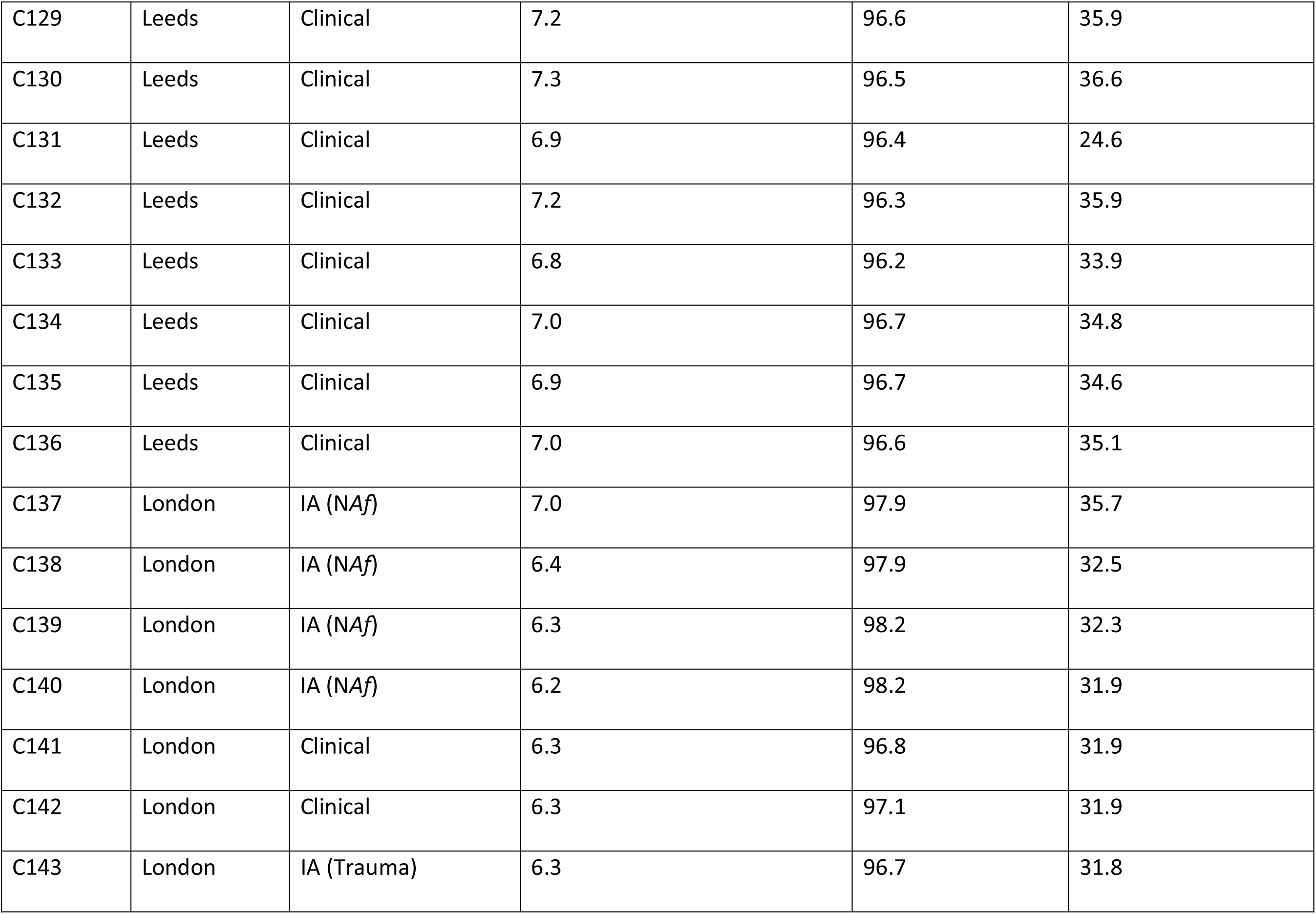
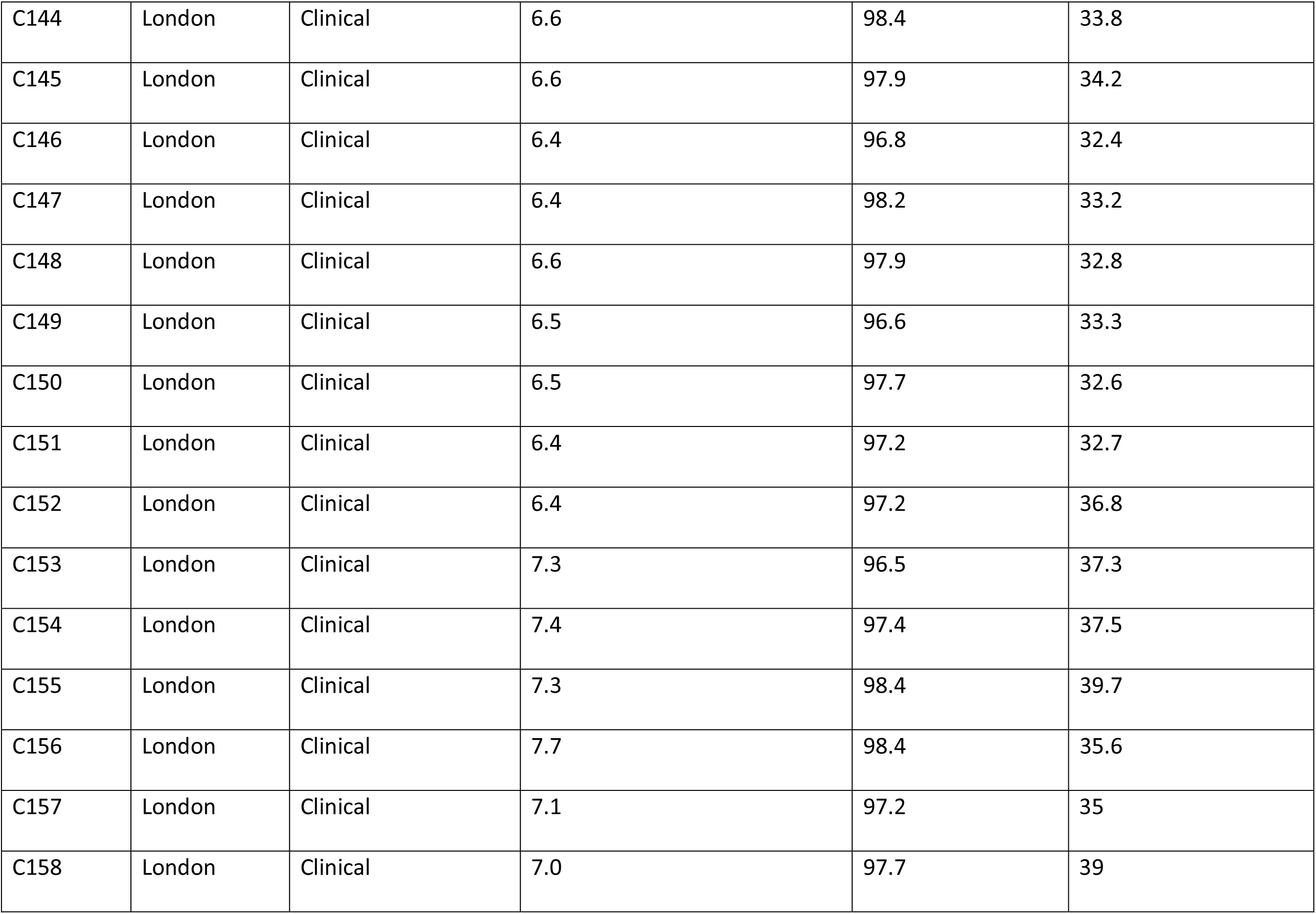
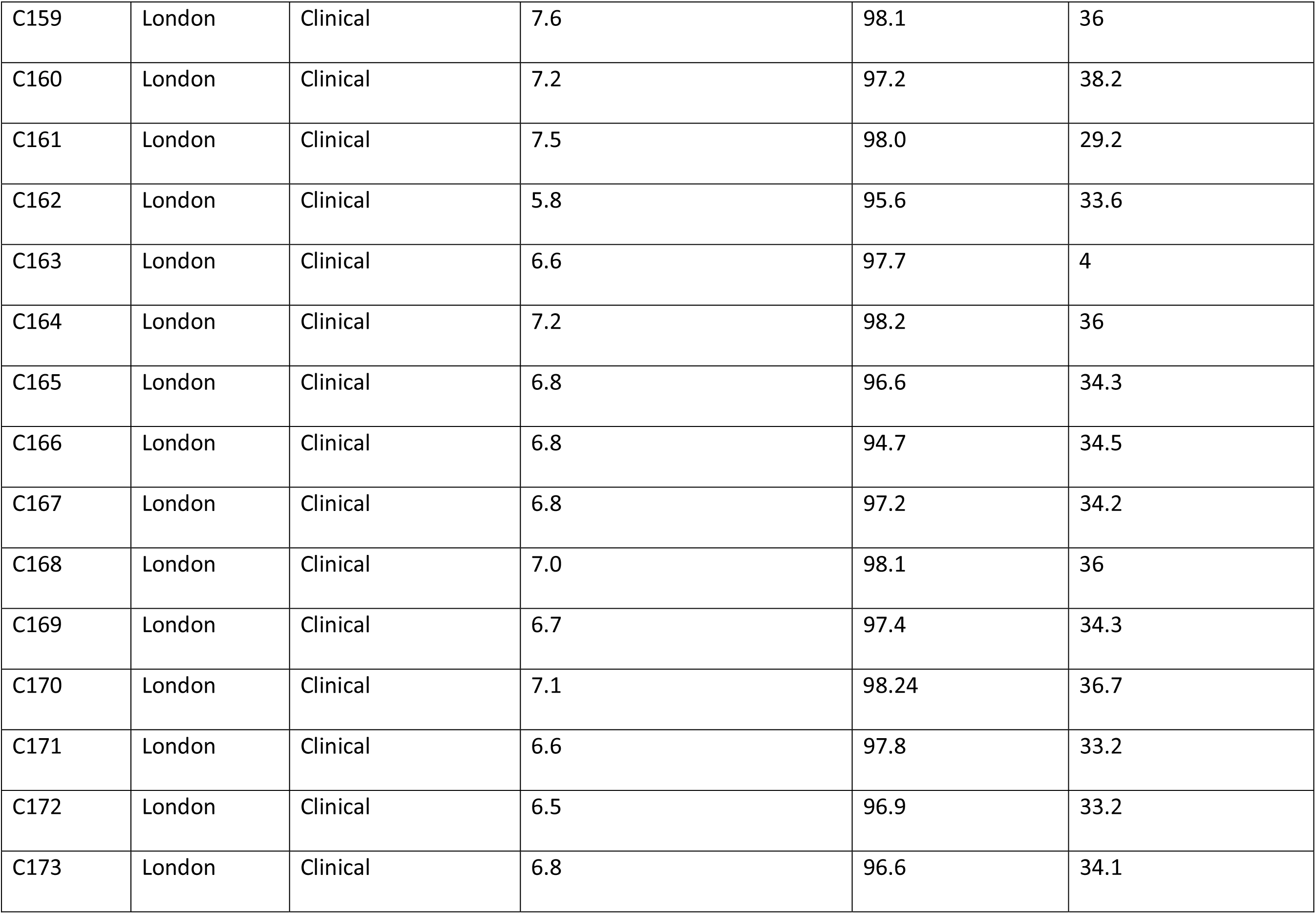
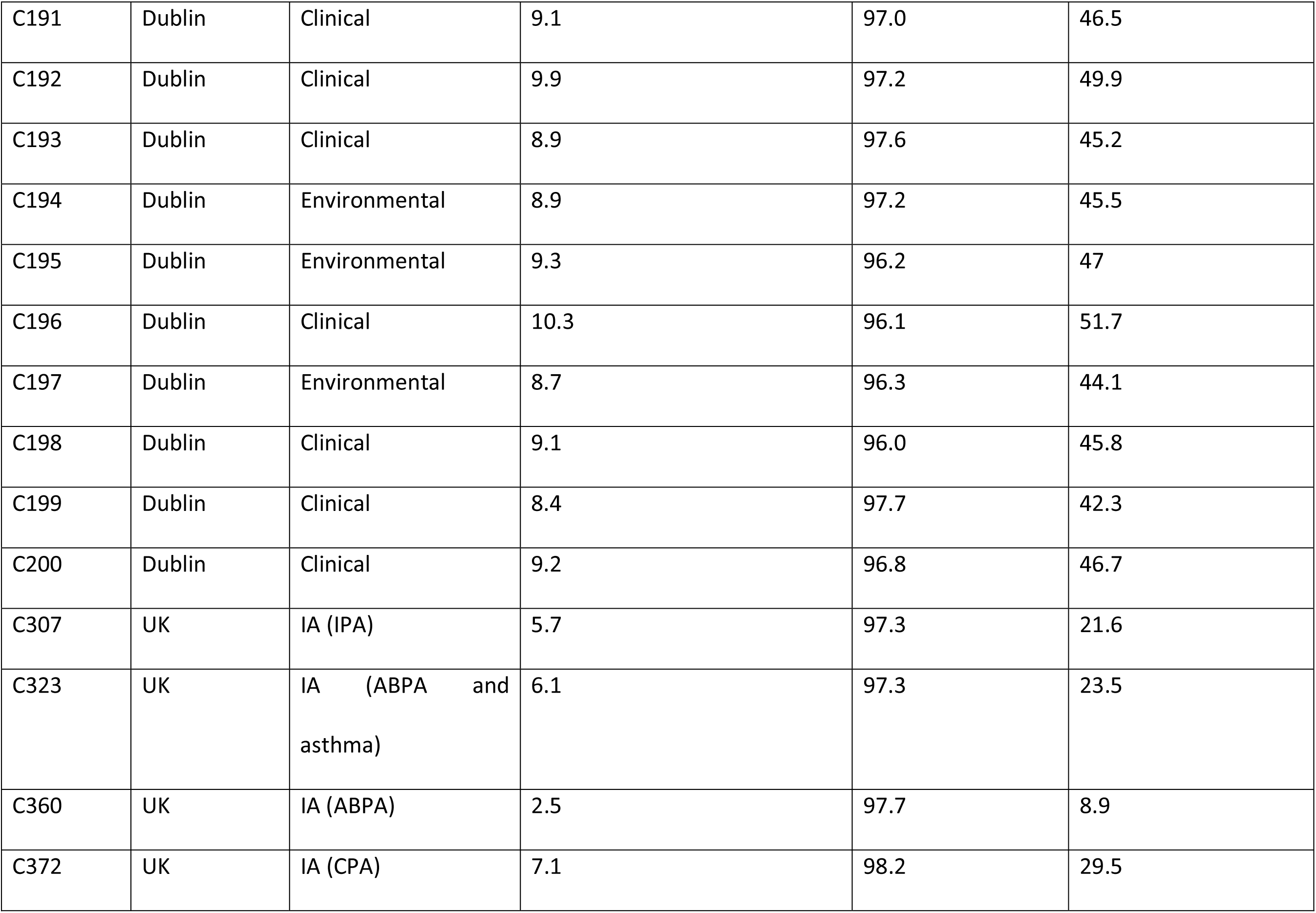
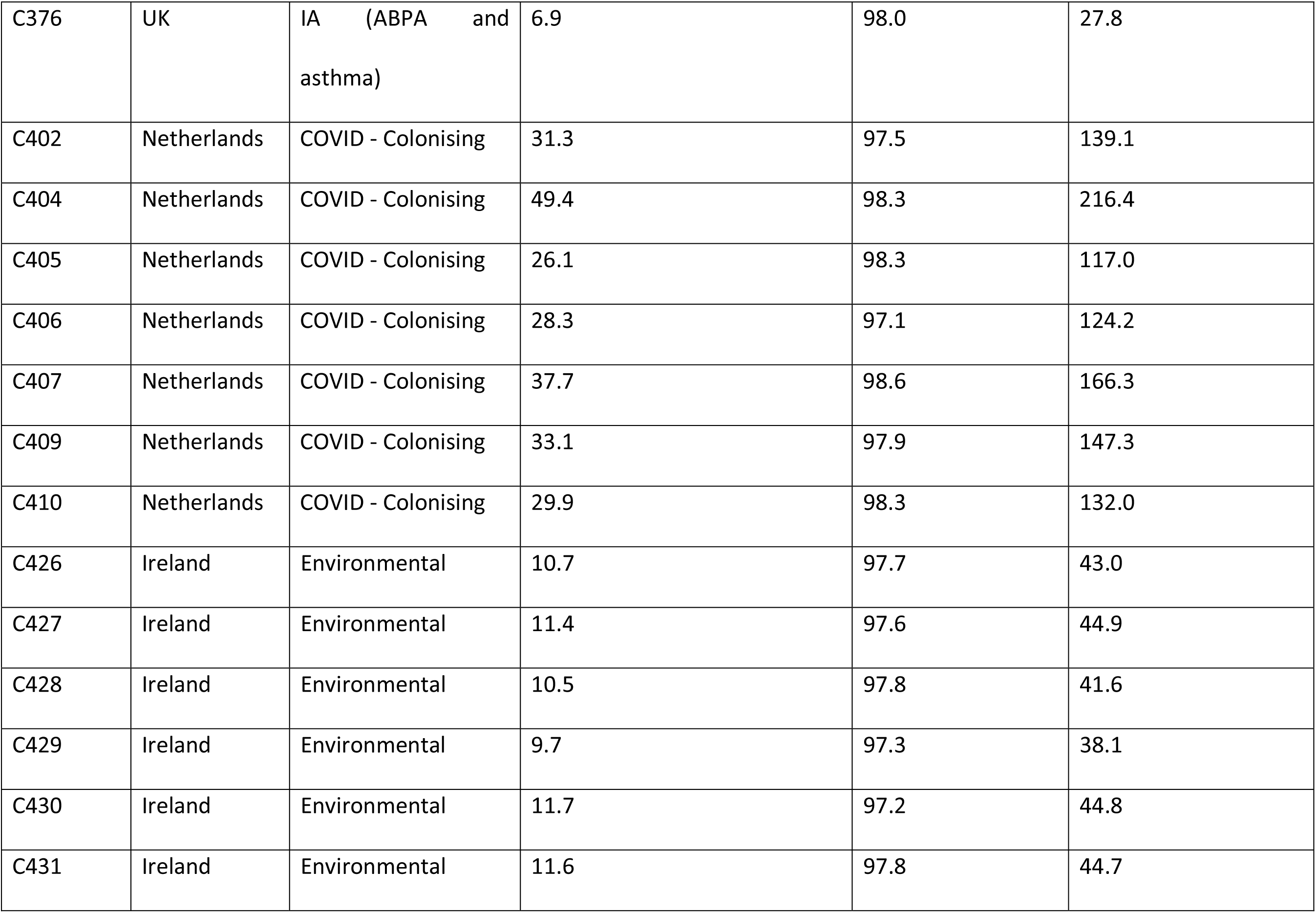
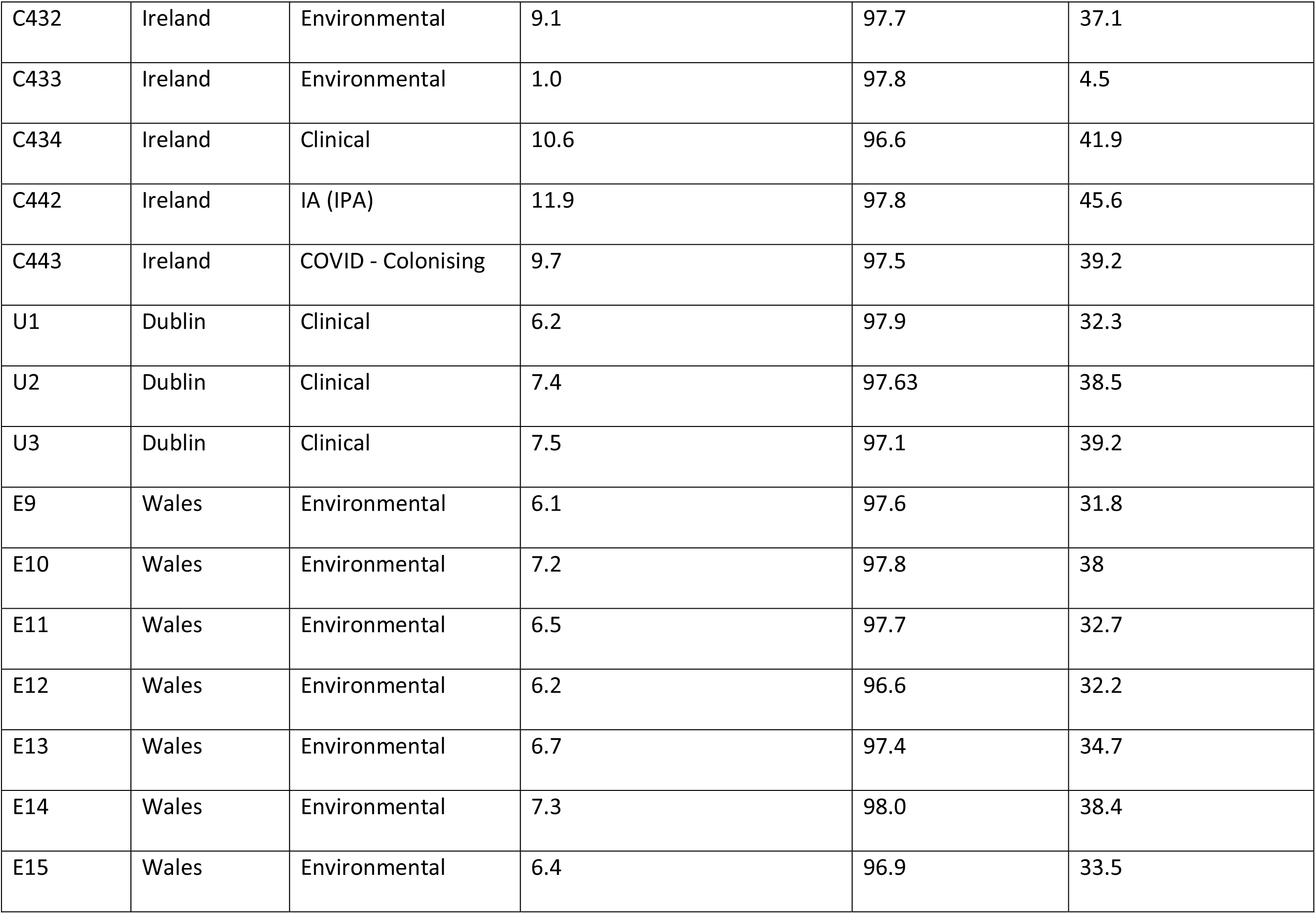
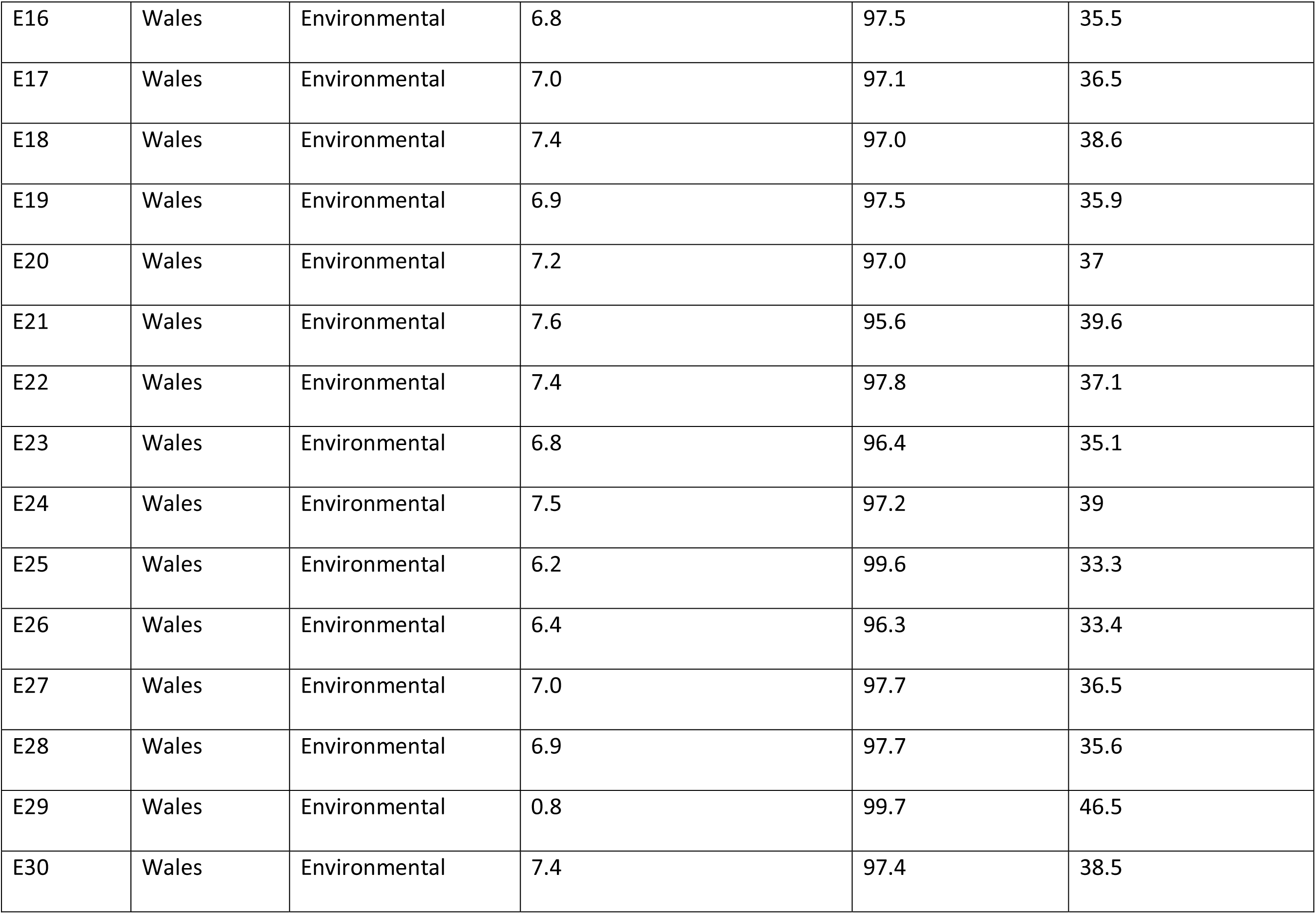
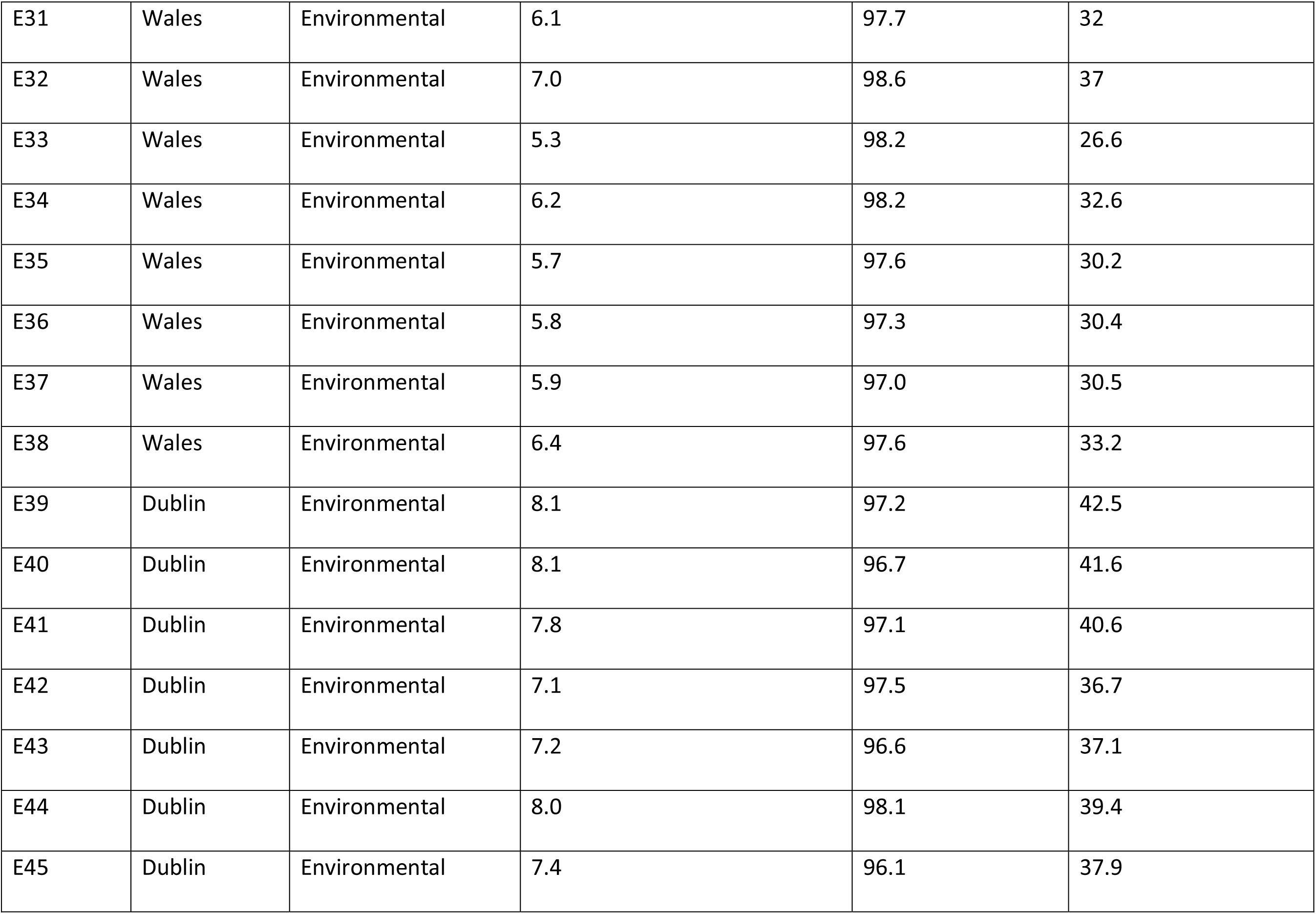
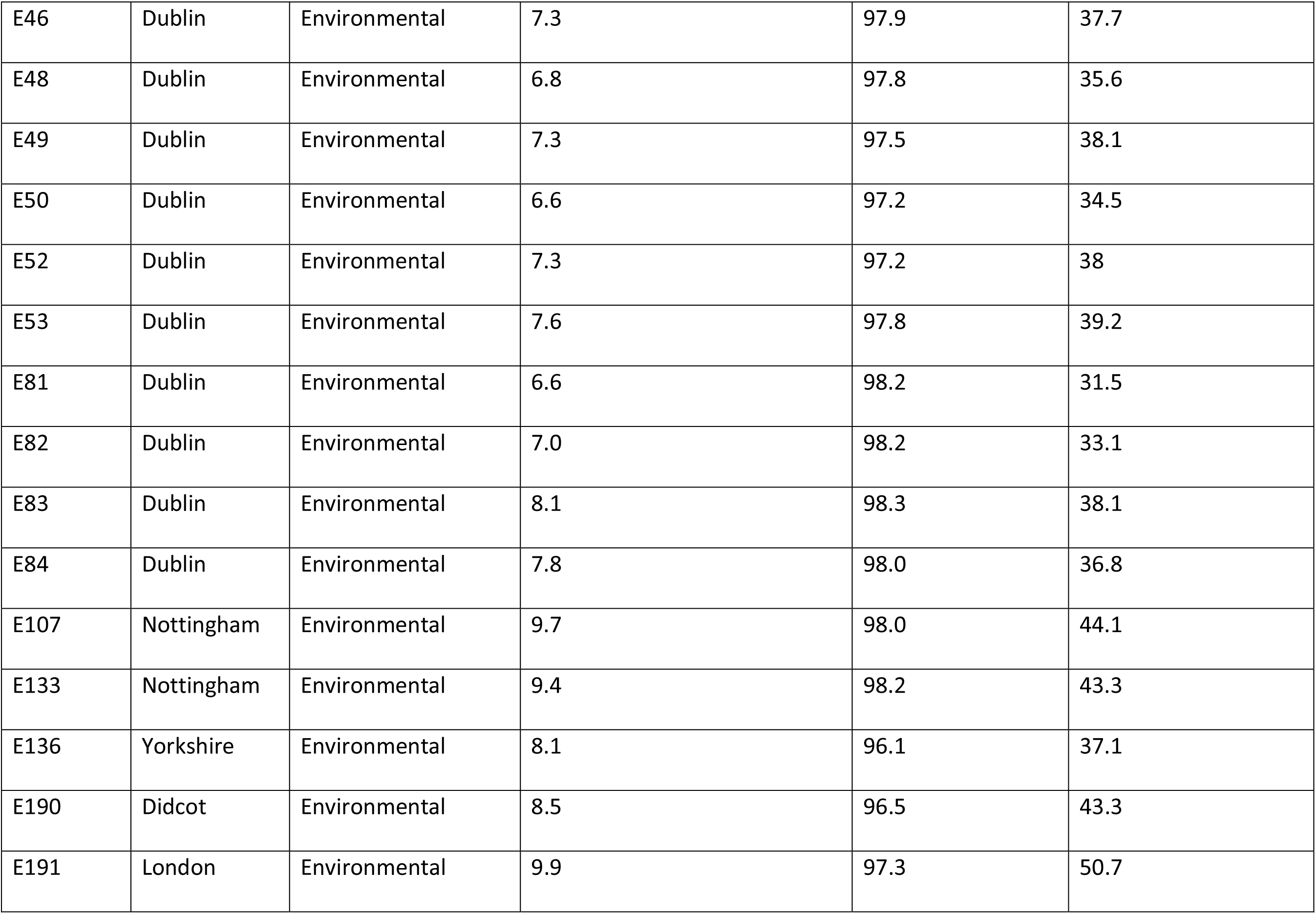
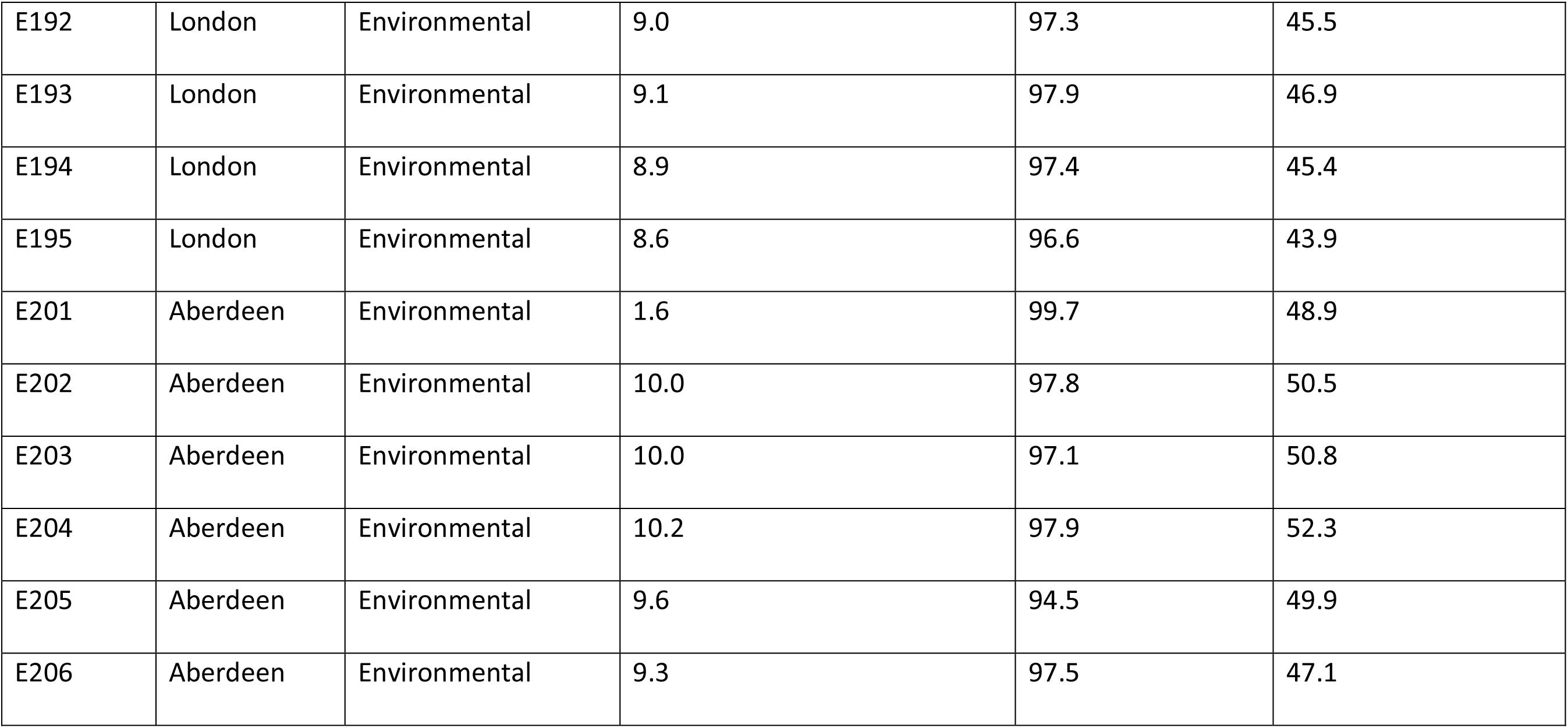
Characteristics of additional clinical and environmental isolates of *A. fumigatus* used in this study to provide context for CAPA isolates, and details of WGS alignments. ABPA, allergic bronchopulmonary aspergillosis. CPA, chronic pulmonary aspergillosis. IA, invasive aspergillosis. IPA, invasive pulmonary aspergillosis. N*Af*, necrotising *A. fumigatus*.

**Supp Info Table 3.**
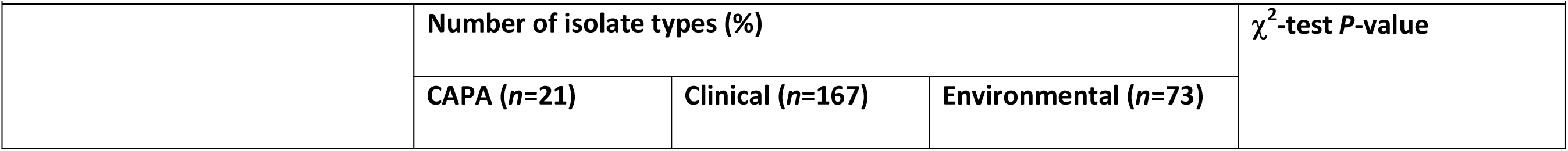
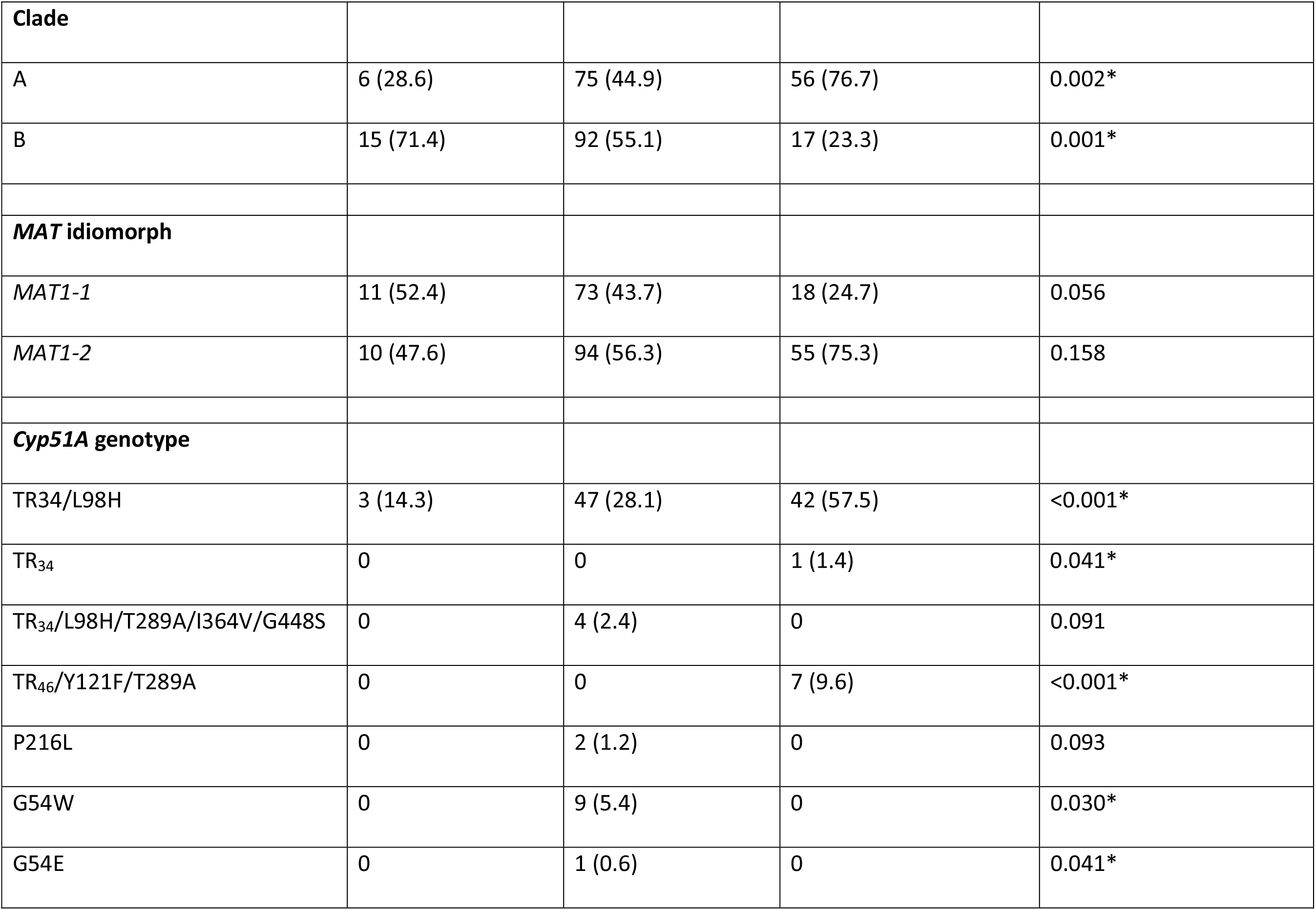
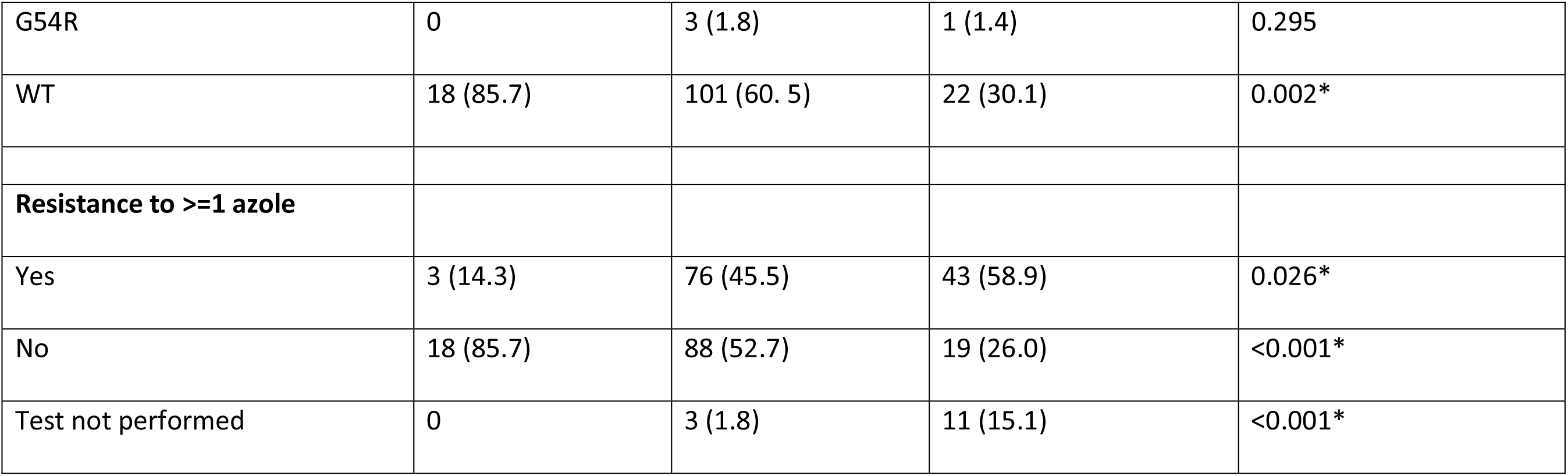
Relative frequencies of clade assignment, mating type idiomorph and *cyp51A* polymorphisms for CAPA, clinical and environmental *A. fumigatus* isolates in this study. (*P*-value*, is significant if the value p<0.05).

**Supp Info Table 4.**
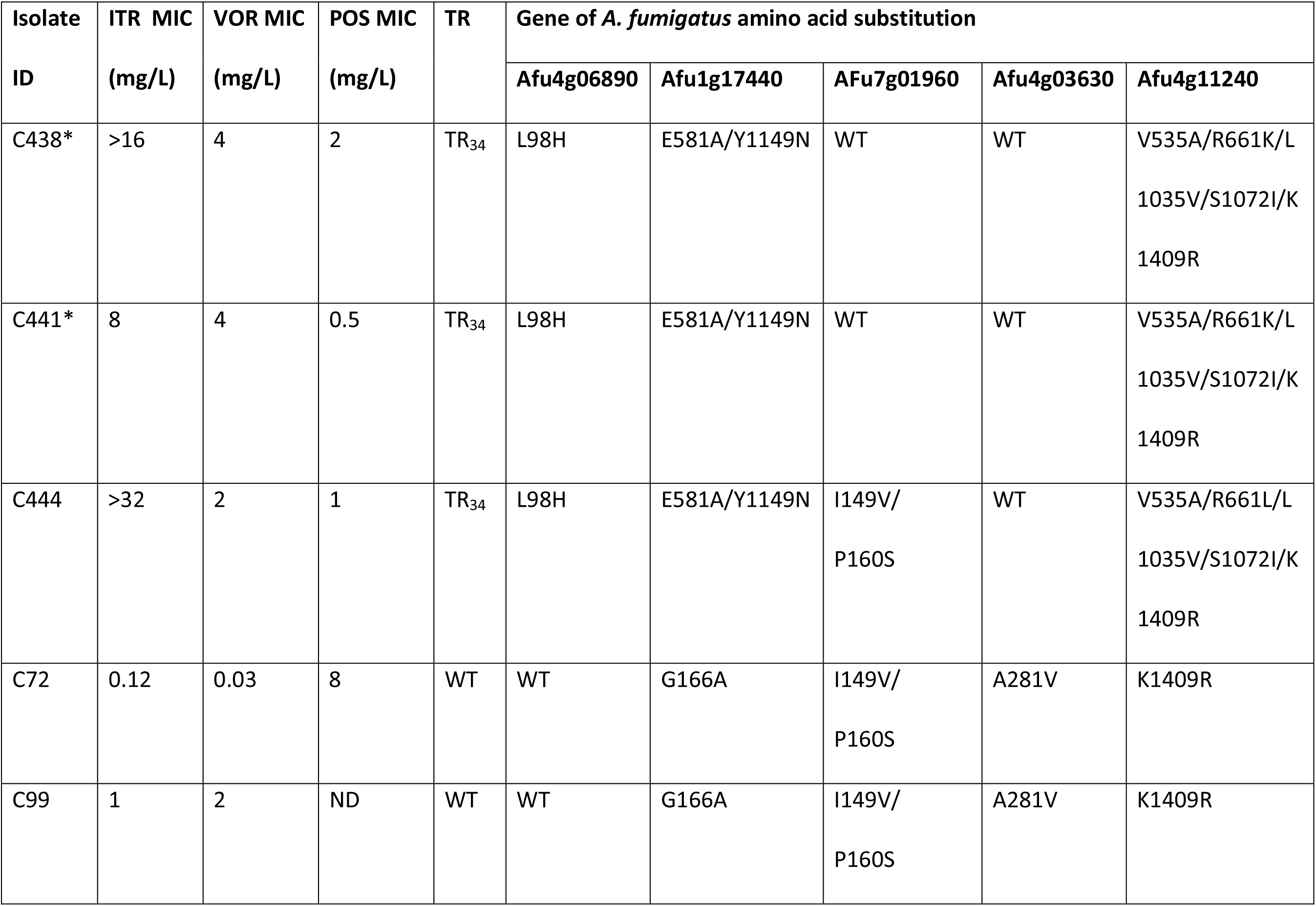
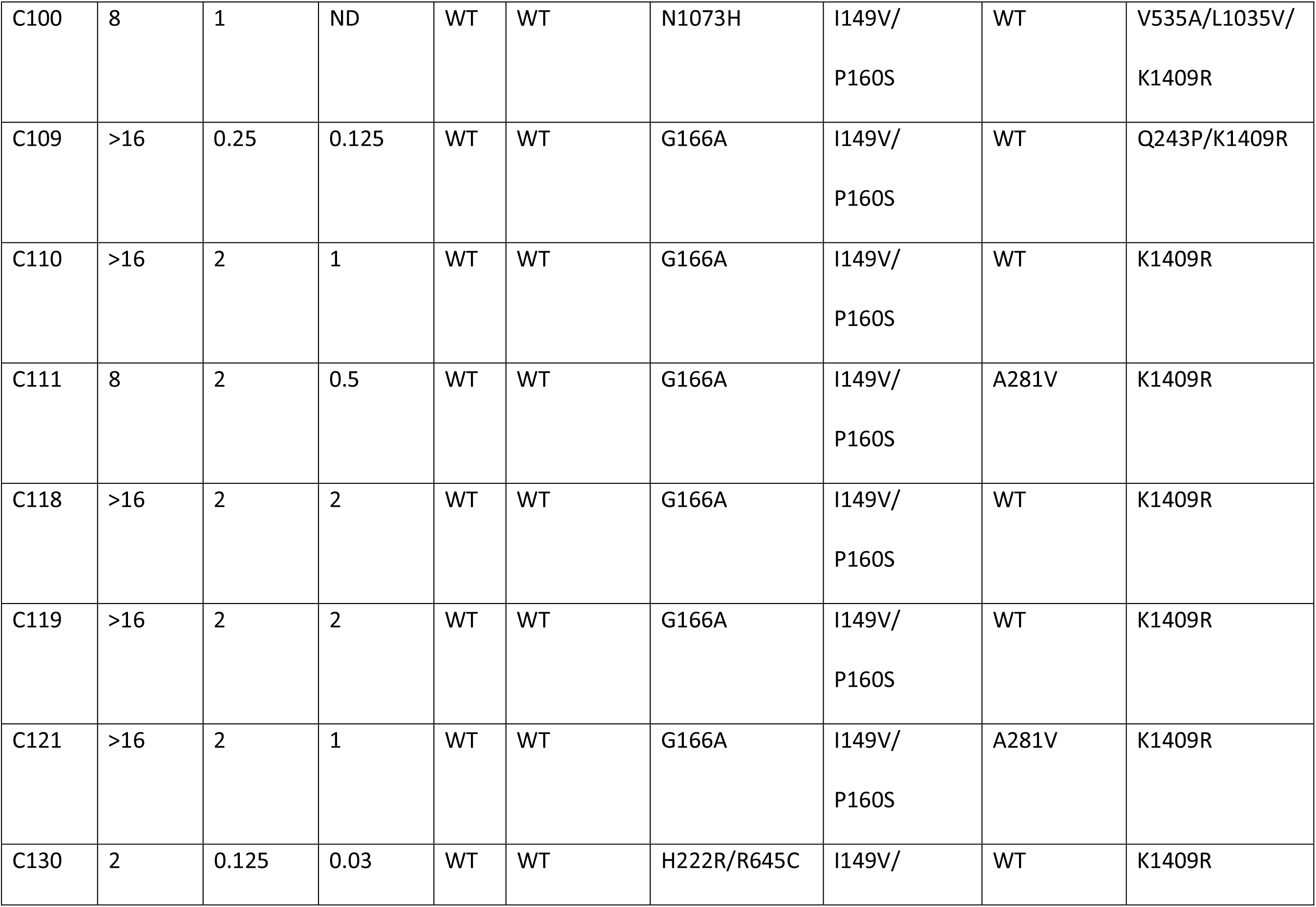
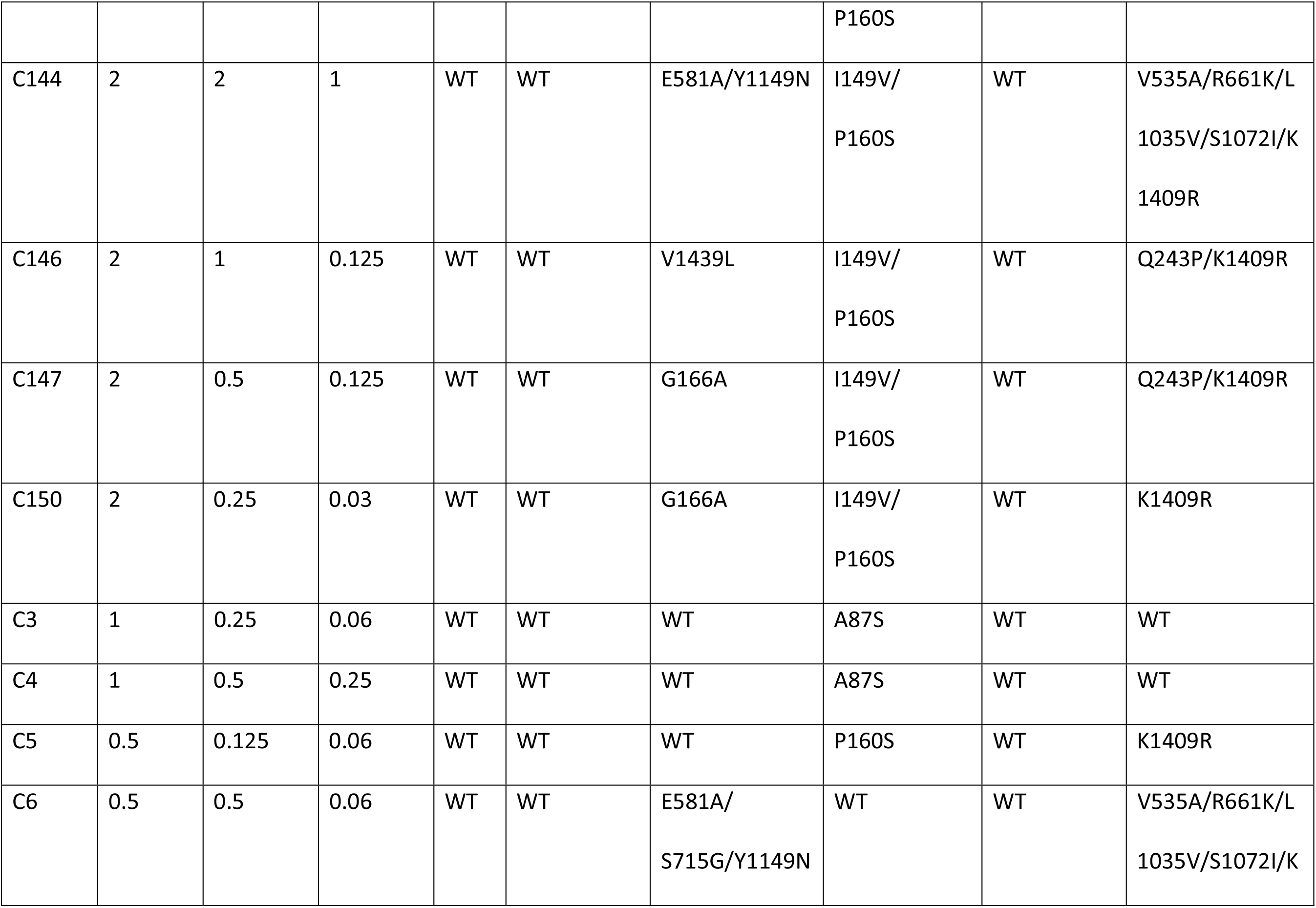
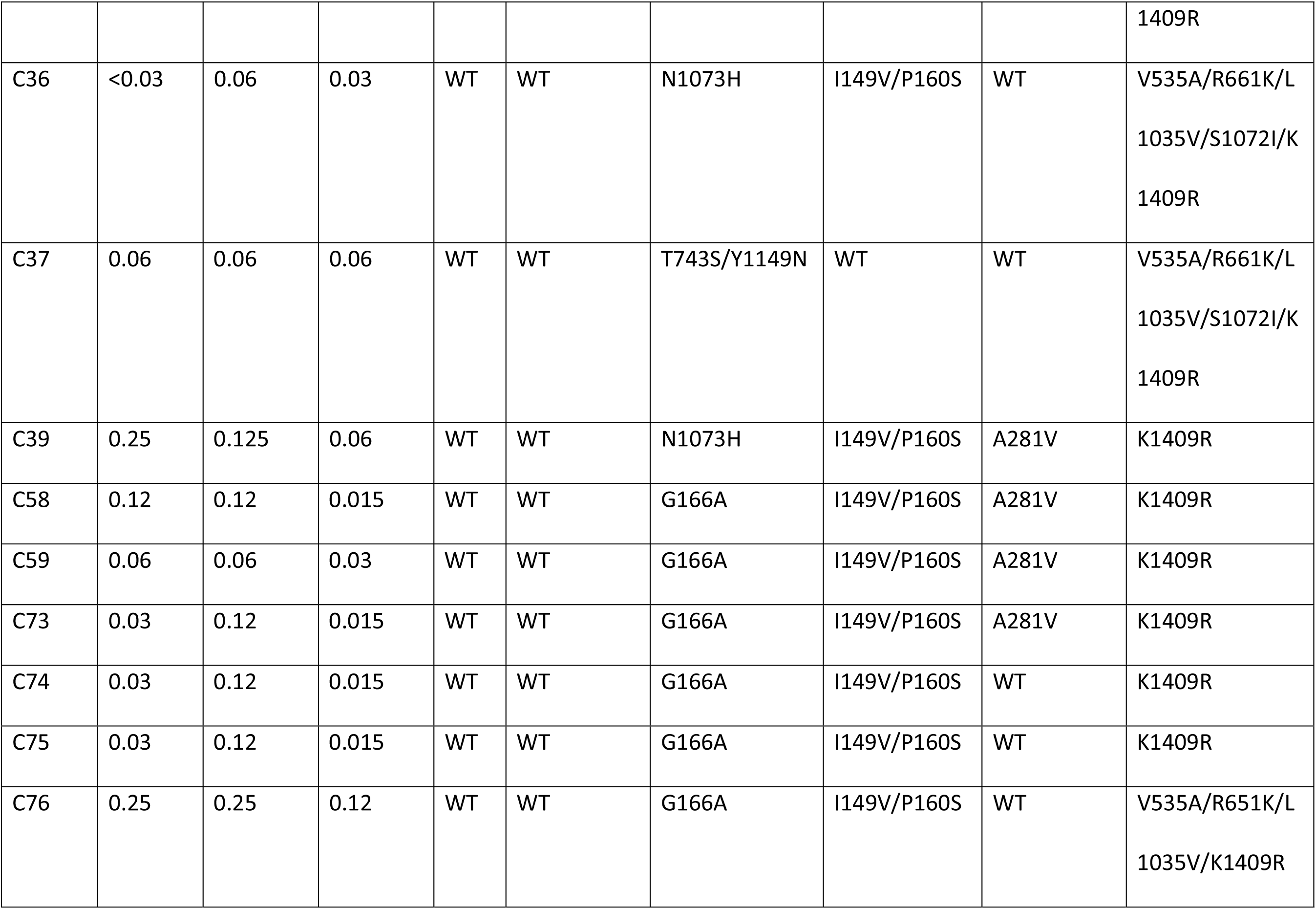
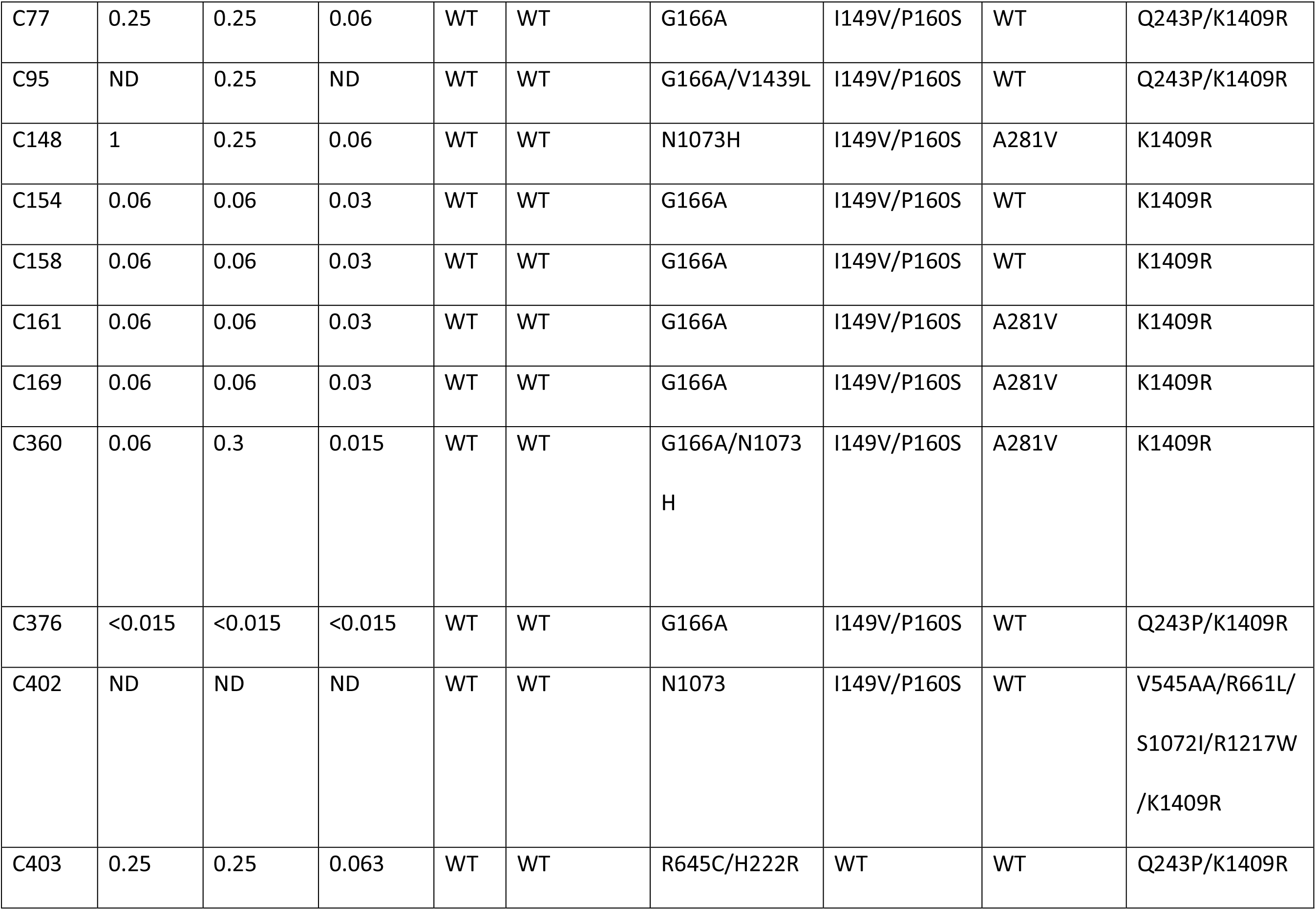
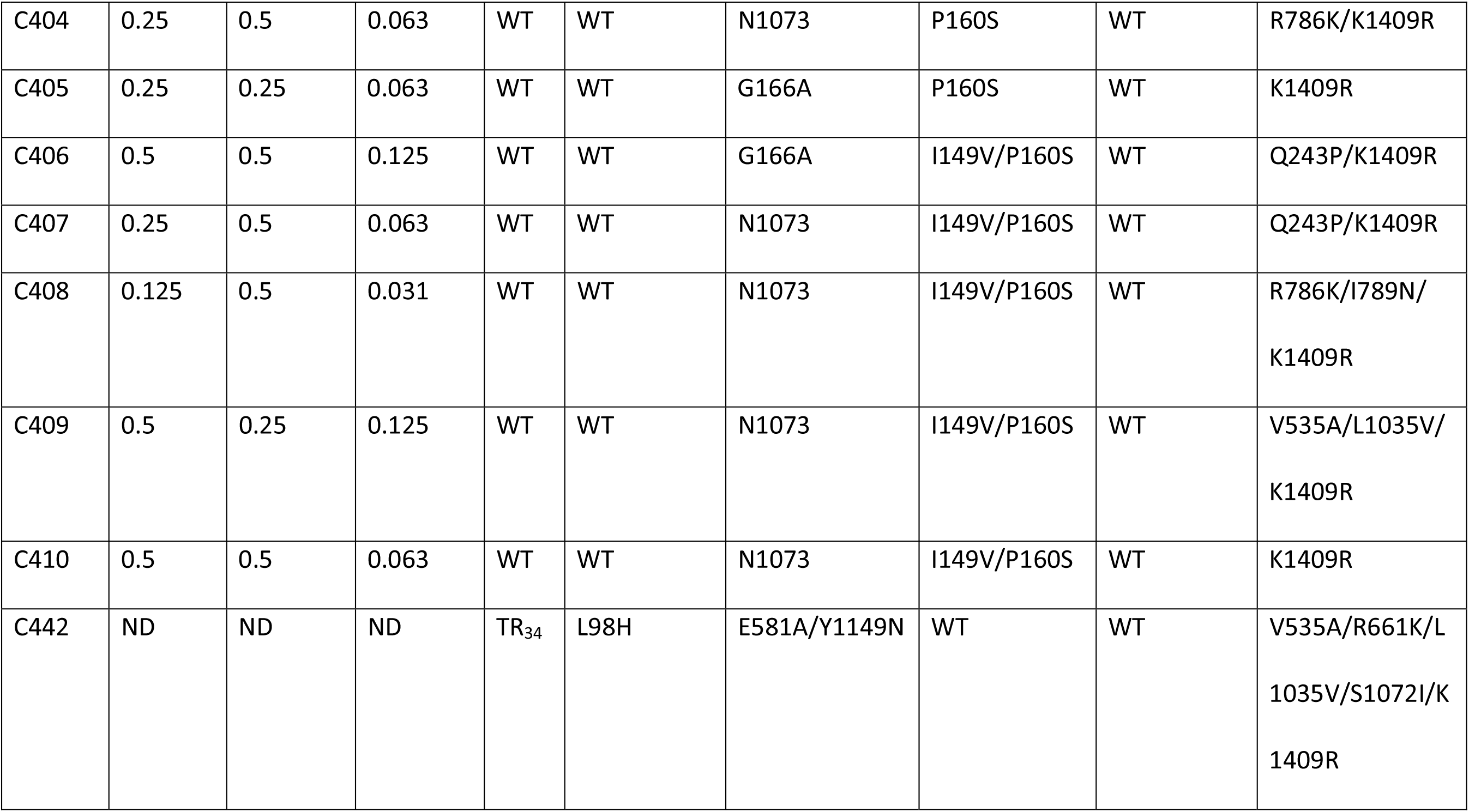
Azole drug susceptibility of CAPA, clinical and environmental *A. fumigatus* isolates grown in minimal medium and the associated candidate polymorphisms. ITR, Itraconazole (clinical breakpoint > 1 mg/L) (58). VOR, Voriconazole (clinical breakpoint > 1 mg/L) (58). POS, Posaconazole (clinical breakpoint > 0.25 mg/L) (58). TEB, Tebuconazole (breakpoint > 4 mg/L). ND, Test not done. C1-C200, E9-E206 and U1-U3 susceptibility testing have previously been reported by Rhodes *et al.* (47). C444 had susceptibility testing done by Mohamed *et al.* (102). C438 and C441 were tested using the Clinical and Laboratory Standards Institute (CLSI) method. C307, C323, C360, C372 and C376 have not been reported before but were conducted by Scourfield (106) and Armstrong-James (105).

**Supp Info Table 5.**
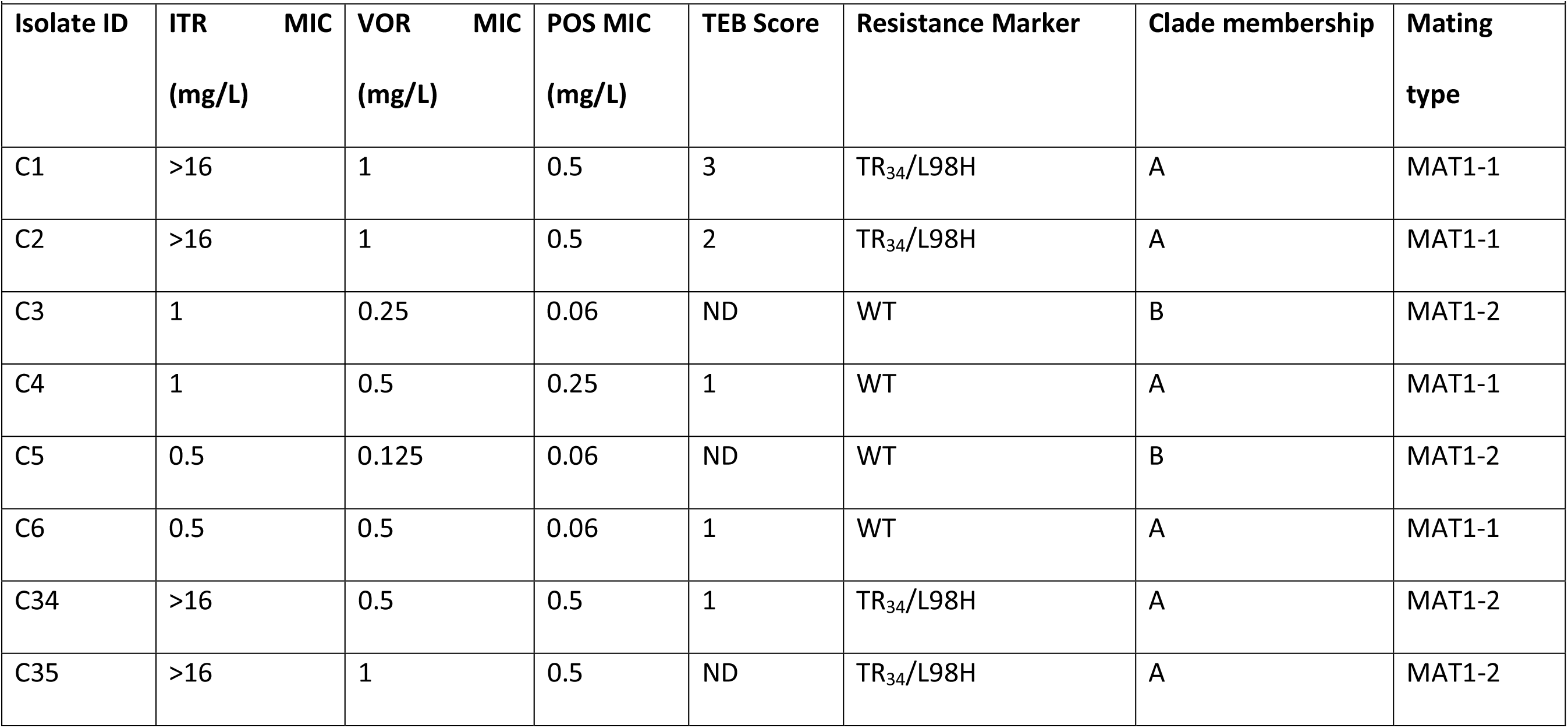
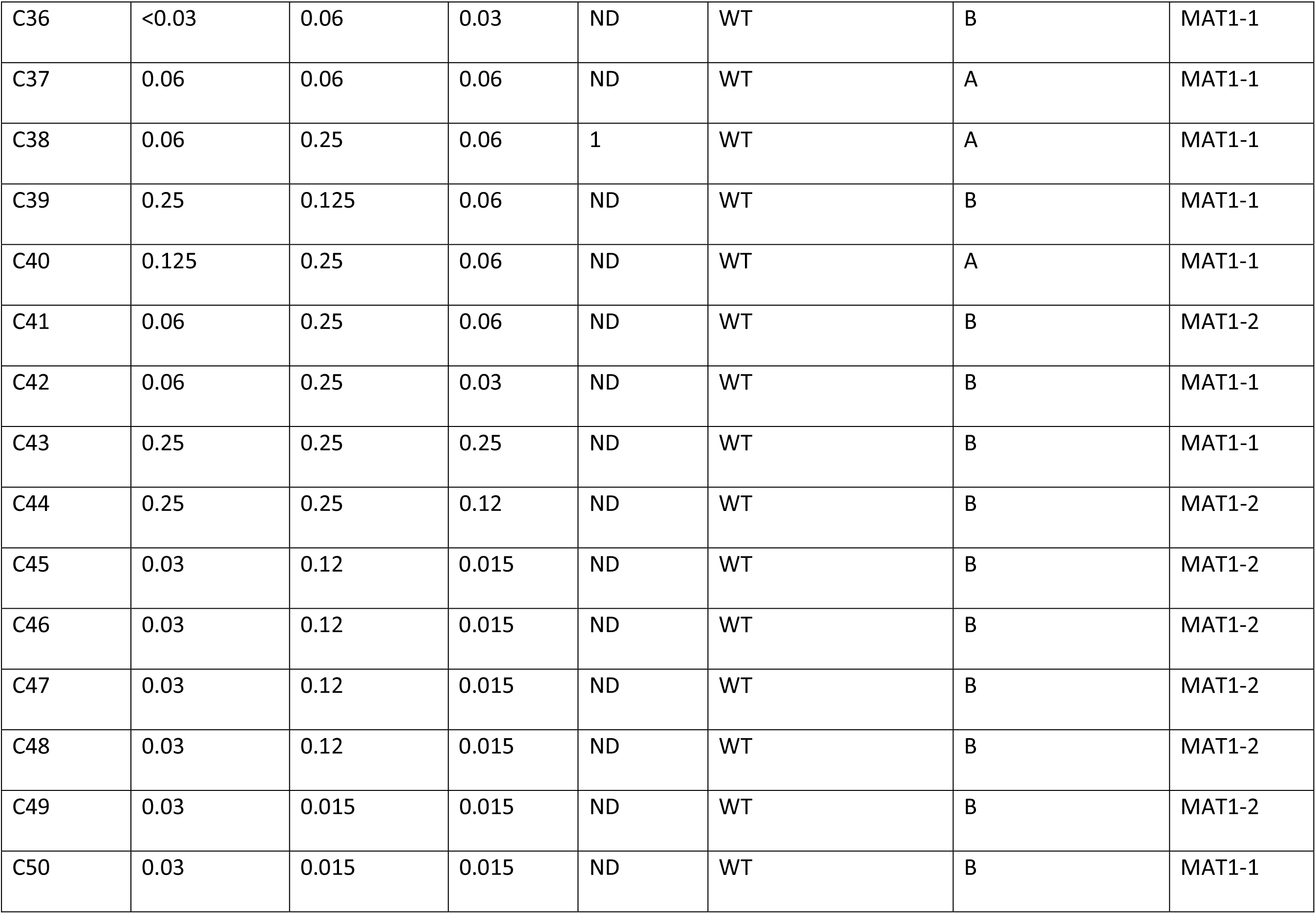
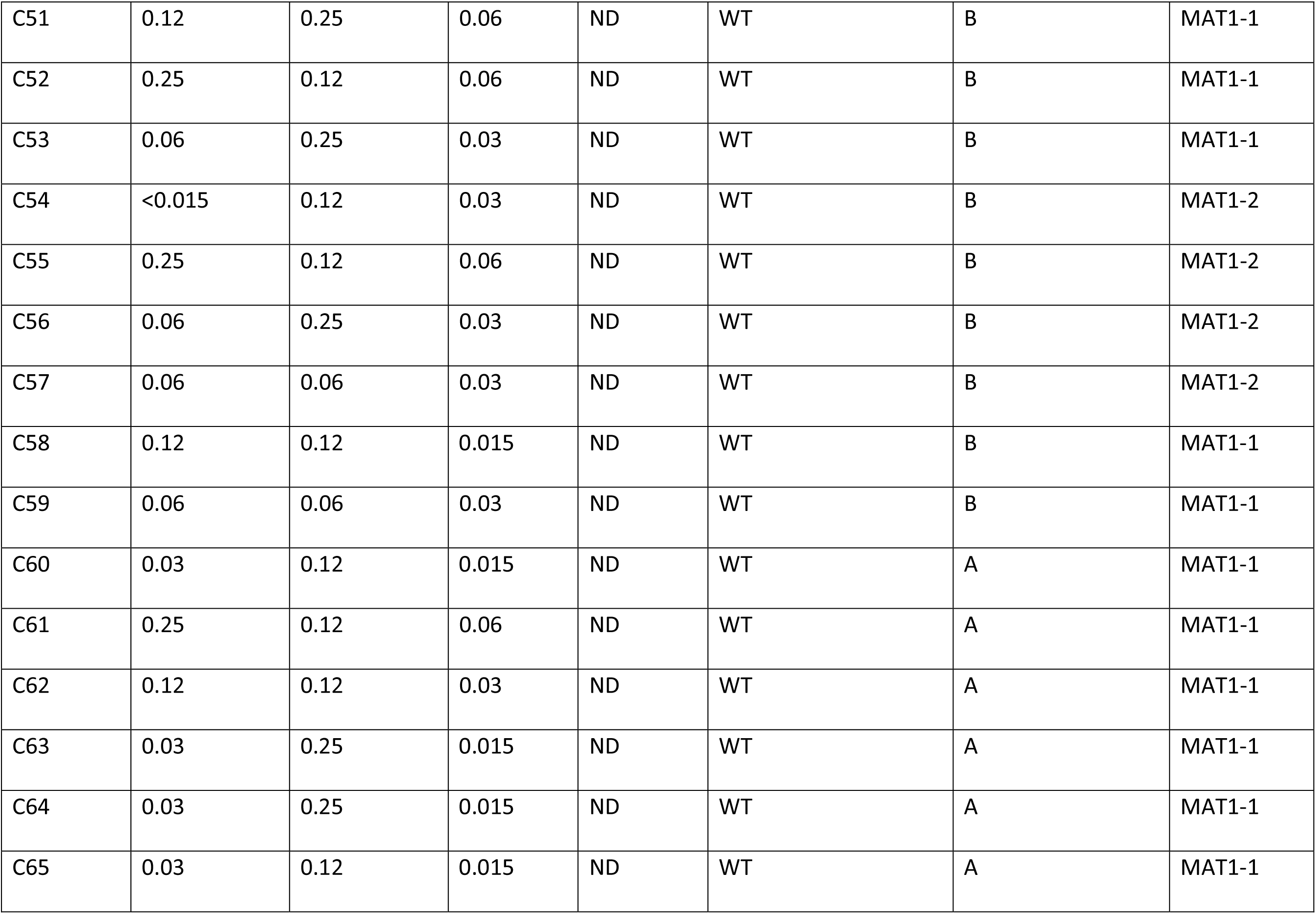
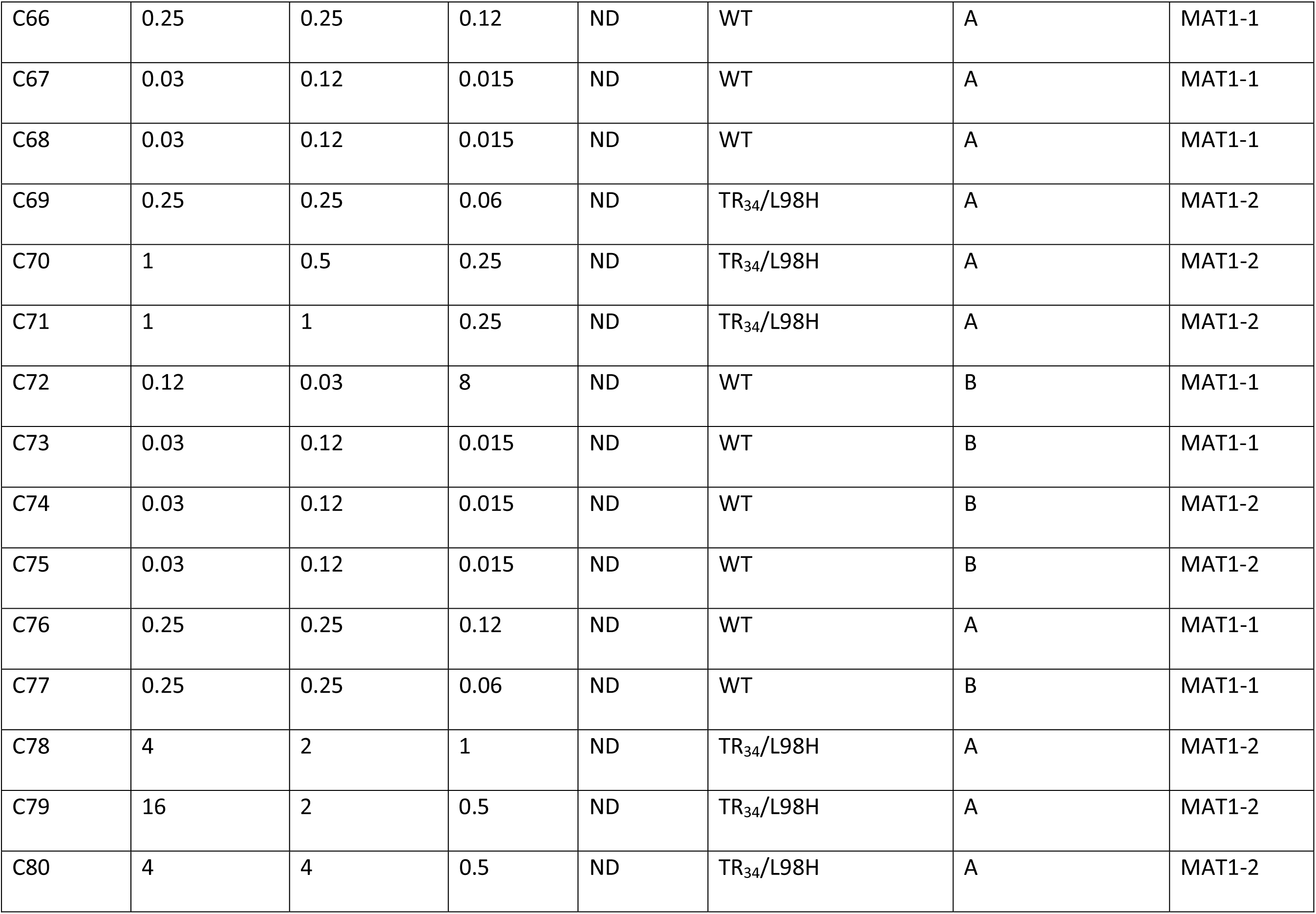
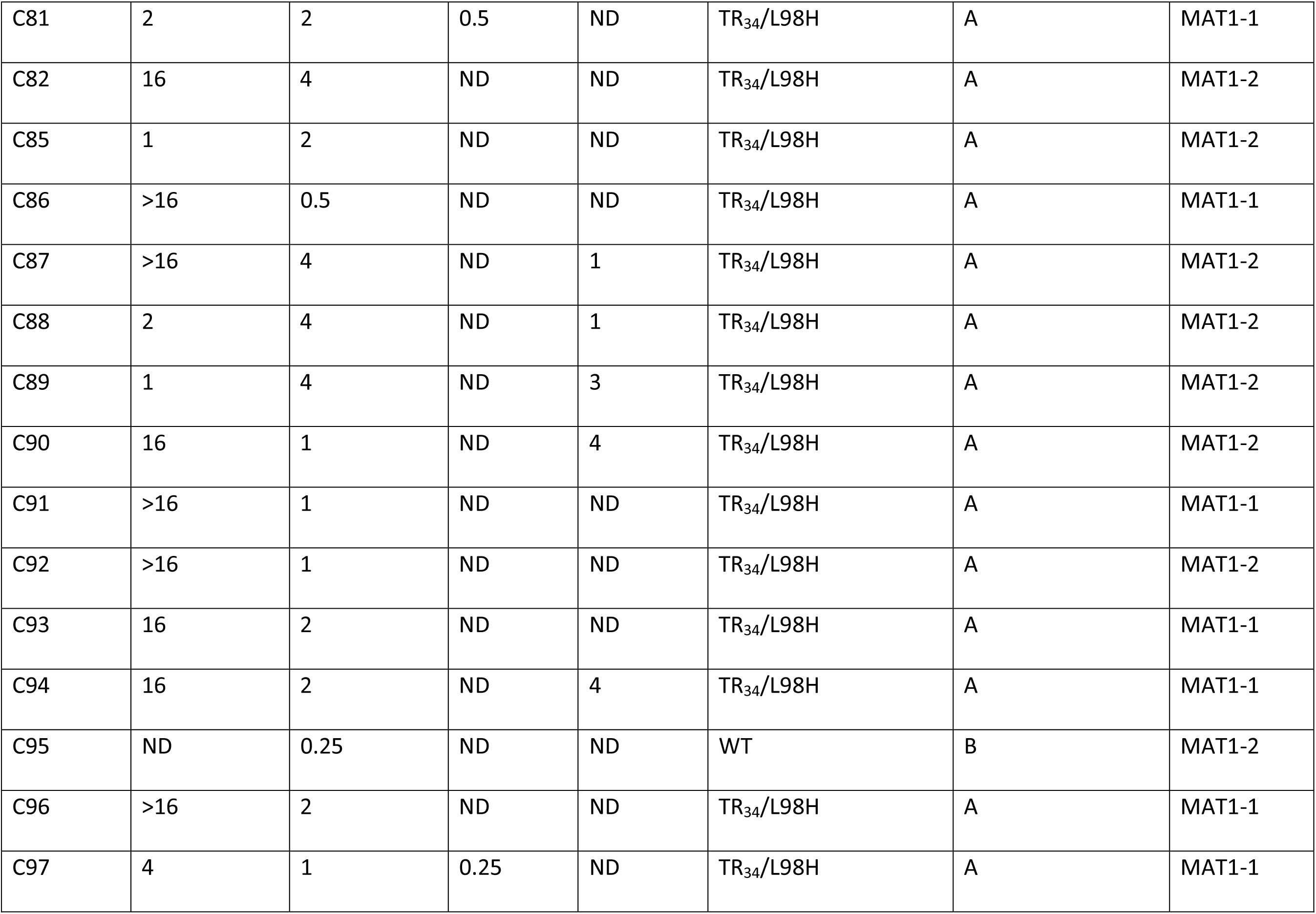
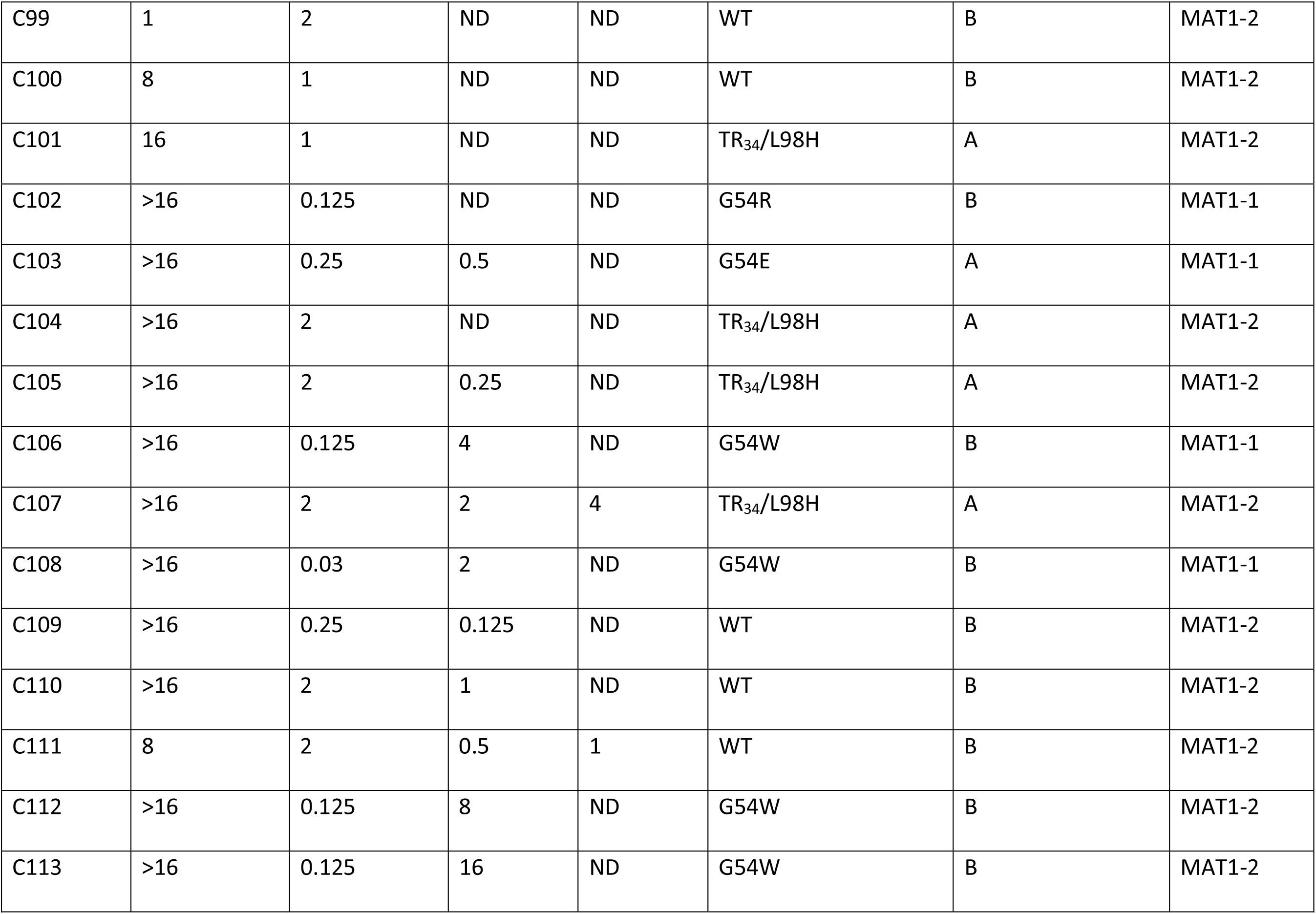
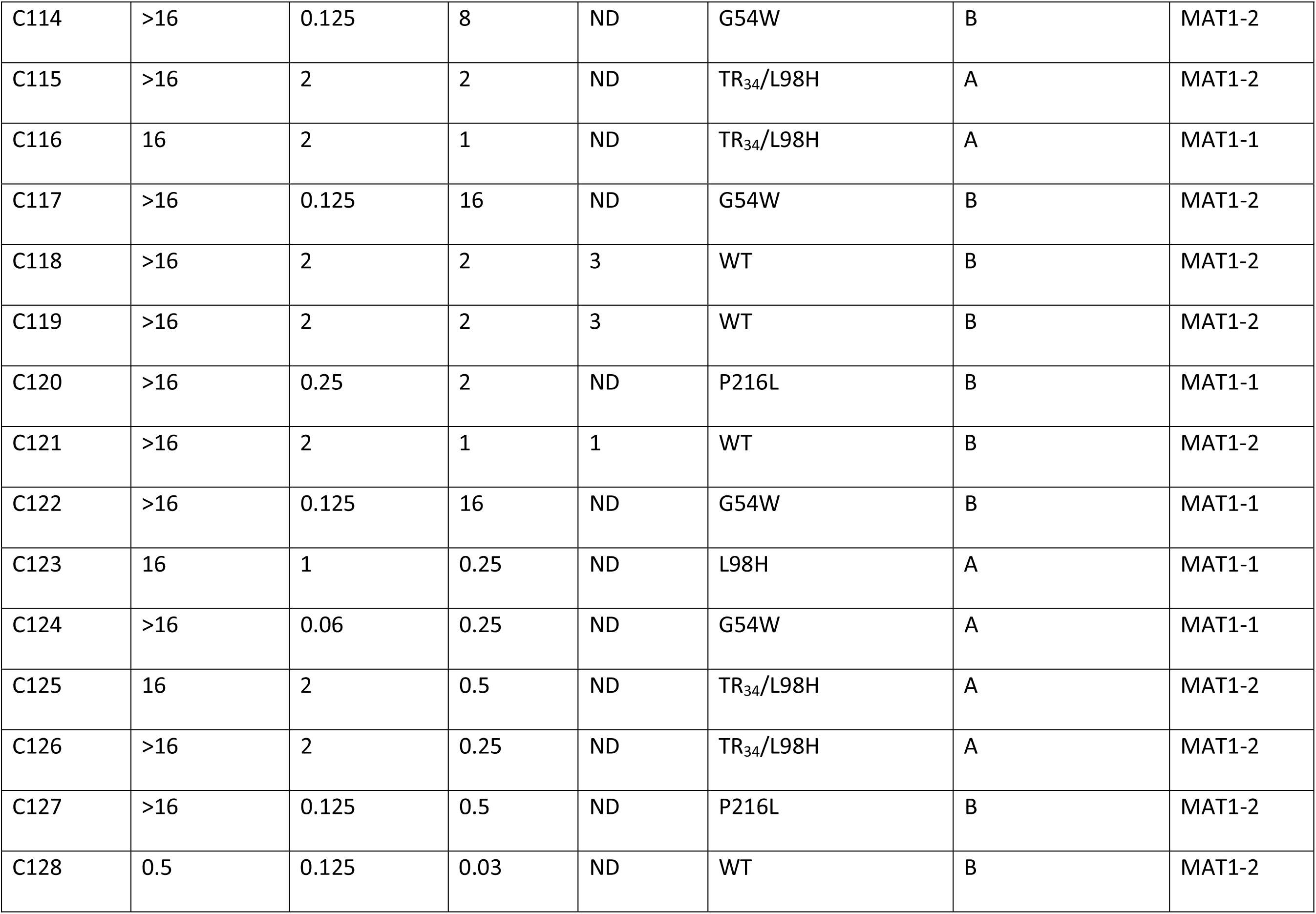
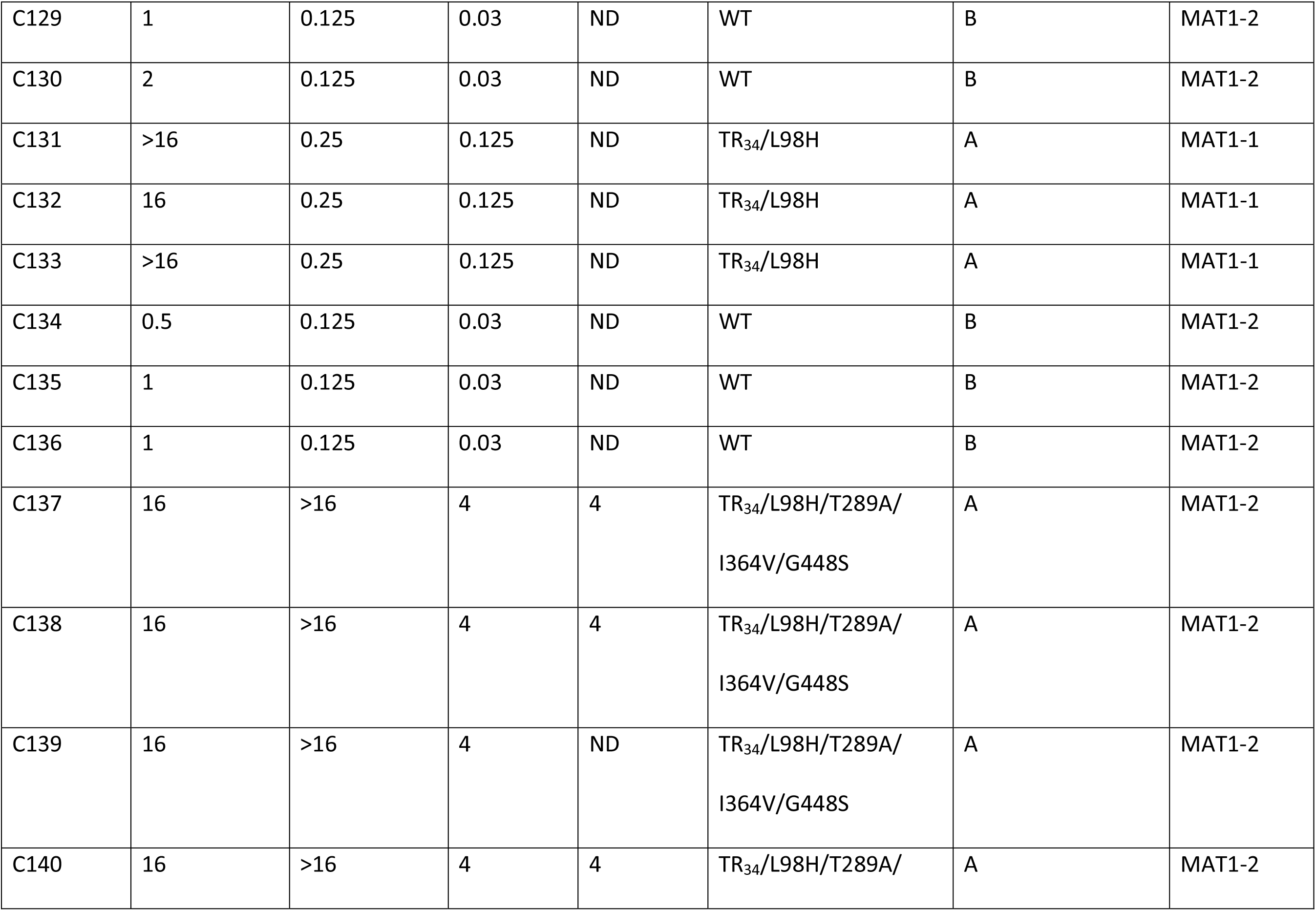
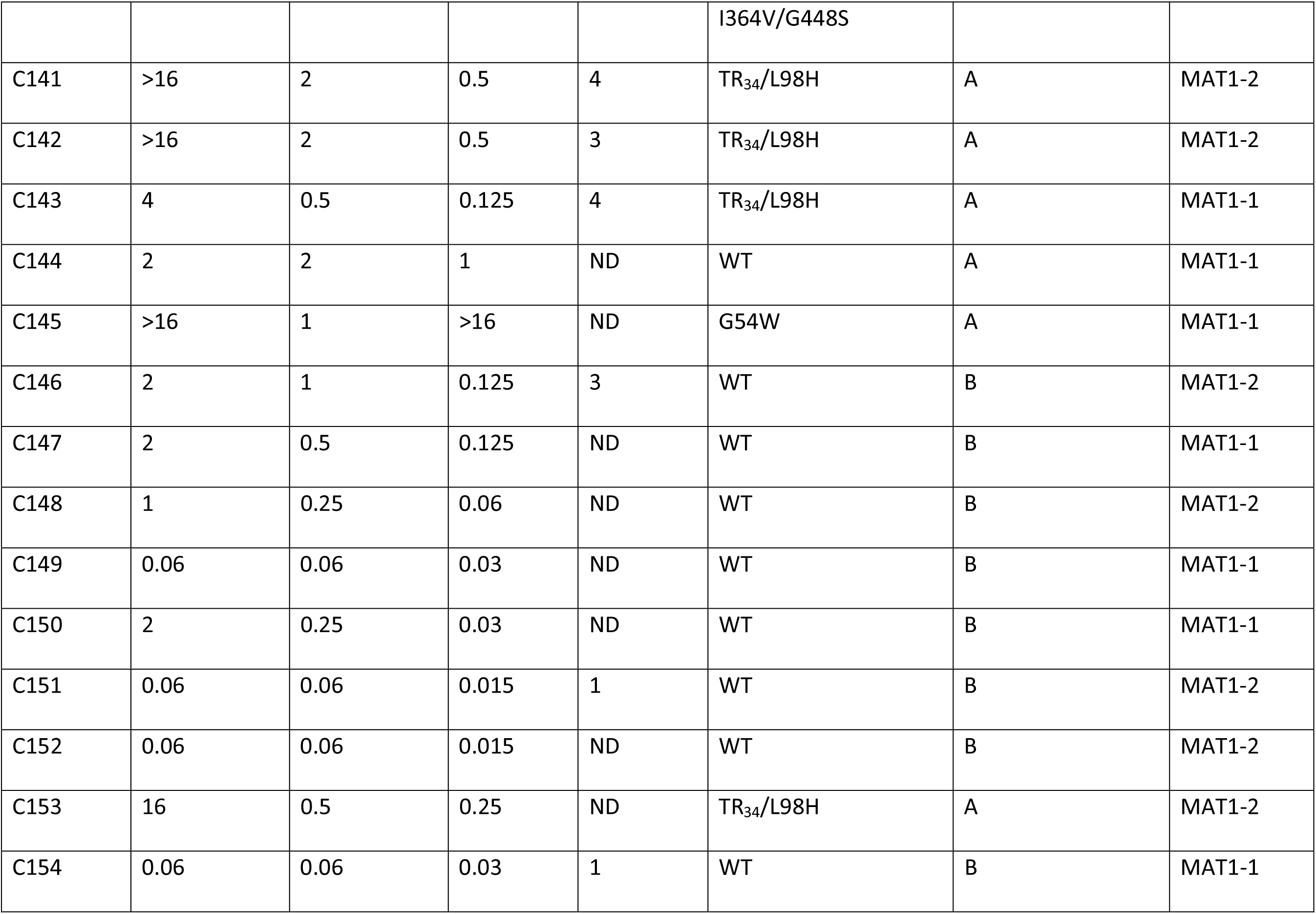
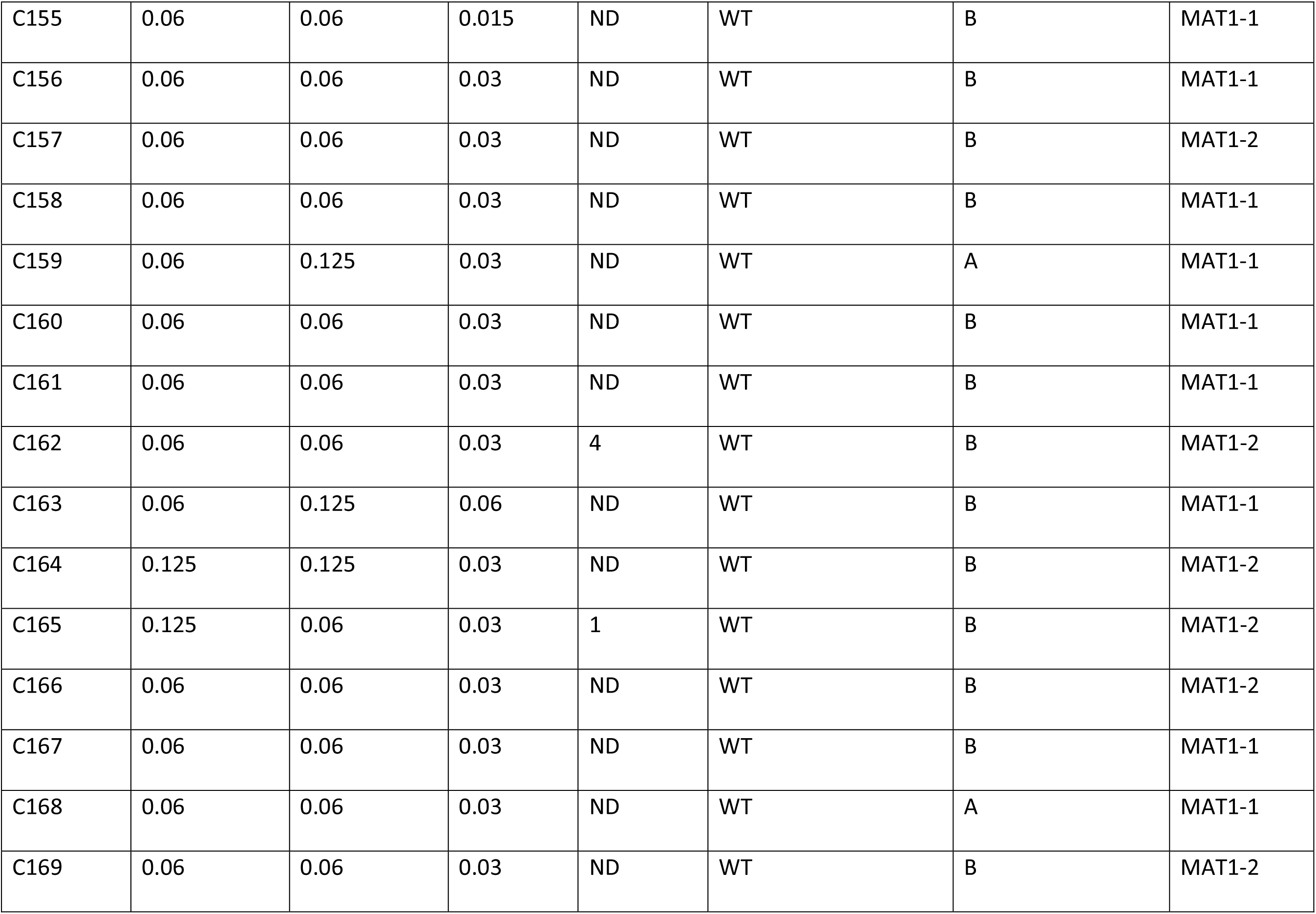
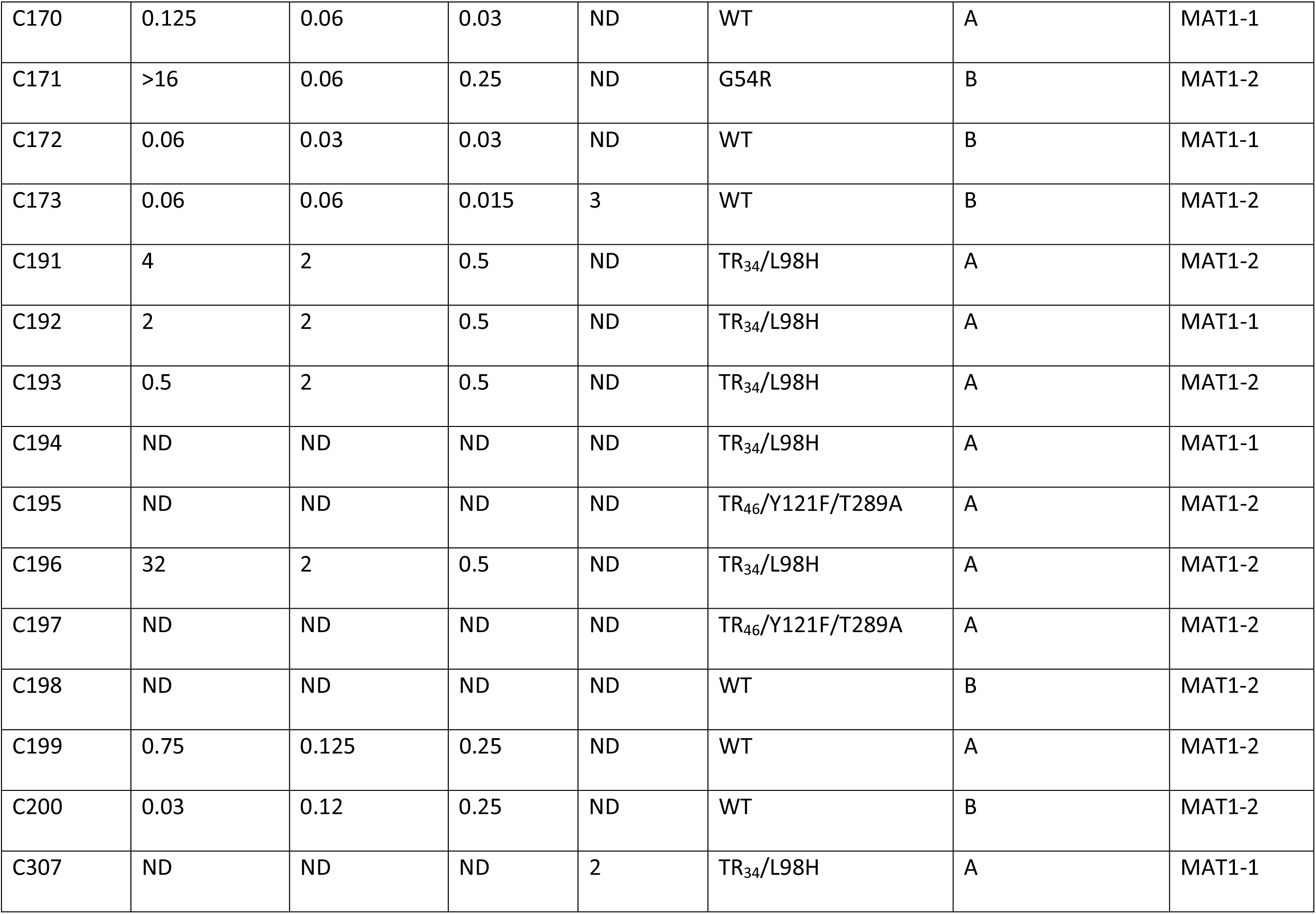
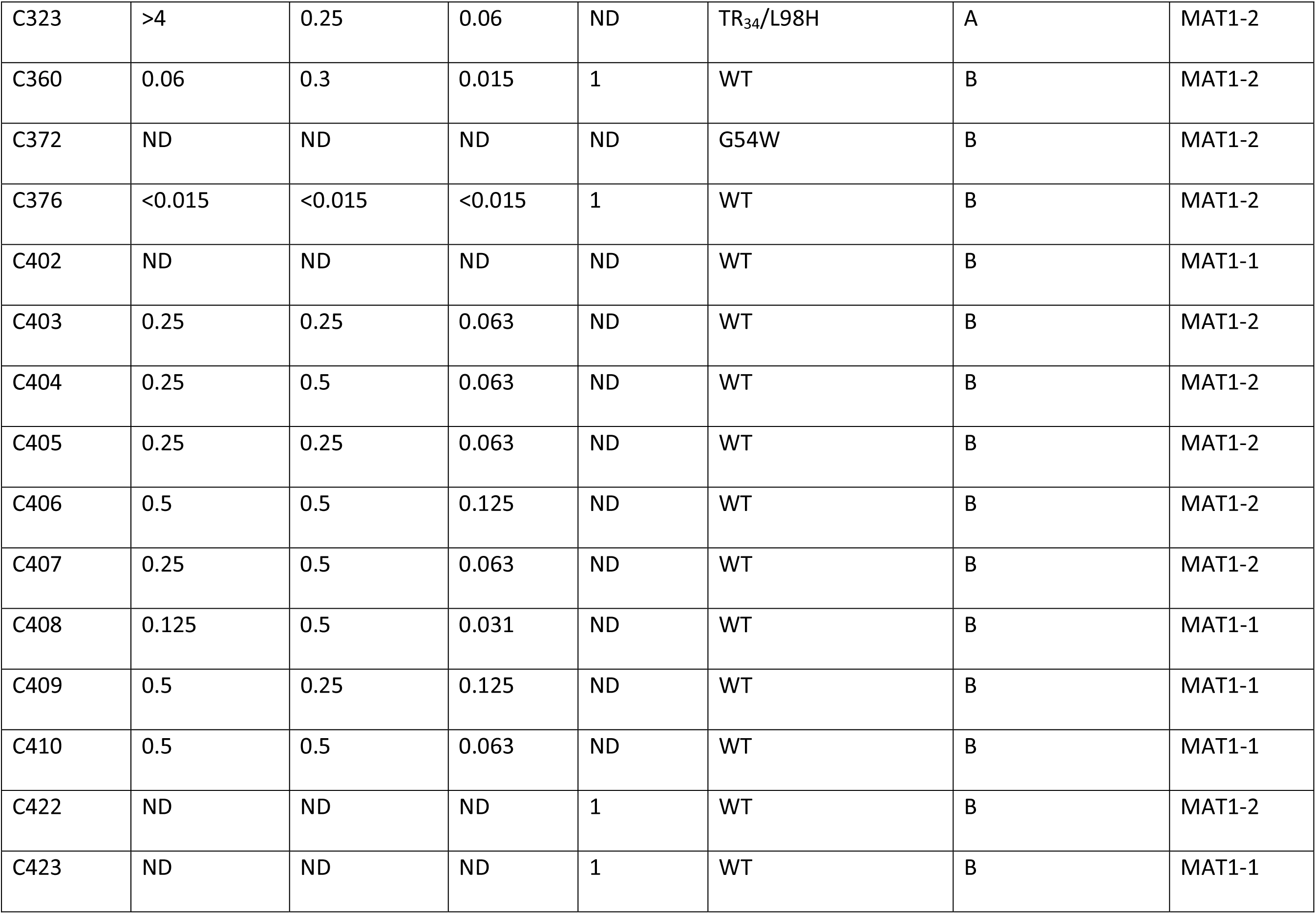
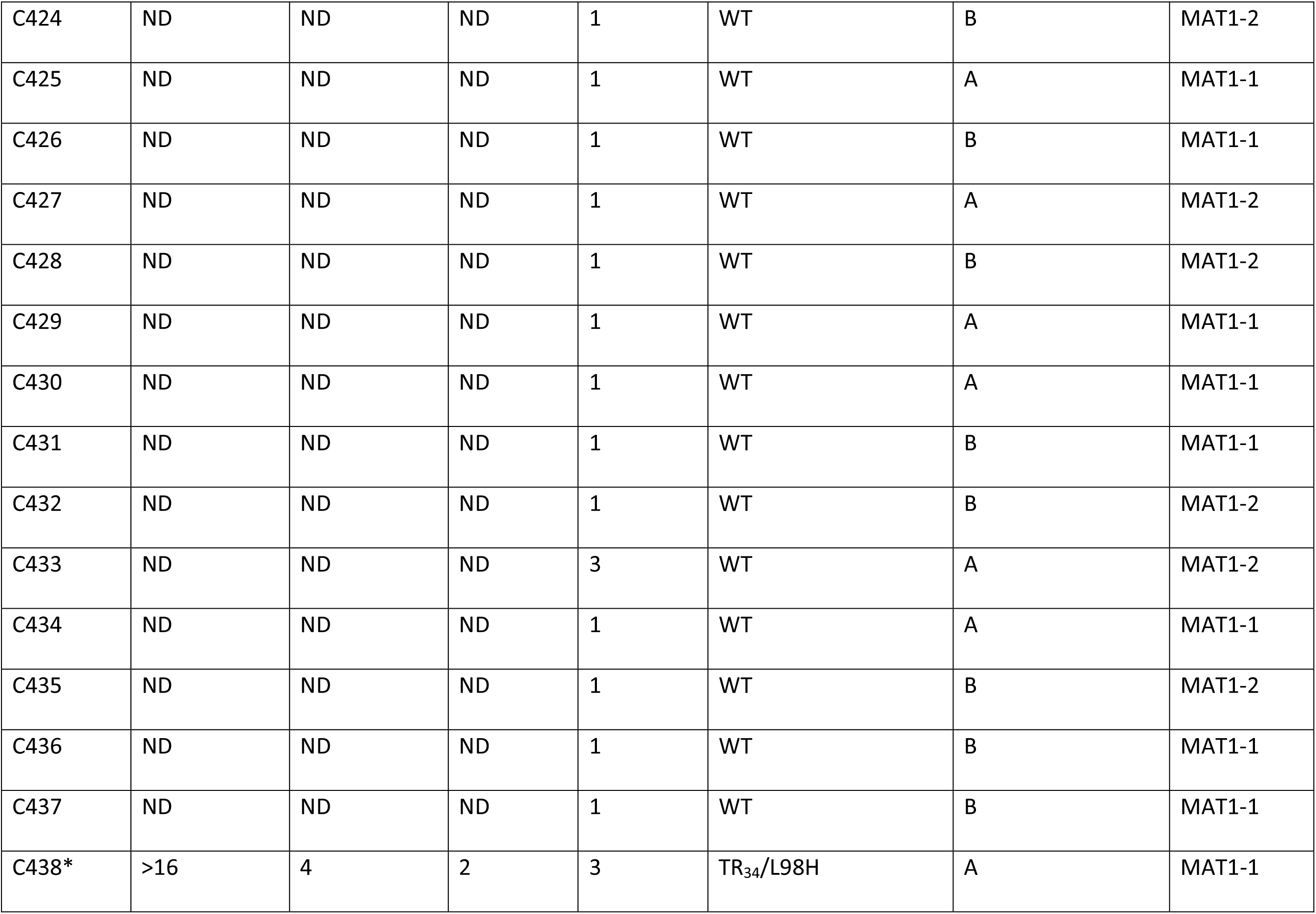
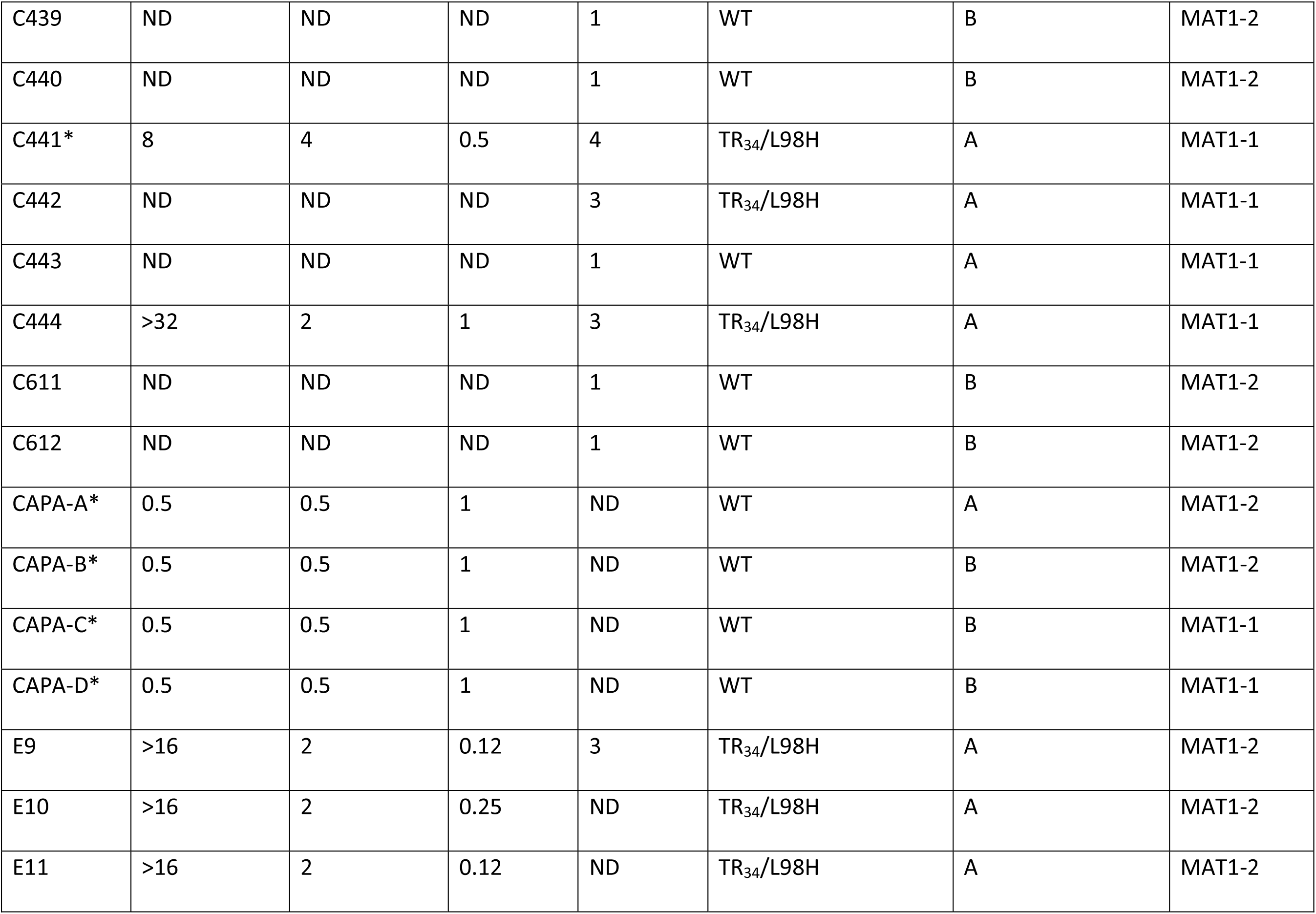
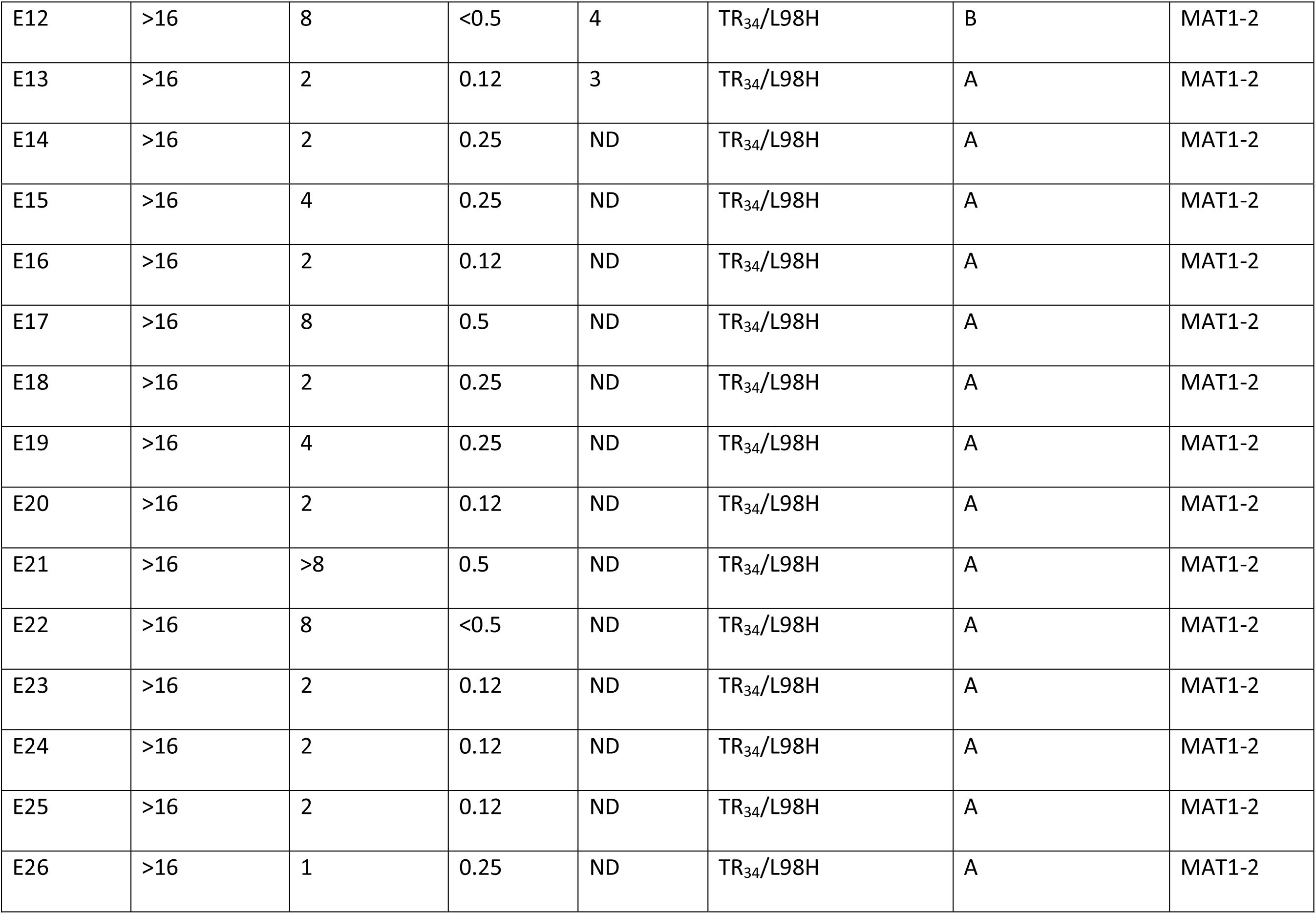
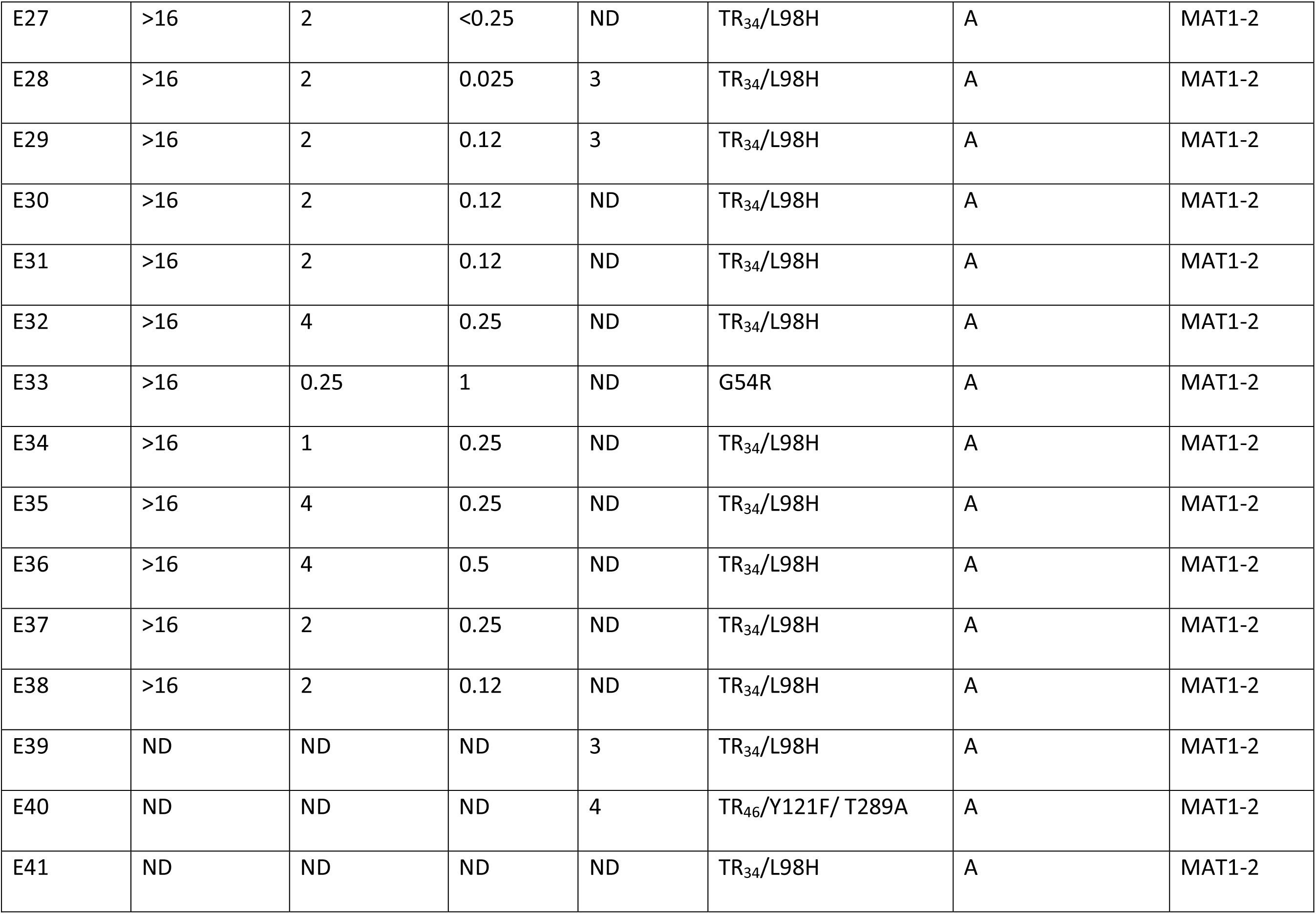
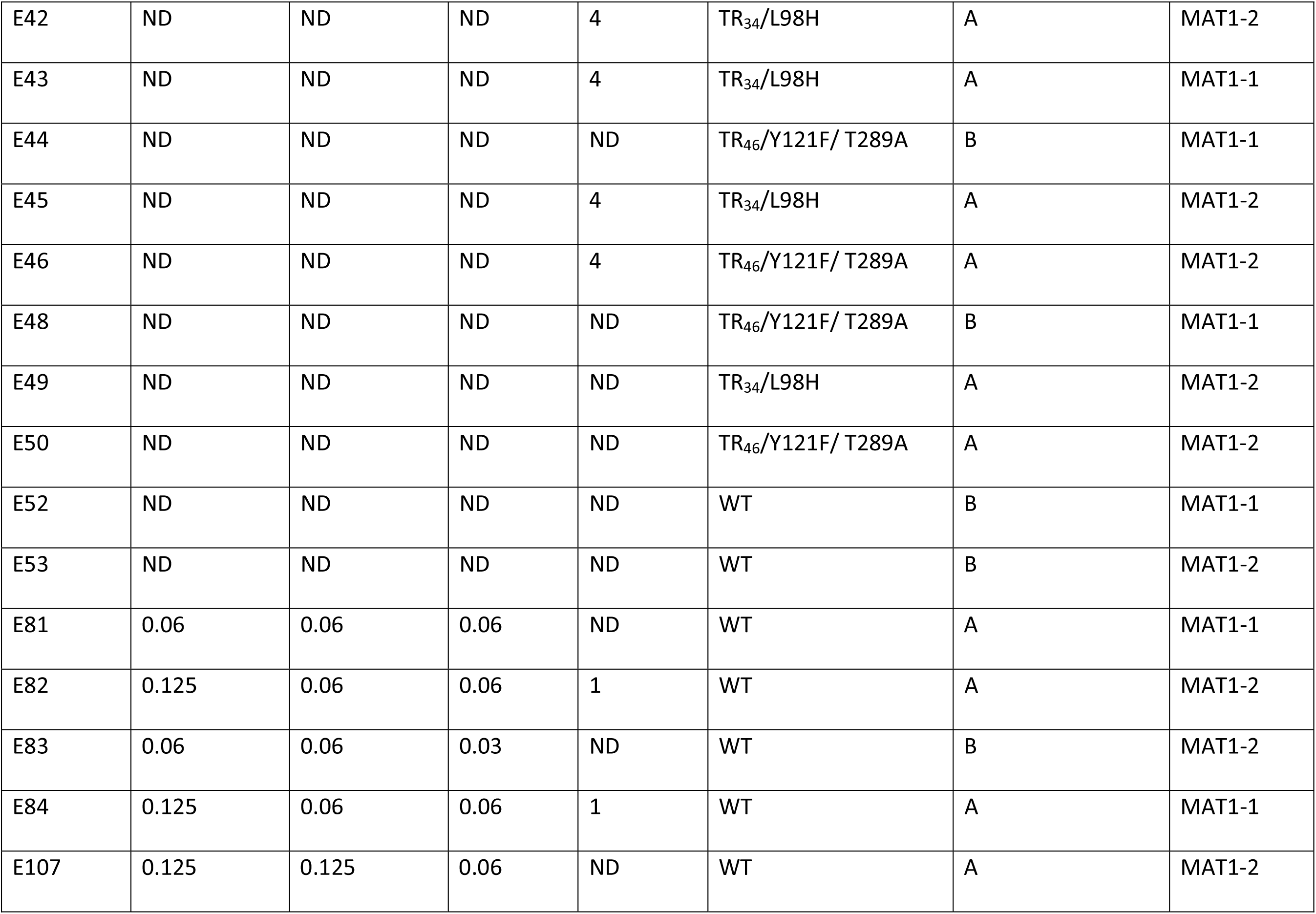
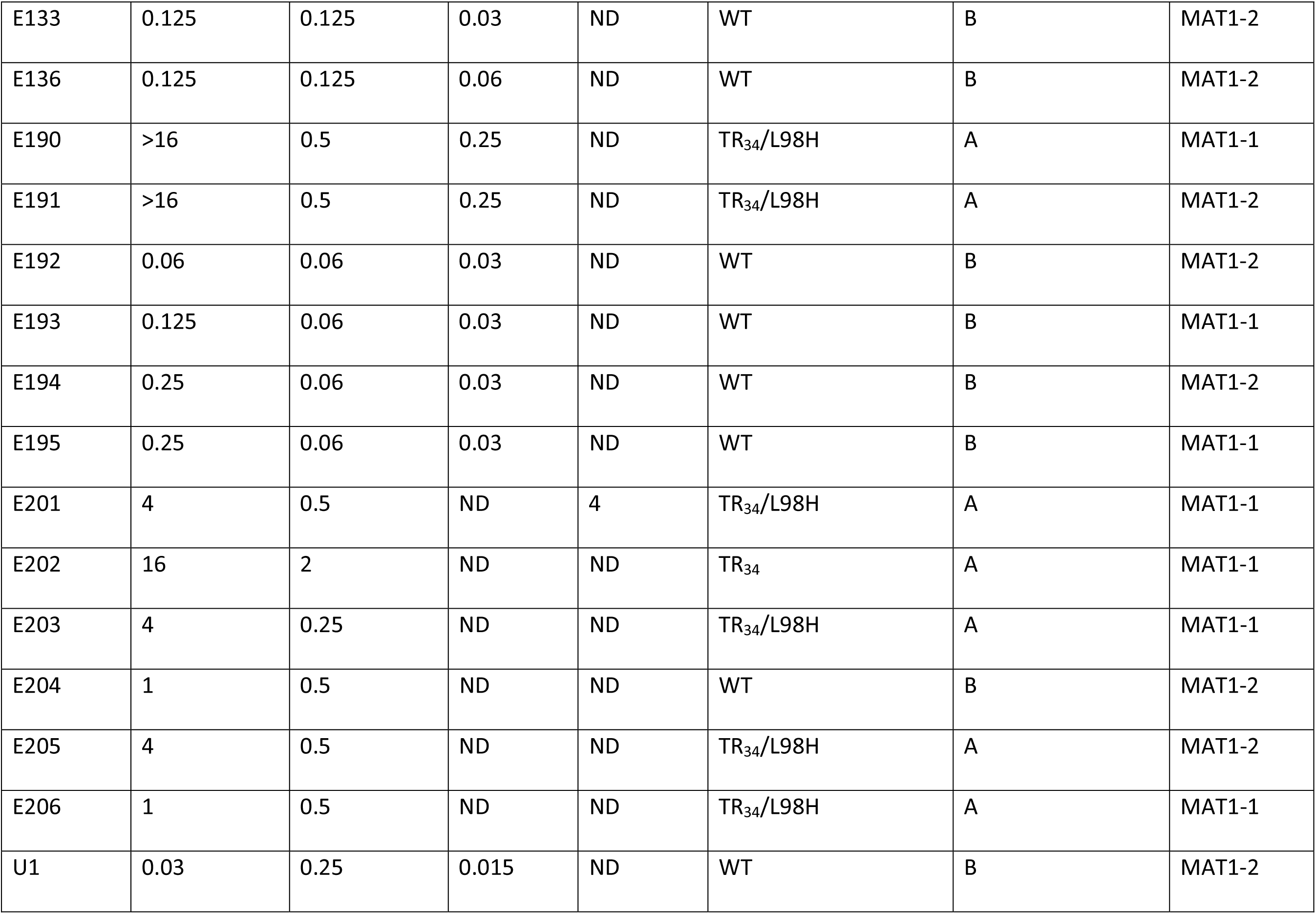
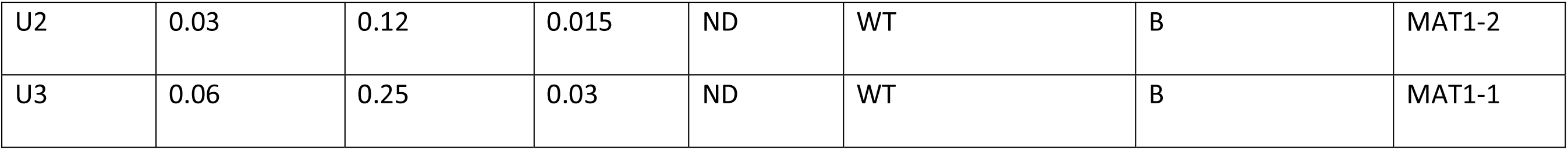
Azole drug susceptibility of *A. fumigatus* isolates used in study grown in minimal medium and the associated polymorphisms. ITR, Itraconazole (clinical breakpoint > 1 mg/L) (58). VOR, Voriconazole (clinical breakpoint > 1 mg/L) (58). POS, Posaconazole (clinical breakpoint > 0.25 mg/L) (58). TEB, Tebuconazole (breakpoint > 4 mg/L). ND, Test not done. C1-C200, E9-E206 and U1-U3 susceptibility testing have previously been reported by Rhodes *et al.* (47). C444 had susceptibility testing done by Mohamed *et al.* (102). CAPA-A – D, C438 and C441 were tested using the CLSI method. C307, C323, C360, C372 and C376 have not been reported before but were conducted by Scourfield (106) and Armstrong-James (105).

---

**Supp Info Figure 1:**
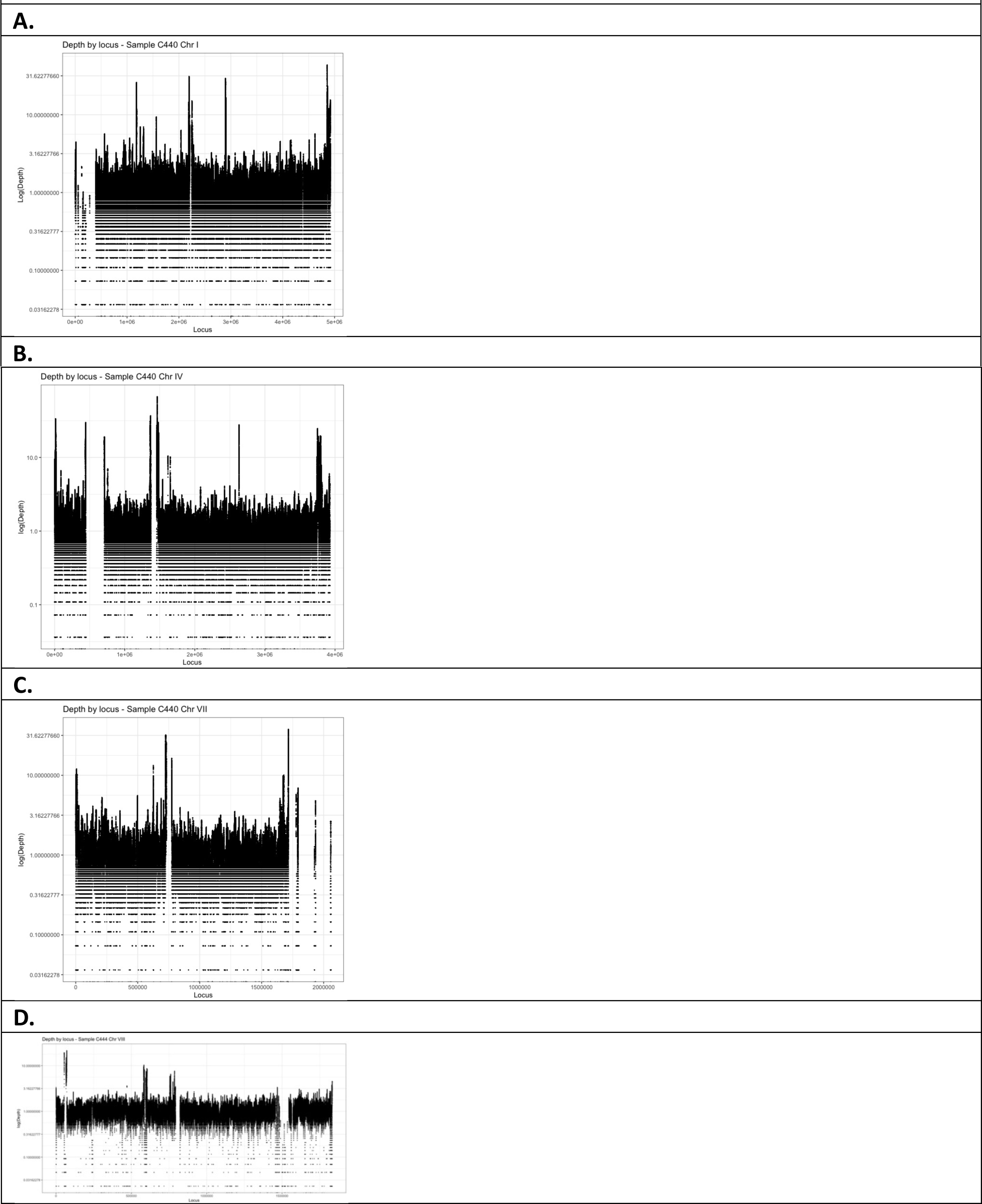
Normalised depth of coverage by locus indicative of duplications and deletion events. **A**. C440 chromosome 1. The deleted region is also found in C408 and C439-C441 **B**. Two regions, a 500 kbp deletion identified in all isolates, and a 1300 kbp deletion around the centromere of chromosome 4. **C**. Two deletions in chromosome 7, relating to the centromere and a region of 1600 kbp found in CAPA-A, CAPA-B, C403, 408, C436 and C439-C441, 444. **D**. Small deletion of 1400 kbp found in chromosome 8 in isolates CAPA-B, CAPA-C, C424, C436, C441, 444.

**Supp Info Figure 2:**
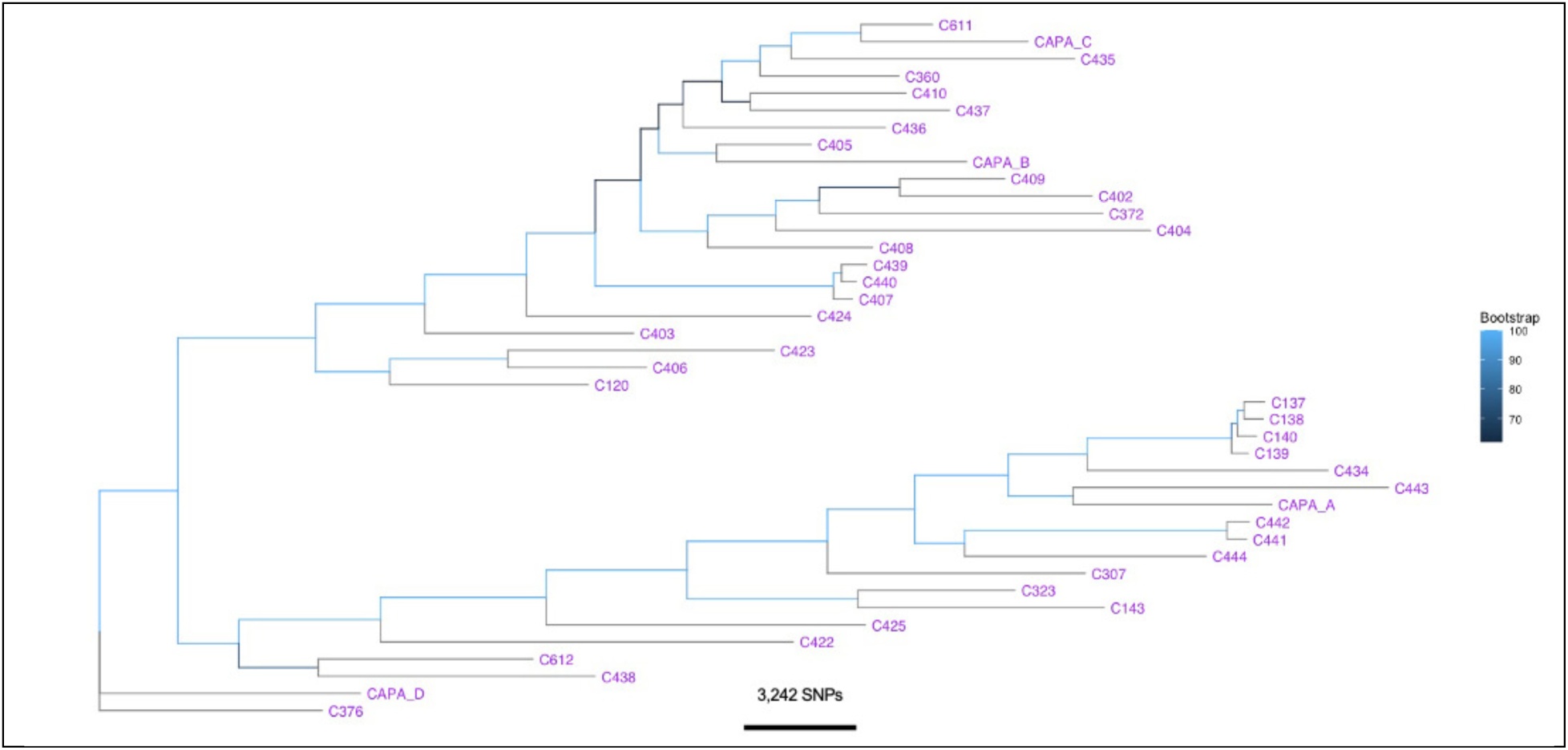
Phylogenetic tree of CAPA isolates, *A. fumigatus* IA control isolates from London, UK, and isolates from colonising COVID-19 patients (Netherlands and Ireland). Rooted ML phylogenetic tree with bootstraps support over 1000 replicates performed on WGS SNP data. Branch length represents the average number of SNPs.

**Supp Info Figure 3:**
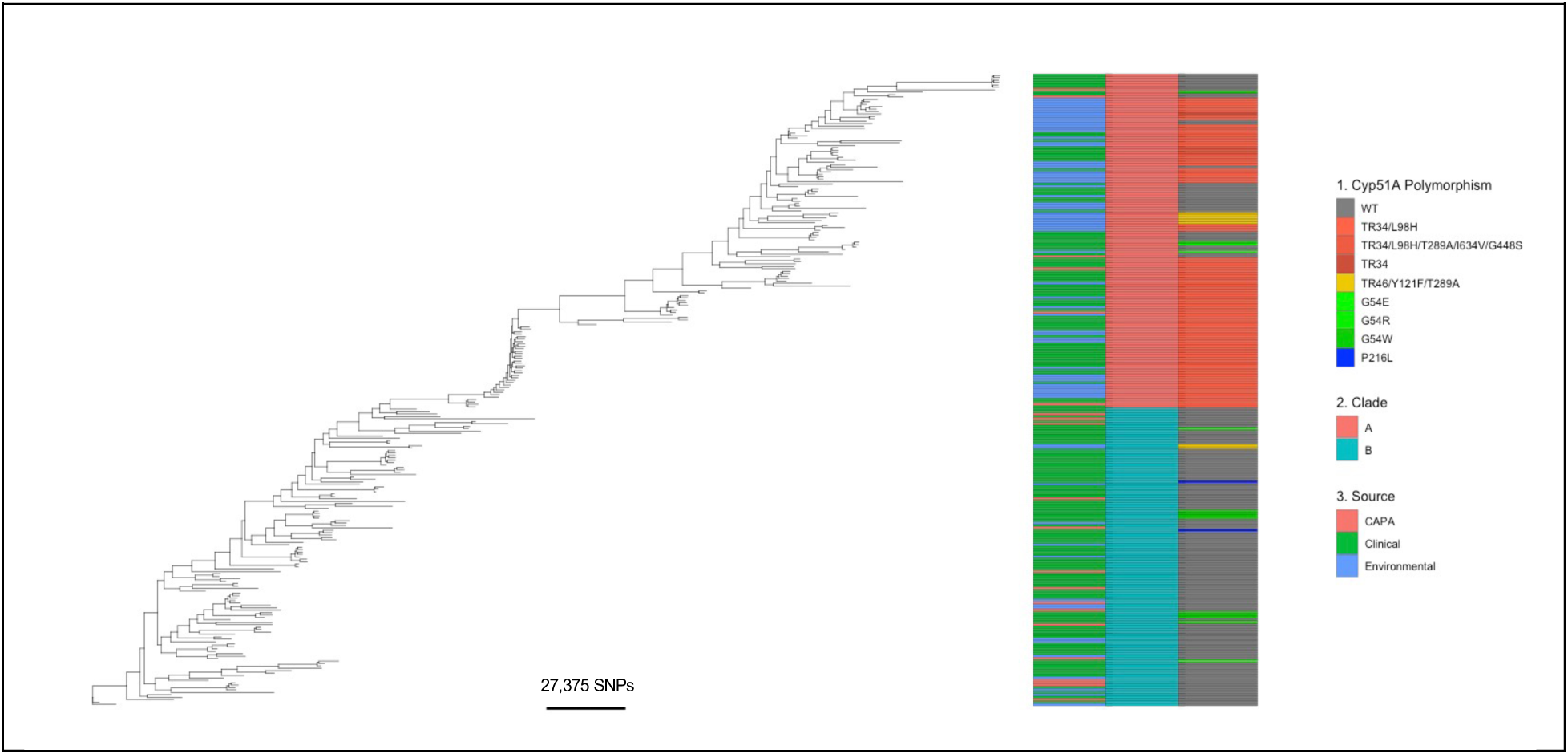
Phylogenetic tree of CAPA and *A. fumigatus* clinical and environmental isolates from Ireland and the UK. Rooted ML phylogenetic tree, showing: Right track 1., if the isolate contains *cyp51A* polymorphism; Middle track 2., the clade the isolate is located; and Left track 3., the source of the isolate (CAPA, Clinical or Environmental). Branch length represents the average number of SNPs.

**Supp Info Figure 4:**
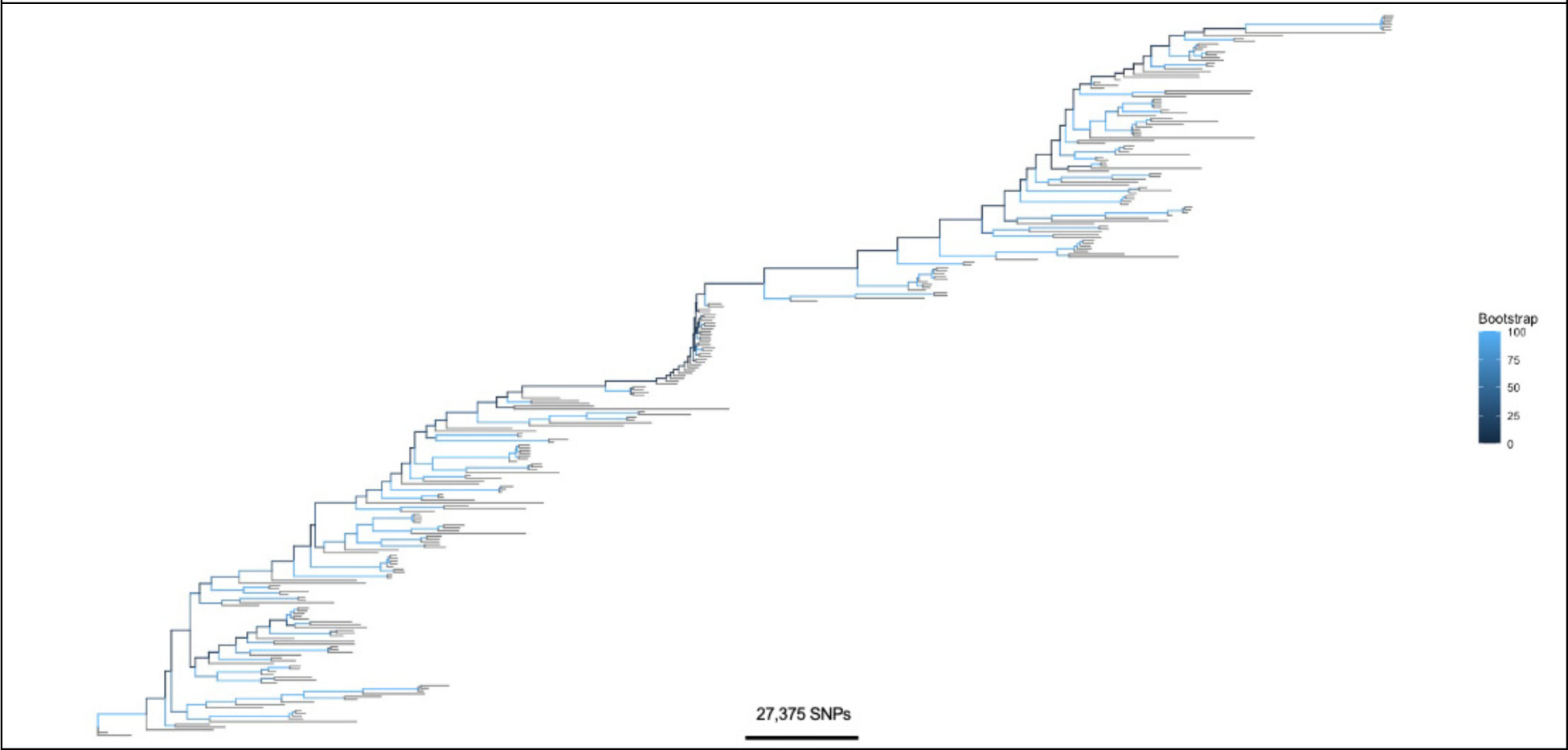
Phylogenetic tree of CAPA and *A. fumigatus* clinical and environmental isolates from Ireland and the UK. Rooted ML phylogenetic tree with bootstraps support over 1000 replicates performed on WGS SNP data. Branch length represents the average number of SNPs.

